# Enhanced Sampling Enables Binding Site Water Reorganization in Polarizable Absolute Binding Free Energy Calculations

**DOI:** 10.1101/2025.10.17.683050

**Authors:** Marharyta Blazhynska, Narjes Ansari, Louis Lagardère, Jean-Philip Piquemal

## Abstract

Water molecules play a critical role in mediating protein–ligand interactions by forming hydrogen-bonding networks and contributing to ligand solvation. However, their behavior—ranging from rapid solvent exchange to persistent occupancy of buried sites—makes accurate binding free-energy estimation with molecular dynamics challenging. Inadequate sampling of water reorganization can bias computed affinities and obscure key interactions by preventing representative hydration states during calculations. To address this, we employ the polarizable AMOEBA force field together with Lambda-ABF-OPES, an integrated enhanced-sampling framework combining lambda-dynamics, multiple-walker adaptive biasing force, and on-the-fly probability enhanced sampling. AMOEBA provides a polarizable description of protein–ligand, ligand–water, and protein–water interactions, while the dynamic alchemical coordinate and adaptive biasing facilitate exploration of coupled ligand–pocket–solvent configurations during ligand decoupling. This construction allows water exchange, binding-site rehydration and solvent reorganization to emerge along the alchemical pathway without explicitly biasing water-based collective variables. Applied to five protein–ligand complexes spanning buried and semi-buried environments, the approach yields binding affinities in good agreement with experiment and captures relevant water-mediated reorganization to provide reproducible absolute binding free energies.

## Introduction

The structural and dynamic properties of water are fundamental determinants of molecular recognition and protein–ligand binding specificity.^1–4^ At the binding interface, water molecules can bridge protein and ligand interactions, potentially modulating both their affinity and selectivity. Structural studies reveal that some crystallographically resolved interfacial waters may form persistent protein–water–ligand hydrogen-bonding networks that reinforce recognition, whereas others exchange rapidly with bulk solvent, dynamically contributing to the specificity and strength of binding.^2,3,5,6^ Such behavior presents substantial complexity for the accurate evaluation of absolute binding free energies, particularly due to the difficulties in adequately sampling the associated solvation process.^5,7–10^ From a thermodynamic perspective, the relevant contribution is determined by the statistical populations of coupled protein–ligand–solvent configurations, rather than by the continuous residence of a particular water molecule at a crystallographic site.

To account for hydration effects, several Monte Carlo (MC)-based water placement and rehydration approaches have been developed.^11–14^ Grand-canonical MC and related insertion/deletion schemes provide an efficient means of sampling water occupancy in buried or semi-buried cavities and have been widely used to identify conserved, displaceable, or energetically unfavorable hydration sites, including those found in bromodomain binding pockets.^15–17^ Their effectiveness nevertheless depends on the protein and ligand conformational ensemble, the definition of the sampled region, and adequate relaxation of the surrounding binding pocket. When water accessibility is controlled by slow side-chain rearrangements, backbone motions, or cavity opening, insertion and deletion moves alone do not guarantee equilibration of these coupled degrees of freedom. Hydration and conformational sampling may therefore remain mutually dependent, motivating approaches in which solvent reorganization is sampled concurrently with ligand and pocket dynamics.

A distinct class of approaches estimates hydration contributions from end-point molecular-dynamics ensembles rather than through explicit water insertion and deletion. A widely employed molecular-dynamics (MD)-based class of approaches for estimating binding affinities comprises the molecular mechanics/Poisson–Boltzmann surface area and molecular mechanics/generalized Born surface area (MM-PBSA/MM-GBSA) methods, which combine molecular mechanics (MM) energies, solvation free energies, and entropic contributions derived from MD simulations.^18,19^ In these approaches, the solvation component approximates the free energy associated with transferring the isolated ligand, the protein, and their complex from vacuum to solvent. Several related strategies have been developed to treat binding-site waters more explicitly. In water-MM-PBSA/MM-GBSA variants, selected explicit waters are retained as part of the receptor, ligand, or complex during end-point free energy evaluation, which can improve the treatment of conserved bridging waters but remains sensitive to the choice and residence of the retained water molecules^20–22^. Despite their computational efficiency and broad applicability, MM-PBSA/MM-GBSA methods do not themselves accelerate transitions among alternative hydration or protein conformational states. Their accuracy can therefore remain sensitive to the representativeness of the underlying ensemble, particularly for complexes involving substantial solvent displacement or conformational change.^23–30^

Hydration thermodynamic mapping methods, including grid inhomogeneous solvation theory (GIST) and WaterMap, provide spatially resolved descriptions of solvent structure and energetics around a receptor or protein–ligand complex.^31–34^ By comparing local water populations and thermodynamic signatures with those of bulk solvent, these approaches can identify strongly localized hydration regions and sites whose displacement may be favorable or unfavorable. Their interpretation nevertheless remains conditional on the sampled protein–ligand ensemble and on the underlying solvent model. Moreover, hydration mapping is not, by itself, equivalent to a continuous alchemical binding free-energy calculation in which ligand, pocket, and solvent reorganize simultaneously along the coupling coordinate. The present work is therefore complementary to these methods. In particular, the *λ*-conditioned hydration distributions and projected free-energy profiles analyzed below characterize solvent response within the sampled alchemical ensemble and are not interpreted as independent absolute water-insertion or water-displacement free energies.

Theory-driven alternatives to these end-point and insertion/deletion strategies employ enhanced-sampling algorithms that explore the free-energy landscape along carefully chosen collective variables (CVs) governing the slow degrees of freedom in protein–ligand complexes.^35^ These methods aim to overcome sampling bottlenecks and provide a more rigorous statistical-mechanical foundation for binding free-energy estimation. Prominent examples include geometrical pathway methods, metadynamics-based approaches, and alchemical free-energy schemes.^29,36–48^ Depending on the chosen formalism, the ligand is either physically separated from the binding site or reversibly decoupled/annihilated through a series of intermediate states. In both cases, the vacated pocket becomes accessible to solvent molecules, and water infiltration can alter critical protein–ligand interactions or hinder the reformation of bridging waters during ligand reassociation. Inadequate sampling of this rehydration process can bias the computed thermodynamics and impede recovery of representative interfacial structures. Adequate sampling of pocket rehydration remains particularly challenging for buried or semi-buried ligands, and has been shown to be crucial for achieving quantitative agreement with experiment in alchemical simulations.^9,49,50^

Among the methods developed to address solvent-sampling bottlenecks directly, water-based collective variables have emerged as powerful extensions of enhanced-sampling approaches.^51–53^ Such variables may describe local water occupancy, density, coordination, or hydrogen-bonding patterns and can be biased to promote transitions between dry and hydrated binding-site states. They are particularly useful when spontaneous solvent entry or displacement is slow on the accessible simulation timescale. Their application, however, requires the identification and validation of system-specific solvent descriptors, and the resulting sampling efficiency depends on whether the selected variables capture the relevant coupling among water, ligand, and protein conformational degrees of freedom.

Despite these developments, it remains unclear whether continuous alchemical sampling can recover the relevant binding-site hydration ensemble without explicitly biasing water occupancy, particularly in buried and semi-buried complexes containing persistent or slowly exchanging hydration motifs. We therefore examine whether solvent exchange and binding-site rehydration can emerge from the coupled response to ligand decoupling when the alchemical coordinate is sampled dynamically using a polarizable force-field model.

To address this question, we employ Lambda-ABF-OPES,^54,55^ an alchemical enhanced-sampling framework built upon continuous *λ*-dynamics.^56,57^ In this formulation, *λ* is treated as a dynamical variable that scales the non-bonded interactions between the ligand and its environment. Repeated exploration of partially coupled and decoupled states reduces the ligand steric and electrostatic footprint, transiently increasing solvent accessibility to the binding pocket. Water entry, exchange, and displacement can therefore occur as part of the coupled response of the ligand–pocket–solvent system, without directly biasing water positions or occupancies.

This mechanism does not imply that water-specific collective variables are unnecessary in every system. Additional orthogonal sampling may remain required when solvent accessibility is controlled by slow protein conformational gating that is only weakly coupled to *λ*. The present work instead tests whether dynamic alchemical sampling is sufficient to recover the relevant solvent response in buried and semi-buried complexes for which hydration is strongly coupled to ligand decoupling.

When the relevant hydration transitions are sufficiently coupled to *λ*, solvent reorganization may be sampled without introducing an explicit water-based CV. By integrating *λ*-dynamics with multiple-walker adaptive biasing force (ABF) and OPES-Explore (OPES*_e_*),^58^ the approach facilitates efficient exploration of relevant configurational space. Specifically, multiple-walker ABF enables independent simulations to share mean-force information, while OPES*_e_* constructs an adaptive bias from the evolving sampled distribution using a well-tempered target.^59–61^ Together, these components enhance sampling along *λ* and may facilitate relaxation of degrees of freedom coupled to the alchemical transformation, without guaranteeing equilibration of slow modes that are only weakly coupled to *λ*.

Importantly, efficient sampling depends on the careful definition of a restraint that confines the accessible phase space when the ligand is partially or fully decoupled, thereby improving convergence.^62,63^

To this end, the Lambda-ABF-OPES^55^ scheme employs a single structural restraint coordinate, the distance-to-bound-configuration (DBC) CV,^55,64^ defined as the root-mean-square deviation (RMSD) of the ligand relative to a moving reference of the protein binding site. A flat-bottom harmonic restraint applied to the DBC defines the bound configurational domain by confining the ligand position, orientation, and conformation while still allowing relevant conformational changes. This strategy confines the ligand to the relevant bound region while allowing native-like fluctuations, thereby improving sampling efficiency and reducing excursions to uninformative decoupled states.

It should be noted that the accuracy of absolute binding free-energy estimates also critically depends on the choice of force field. Fixed-charge force fields do not respond explicitly to changes in the local electrostatic environment, whereas polarizable models^65–67^ include induction effects in the description of solvation and protein–ligand interactions.^68,69^ In particular, polarizable water models allow dynamic adjustment of dipole moments, thereby providing an environment-dependent description of hydration structure and water response.^70–73^ Combining Lambda-ABF-OPES with the polarizable AMOEBA force field^74^ provides an explicit polarizable description of protein–ligand, ligand–water, and protein–water interactions. We apply this framework to five protein-ligand complexes encompassing diverse ligand chemotypes (cyclic and aliphatic), binding affinities (moderate to strong), and pocket topologies (buried and semi-buried, Table 1 and Fig. 1). These complexes span bromodomains, heat shock proteins, and a major urinary protein, covering a range of binding environments where water-mediated and polarization effects are expected to be critical.^9,13,75^ Across these systems, the calculated binding affinities are compared with experiment (Table 2), while conventional and alchemical simulations are analyzed to determine whether structural hydration motifs, water exchange, and pocket resolvation are recovered without explicit water-based CVs. Accordingly, the primary scope of this work is to assess the accuracy and reproducibility of the Lambda-ABF-OPES binding free-energy calculations and to characterize the solvent response sampled along the alchemical pathway.

**Figure 1.**
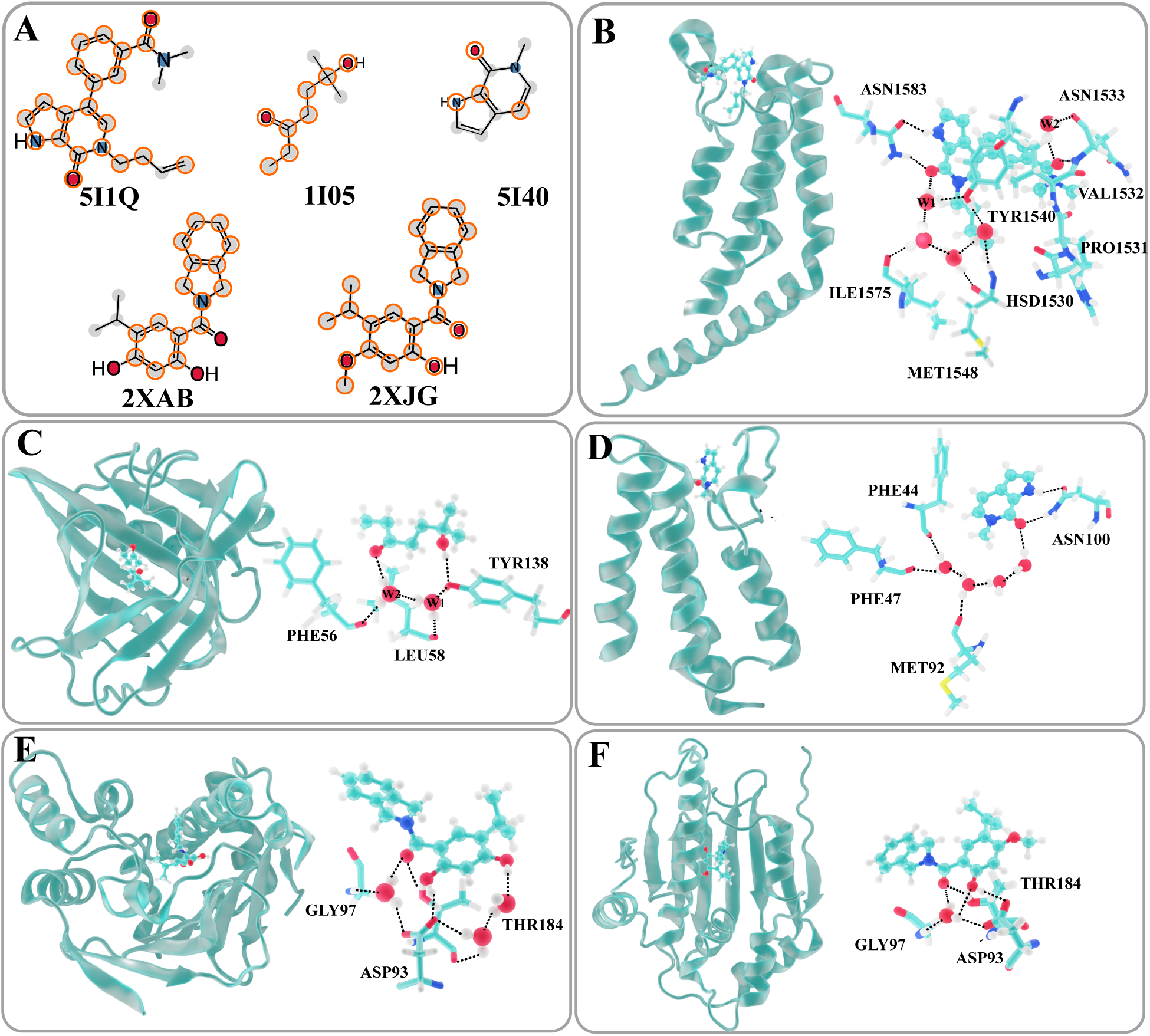
Structural analysis of protein-ligand complexes and their bridging hydration networks. (A) 2D chemical structures of the studied ligands, identified by their PDB accession codes: 5I1Q (TAF1(2) in complex with [6-(but-3-en-1-yl)-7-oxo-6,7-dihydro-1H-pyrrolo[2,3-c]pyridin-4-yl]-N,N-dimethylbenzamide), 1I05 (MUP-I in complex with 6-hydroxy-6-methyl-3-heptanone), 5I40 (Bromo-domain containing protein in complex with 6-methyl-1,6-dihydro-7H-pyrrolo[2,3-c]pyridin-7-one), 2XAB (Heat shock protein HSP90-*α* in complex with (1,3-dihydro-isoindol-2-yl)-(2,4-dihydroxy-5-isopropyl-phenyl)-methanone), and 2XJG (Heat shock protein HSP90-*α* in complex with (1,3-dihydro-isoindol-2-yl)-(2-hydroxy-5-isopropyl-4-methoxy-phenyl)-methanone). Ligand atoms involved in the DBC definition are indicated by orange circles. (B)-(F) Representative 3D views of the 5I1Q, 1I05, 5I40, 2XAB, and 2XJG protein-ligand complexes, respectively. Proteins are rendered in cyan cartoon representation, while ligands are displayed in ball-and-stick models. For each system, buried water molecules (red spheres) and the polar-contact network between water, the ligand, and key amino-acid residues are shown.

**Table 1.** Overview of Studied Protein-Ligand Complexes and Binding Environments.

| PDB id | Protein | Ligand | Resolution (Å) | Complex type* (% Shielded) | Total atoms |
| --- | --- | --- | --- | --- | --- |
| 5I1Q<sup>76</sup> | Second bromodomain of Transcription initiation factor 1 (TAF1(2)) | 3-[6-(but-3-en-1-yl)-7-oxo-6,7-dihydro-1H-pyrrolo[2,3-c]pyridin-4-yl]-N,N-dimethylbenzamide | 1.50 | semi-buried (85.0%) | 61,679 |
| 1I05<sup>75</sup> | Major urinary protein 1 | 6-hydroxy-6-methyl-heptan-3-one | 2.0 | buried (95.8%) | 51,743 |
| 2XJG<sup>77</sup> | Heat shock protein HSP90-α | (1,3-dihydro-isoindol-2-yl)-(2-hydroxy-5-isopropyl-4-methoxy-phenyl)-methanone | 2.25 | semi-buried (82.0%) | 37,806 |
| 2XAB<sup>77</sup> | Heat shock protein HSP90-α | (1,3-dihydro-isoindol-2-yl)-(2,4-dihydroxy-5-isopropyl-phenyl)-methanone | 1.90 | semi-buried (83.8%) | 57,296 |
| 5I40<sup>76</sup> | Bromo - domain containing protein 9 | 6-methyl-1,6-dihydro-7H-pyrrolo[2,3-c]pyridin-7-one | 1.04 | buried (92.4%) | 37,935 |

\*To provide a rigorous quantitative classification of the binding environments, we evaluated the degree of ligand buriedness by calculating the shielded fraction of its solvent-accessible surface area (SASA). Using a standard 1.4 Å probe radius, we computed the free SASA of the isolated ligand in vacuum ( $SASA_{free}$ ) and the exposed SASA of the ligand when bound within the protein complex ( $SASA_{bound}$ ). The shielded percentage was derived using the relation $(1 - SASA_{bound}/SASA_{free}) \times 100\%$ . We defined a binding site as "buried" if $\geq 90\%$ of the ligand's surface area is occluded by the protein (e.g., 1I05 and 5I40), and "semi-buried" if the shielding falls between 70% and 90% (e.g., 5I1Q, 2XAB, and 2XJG), indicating significant partial exposure to the bulk solvent.

**Table 2.**
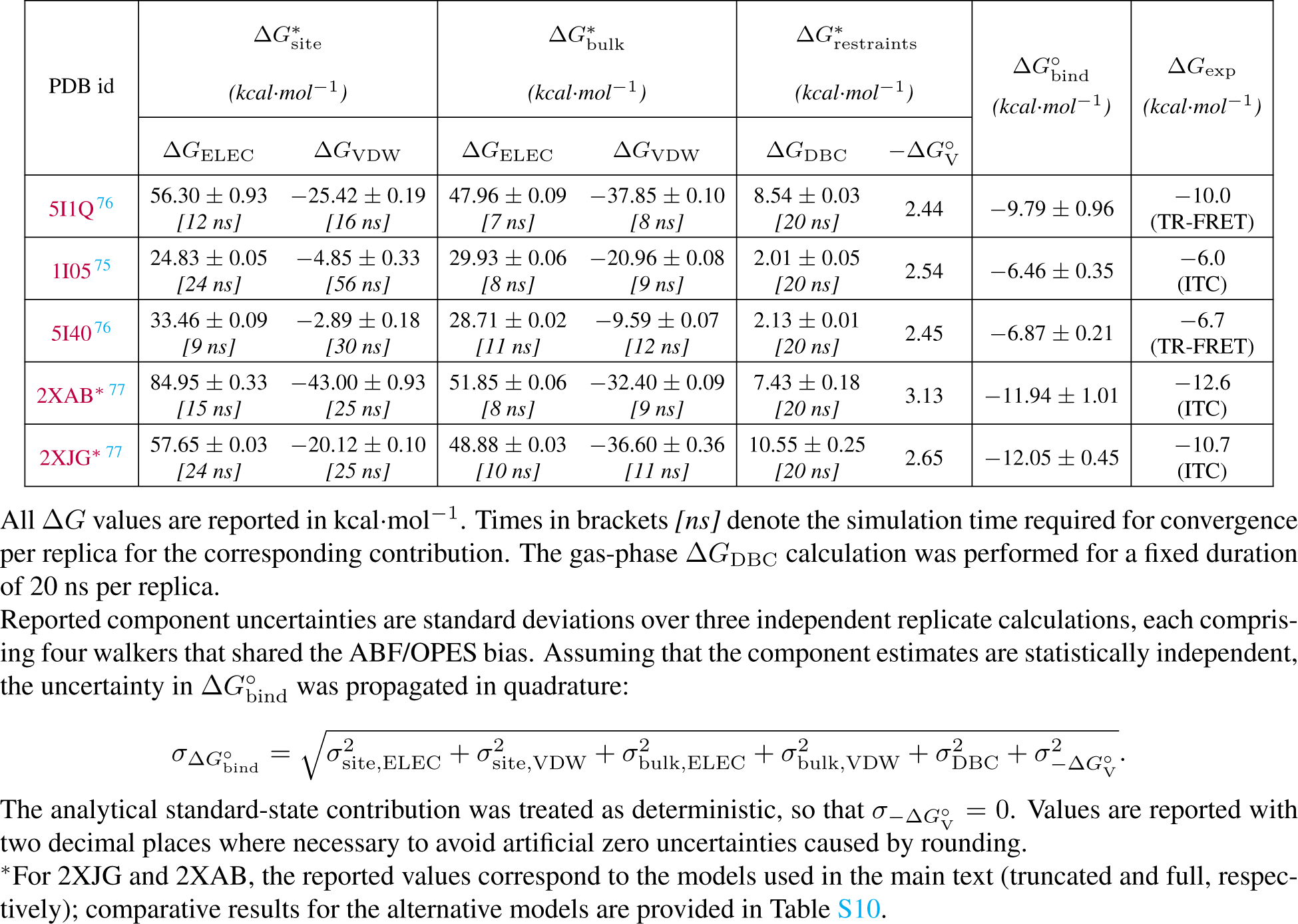
Alchemical contributions to the standard binding free energy and convergence times per replica.

## Methods

### Molecular Dynamics Protocol

Each protein–ligand complex was first prepared using CHARMM-GUI v3.8^78,79^ to generate a fully solvated system with neutralizing counterions and 0.15 M NaCl. The resulting PDB file was then split into separate protein, ligand, water, and ion components, each converted to Tinker XYZ format using the *pdbxyz* program from Tinker suite ^80,81^. Ligand coordinates were converted to SDF format and processed with Poltype2^82–84^ to generate the corresponding force-field parameter set, including atomic multipoles and polarizabilities. The complete system was subsequently reassembled using the *xyzedit* program from the same Tinker release to produce the final Tinker-HP input configuration. All molecular interactions were modeled using the polarizable AMOEBA Bio 2018 force field^74,85–87^.

Protein protonation states were assigned using PROPKA 3.1.5 and subsequently inspected in the local hydrogen-bonding environment, with particular attention to histidine tautomeric states. The final protonation-state assignments are reported in Table S1.

Each studied protein–ligand complex underwent energy minimization using a limited-memory Broyden–Fletcher–Goldfarb–Shanno (L-BFGS) algorithm. The minimization was carried out in two steps: first, the complex was minimized with electrostatic and polarization interactions disabled to alleviate steric clashes. In the second step, the minimization was repeated with electrostatic and polarization interactions enabled. The complexes were then pre-equilibrated for 1 ns under sequential NVT and NPT conditions to stabilize the temperature and system density, respectively. MD production simulations in the NPT ensemble (300 K, 1 atm) were then conducted using the BAOAB-RESPA multiple-time-step integrator^88^, implemented in the GPU-accelerated Tinker-HP v1.3^89–91^. Hydrogen mass repartitioning^92^ was applied to enable a 3-fs integration time step.

The simulations were conducted with a van der Waals cutoff of 12 Å and a real-space Ewald cutoff of 7 Å. Long-range electrostatic interactions were evaluated in reciprocal space using particle-mesh Ewald summation. These settings follow previously benchmarked Tinker-HP protocols for AMOEBA simulations.^89^

### Design of the Distance-to-Bound-Configuration Collective Variable

An accurate definition of the DBC CV is essential for restraining the ligand within the binding site while allowing for its relevant conformational flexibility.^44,93^ The DBC is defined as the root-mean-square deviation (RMSD) of the ligand relative to the receptor, requiring two specific atom subsets: (i) a structurally stable subset of ligand atoms, typically chosen for their minimal internal flexibility to represent the ligand orientation and conformation relative to the binding site, and (ii) a set of protein anchor atoms that provide a moving reference frame. Operationally, the protein anchor atoms define the alignment frame in which the ligand RMSD is evaluated.

To identify system-specific ligand–anchor atom subsets, we employed a systematic procedure based on conventional MD trajectory analysis. Atomic stability was quantified by computing the root-mean-square fluctuation (RMSF) over the trajectory, ensuring that only conformationally stable atoms contribute to the DBC calculation. We define the anchor region as the set of C*_α_* atoms located within 6 Å of the ligand in the reference structure. These atoms are spatially close enough to represent the local binding-site environment while minimizing contributions from distal protein motions. From this initial pool, only atoms exhibiting minimal fluctuations (RMSF *<* 0.7 Å) are retained (the corresponding protein RMSF profiles and selected anchor residues are provided in Fig. S1).

Similarly, for the ligand, all heavy atoms are initially considered, and only those with RMSF values below a system-dependent cutoff (typically 0.6–1.0 Å) are selected. The cutoff was chosen to retain a sufficient set of atoms representing the comparatively rigid ligand core; the system-specific selections are reported with the complete DBC definitions in Fig. S1 and Table S2. Such CVs were defined, monitored, and biased throughout the alchemical transformations using Colvars version 2026-09-04^62^, linked to Tinker-HP.

The distribution of DBC values across the trajectory reflects the range of native fluctuations between the ligand and anchor atoms. The upper restraint limit is defined as the 95th percentile of this distribution, accommodating natural motion without artificially biasing the system. The selected ligand and anchor atom subsets were then subjected to a flat-bottom harmonic restraint with a force constant of 100 kcal·mol*^−^*^1^·Å*^−^*^2^. This value follows previous applications of DBC restraints in absolute binding free-energy calculations.^36,44^ The restraint exerts no force within the flat-bottom region and limits excursions only when the DBC exceeds the prescribed upper boundary. Reference coordinates are taken from the last frame of the conventional MD trajectory. This selective procedure not only provides a physically meaningful definition of the DBC CV but also offers an implicit check on complex dynamics. For instance, if a stringent DBC selection threshold results in very few anchor atoms being selected, this may indicate substantial protein flexibility at the binding site and motivate reassessment of the anchor definition.

Water atoms were intentionally excluded from the DBC definition so that the restraint acted only on solute coordinates and did not prescribe the identity or occupancy of binding-site waters. Including water coordinates would couple the restraint to molecular water identity and could impede exchange between crystallographic sites and the bulk. Their exclusion therefore allows hydration and rehydration to occur as part of the response to alchemical decoupling rather than as a restraint-imposed process.

To enhance sampling efficiency and improve convergence of the free-energy profiles, we employed the enhanced-sampling strategy described in the next subsection.

### Lambda-ABF-OPES Enhanced Sampling for Binding Free Energy

Accurate binding free-energy calculations require efficient sampling of the extended alchemical coordinate, *λ*, which governs the decoupling of the ligand and the accompanying reorganization of solvent and protein. To address this, we employed Lambda-ABF-OPES,^55^ a hybrid scheme that combines the multiple-walker ABF algorithm^59,94^ with OPES*_e_*, an adaptive enhanced-sampling technique used to promote exploration of the alchemical coordinate.^58,95,96^ In practice, the alchemical variable is treated as an extended CV via the Colvars library^62^ coupled with Tinker-HP.^97,98^ Four independent walkers were run in parallel for each electrostatic and van der Waals free-energy contribution in the bound and unbound states. Within each calculation, the walkers shared the evolving biasing information. Three independent replicate calculations were performed for every contribution, each replicate comprising four walkers.

The smooth evolution of *λ* is propagated using Langevin dynamics. The ABF component estimates the mean force along *λ* and applies the corresponding adaptive bias; the multiple walkers share accumulated mean-force information.^59,94^ OPES*_e_* constructs an adaptive bias from a kernel-density estimate of the sampled distribution using a well-tempered target.^37,95,99,100^ In the combined scheme, ABF provides the mean-force estimator, whereas OPES*_e_* promotes repeated exploration of the alchemical coordinate.

AMOEBA defines the polarizable molecular interaction model, while the dynamic alchemical coordinate continuously modulates the ligand steric and electrostatic coupling to its environment. The combined bias acts directly on *λ* and can facilitate relaxation of solvent, ligand, and binding-site degrees of freedom when these motions are coupled to the alchemical transformation; it does not, however, guarantee equilibration of slow coordinates that are weakly coupled to *λ*. Further technical details on Lambda-ABF-OPES are provided in Refs. 54,55. The relative contributions of selected sampling components were further examined using dedicated control calculations reported in the Supporting Information (see Figs S26-S31).

Building on this sampling framework, we next describe the thermodynamic cycle and restraint strategy used to compute standard binding free energies, including the application of DBC restraints and the decomposition of van der Waals and electrostatic contributions along the alchemical pathway.

### Thermodynamic Framework and Restraint Strategy

The standard binding free energy, 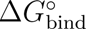, was evaluated using the alchemical thermodynamic cycle shown in Fig. 2. Rather than simulating physical ligand association or dissociation, the cycle connects the bound and unbound states through alchemical transformations in the binding site and in bulk solvent, together with a restraint correction for the fully decoupled ligand. Throughout this work, 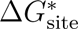 and 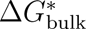 are both defined in the decoupling direction. Thus, 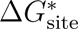 is the free-energy change for restrained decoupling of the ligand from the binding site, whereas 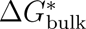 is the free-energy change for unrestrained decoupling of the ligand from bulk solvent. The thermodynamic cycle traverses the binding-site leg in the reverse direction and the bulk-solvent leg in the decoupling direction. Consequently, these terms enter the standard binding free energy as 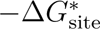 and 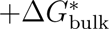, respectively. The three computational steps shown in Fig. 2 are described below.

**Figure 2.**
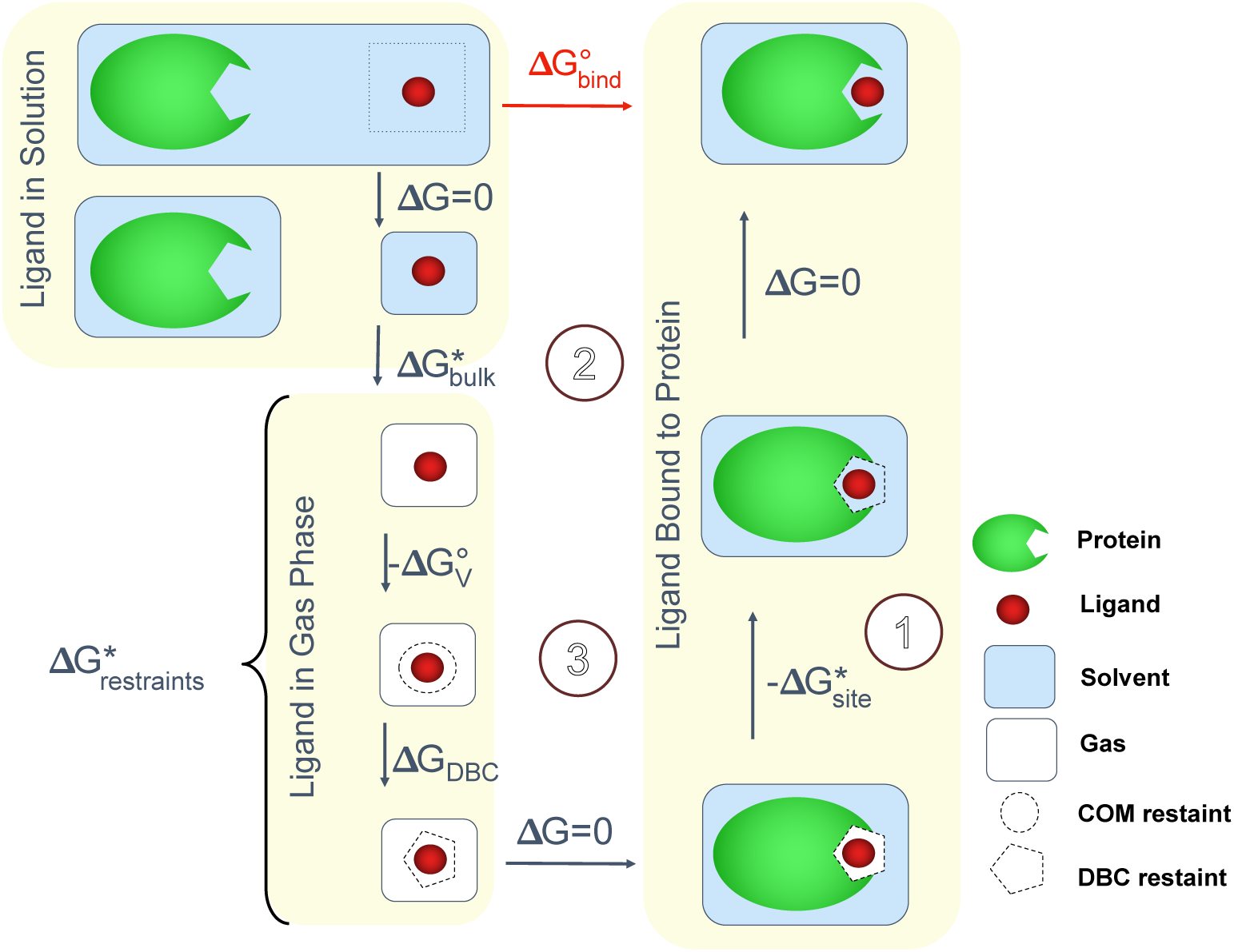
Thermodynamic cycle used to compute the standard binding free energy, 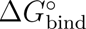. The arrows follow the direction used to assemble the binding free energy. In step 1, the DBC-restrained, fully decoupled ligand is coupled to the protein binding site. This is the reverse of the simulated binding-site decoupling transformation and therefore contributes 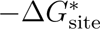. In step 2, the unbound ligand is decoupled from bulk solvent without positional or orientational restraints, contributing 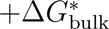. Step 3 applies the restraints to the fully decoupled ligand and contributes 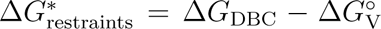. Here, Δ*G*_DBC_ is the work required to apply the DBC restraint while the COM restraint is maintained, and 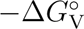 is the analytical work associated with confining the ligand COM from the 1-M standard-state volume to the finite COM-restrained volume. The DBC restraint is inactive over the sampled fully coupled bound-state distribution; its removal from the bound complex is therefore treated as having a negligible free-energy contribution. Protein, ligand, solvent, and fully decoupled states are shown in green, red, light blue, and white, respectively. A dashed circle denotes the COM restraint, and a dashed pentagon denotes the DBC restraint.

#### Binding-site transformation (step ①)

In the bound complex, a distance-to-bound-configuration (DBC) flat-bottom harmonic restraint was applied to prevent the ligand from drifting away from its reference pose as its interactions with the environment were removed. The corresponding calculation yields 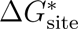 for restrained decoupling of the ligand from the binding site. Because step ① in the binding cycle is traversed in the opposite direction—from the restrained, fully decoupled state to the coupled bound state—its contribution to 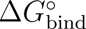 is 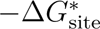.

Within the AMOEBA force field,^65,74,86^ hydrogen bonding and other specific noncovalent contacts are not introduced as separate interaction terms. Instead, they arise from the combined van der Waals and electrostatic interactions, including permanent multipoles and explicit polarization.

The electrostatic and van der Waals contributions were evaluated in successive alchemical legs. Along the global alchemical path used in this work, the van der Waals leg spans 0 ≤ *λ <* 0.5, whereas the electrostatic leg spans 0.5 ≤ *λ* ≤ 1. In the decoupling direction, the electrostatic interactions are therefore removed first, as the global coordinate decreases from 1 to 0.5, while steric exclusion is retained. The van der Waals interactions are then removed as the coordinate decreases from 0.5 to 0. In the reverse coupling direction, the steric interactions are introduced before the electrostatic interactions. This ordering avoids activating electrostatic interactions in the absence of a steric core and prevents the associated end-point singularities.^101–103^

A soft-core potential^104,105^ was used for the van der Waals transformation to avoid singularities and atomic overlap. The buffered 14–7 soft-core potential was

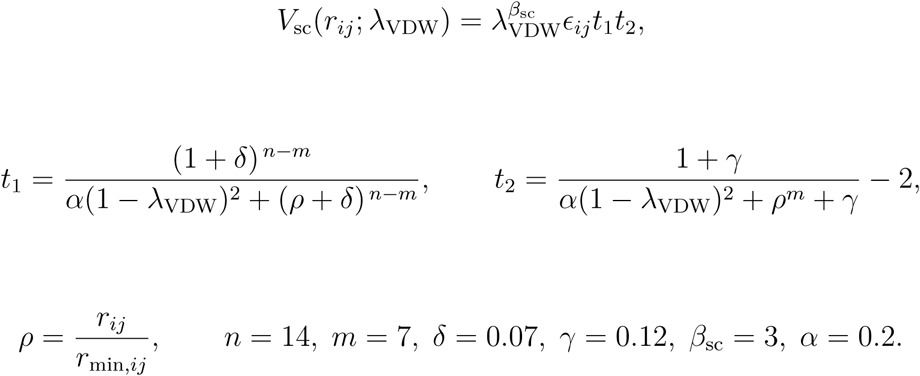

Here, *r_ij_* is the interatomic distance, *ɛ_ij_* is the well depth, and *r*_min_*_,ij_* is the minimum-energy separation. The quantity *λ*_VDW_ is the leg-specific van der Waals coupling coordinate, which varies from 0 in the fully decoupled state to 1 in the fully coupled state; it should not be confused with the global coordinate used to index the two successive alchemical legs. The parameters *β*_sc_ = 3 and *α* = 0.2 control the soft-core scaling and the smoothness of the interpolation, respectively.

#### Bulk-solvent decoupling (step ②)

The unbound ligand was decoupled from bulk solvent without positional or orientational restraints. This transformation removes the ligand–solvent interactions and yields 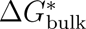, the free-energy change for transferring the solvated ligand to the fully decoupled state. As in the binding-site calculation, 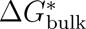 was resolved into electrostatic and van der Waals increments. Because the thermodynamic cycle traverses this leg in the same direction as the calculation, it contributes 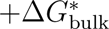 to the standard binding free energy.

#### Fully decoupled restraint correction (step ③)

The DBC flat-bottom restraint was chosen to remain inactive over the sampled distribution of the fully coupled bound state. Its removal from that state was therefore treated as having a negligible free-energy contribution, as indicated by the Δ*G* = 0 leg in Fig. 2.

For the fully decoupled ligand, however, the DBC restraint restricts departures from the reference bound configuration. Its free-energy contribution, Δ*G*_DBC_, was evaluated in a separate thermodynamic-integration (TI) calculation while a flat-bottom center-of-mass (COM) restraint confined the ligand to a finite translational volume. Because the ligand was fully decoupled from its environment, this calculation contains no ligand–environment interaction contribution.^64^ The DBC restraint potential, *U*_DBC_, was progressively coupled from *λ*_DBC_ = 0 (restraint absent) to *λ*_DBC_ = 1 (restraint fully applied), giving

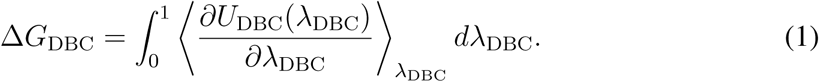

The finite volume imposed by the COM restraint was then related analytically to the 1-M standard-state volume. Treating the ligand COM as a point particle confined within the spherical flat-bottom region gives

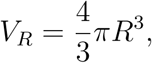

where *R* is the upper boundary of the COM restraint. The standard-state volume per molecule is

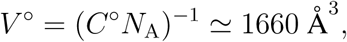

where *C^◦^* = 1 mol L*^−^*^1^ and *N*_A_ is Avogadro’s constant. In the direction followed by step ③, the ligand COM is confined from *V ^◦^* to *V_R_*. The corresponding contribution is

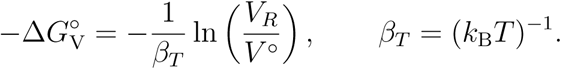

The subscript on *β_T_* distinguishes the inverse thermal energy from the unrelated soft-core exponent *β*_sc_. The complete fully decoupled restraint contribution is therefore

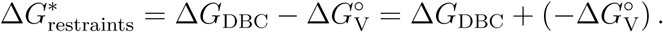

Three independent TI calculations were performed to estimate the mean and replicate variability of Δ*G*_DBC_. In each calculation, the DBC restraint was alchemically coupled while a fixed flat-bottom COM restraint confined the decoupled ligand to a finite translational volume. Both restraints used a force constant of 100 kcal·mol*^−^*^1^ · Å*^−^*^2^. The upper boundary of the DBC restraint was parameterized from the natural thermal fluctuations of the ligand in the bound state, as determined from unbiased MD simulations. The complete DBC definitions used in the production calculations are presented in Fig. S1 and Table S2.

The COM restraint prevented diffusive translation of the decoupled ligand away from the reference binding-site region.^44^ Its upper wall was assigned from the system-specific DBC width. When the DBC width was below 1 Å, an additional offset of 1 Å was applied to avoid overly restricting ligand translation. For a DBC width greater than or equal to 1 Å, the broader DBC boundary was taken to provide sufficient translational flexibility, and the COM upper wall was set equal to the DBC width.

Following the direction and notation of the thermodynamic cycle in Fig. 2, the standard binding free energy is

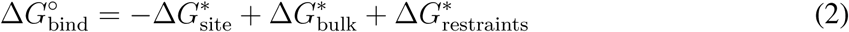

This equation is used without any further sign conversion in resulting site and bulk quantities of 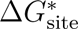 and 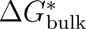 in their respective decoupling directions (Table 2).

The last two terms together define the gas-phase restraint contribution, 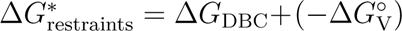.

## Results and Discussion

The decomposed free-energy contributions, together with the calculated and experimental binding affinities for all five complexes (Fig. 1), are summarized in Table 2. For each system, the electrostatic and van der Waals increments were first summed within the binding-site and bulk-solvent decoupling legs. Equation 2 was then applied by subtracting the binding-site sum.

We next focus on two representative complexes chosen to illustrate contrasting binding-site hydration regimes. The 5I1Q complex represents a dynamic regime in which the interfacial hydration pattern is sustained by both resident and exchanging waters, together with redistribution of local water-mediated polar contacts. By contrast, 1I05 contains a buried, persistent crystallographic hydration network.

These cases should be viewed as complementary hydration regimes rather than as separate mechanistic exceptions. In both systems, the Lambda-ABF-OPES calculations probe the coupled response of the ligand, binding pocket, and local solvent as the ligand interactions are alchemically scaled. What differs is the microscopic expression of that response. In 5I1Q, the interfacial hydration pattern is maintained through exchange among resident and incoming waters and through redistribution of local water-mediated polar contacts. In 1I05, the principal crystallographic hydration motif is already persistent during conventional MD, and the alchemical calculation must sample its response together with the steric, cavity, and local-solvent relaxation that accompanies ligand decoupling. Together, the two systems test whether the same alchemical protocol can accommodate both persistent and exchange-mediated hydration without explicitly biasing water-based collective variables.

All necessary starting configurations and input files, including the DBC collective variables used in the condensed-phase simulations (conventional MD and binding free-energy calculations), are provided in the Lambda-ABF-OPES GitHub repository^106^. The remaining three complexes provide an additional assessment across different proteins and pocket topologies and are discussed more concisely below; additional binding-affinity and water-mobility analyses are reported in the SI.

### 5I1Q: Semi-buried Ligand Binding

#### Conventional MD Analysis

The second bromodomain of TAF1 (TAF1(2)) is a reader domain that recognizes acetylated lysines on histones and other proteins, thereby playing a critical role in transcriptional regulation. Its acetyl-lysine binding cavity contains conserved asparagine and tyrosine residues that participate in direct and water-mediated recognition. While this function is essential for normal cellular processes, dysregulation of TAF1(2) has been implicated in neurodegeneration, cancer, and cardiovascular disorders.^107–109^ Small-molecule inhibitors of TAF1(2) commonly exploit the hydrogen-bonding and hydrophobic features of this pocket, including interactions that mimic acetyl-lysine recognition.^76,110,111^ Among these, pyrrolopyridone derivatives have emerged as promising scaffolds due to their capacity to create both direct and water-mediated interactions within the bromodomain site.

We first examined the structural basis of binding of butenyl-pyrrolo-pyridinone-dimethylbenzamide (compound 67C), shown in Fig. 3A. The crystal structure reported by Crawford et al.^76^ reveals a water-mediated hydrogen-bonding network that bridges the ligand and conserved residues within the binding site. Hydrophobic stabilization is provided by the dimethylbenzamide moiety, while the butenyl side chain contributes conformational adaptability that may optimize ligand accommodation within the pocket. To characterize the persistence and reorganization of these interactions, we performed a 100-ns conventional MD simulation and analyzed the protein–ligand polar contacts, local solvation structure, and site-specific water occupancy (Fig. 3).

**Figure 3.**
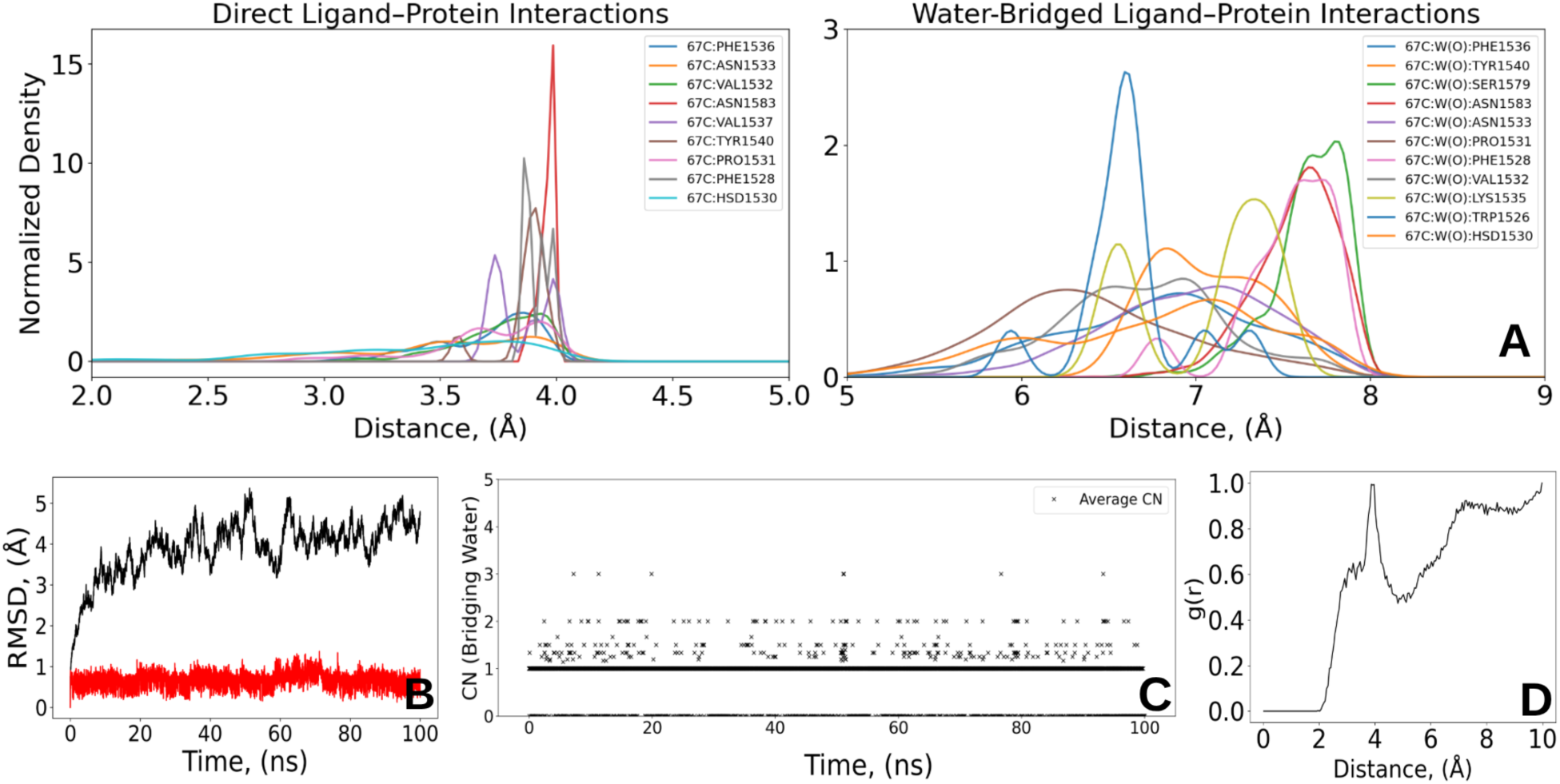
Structural analysis of the second bromodomain of TAF1(2) in complex with the semi-buried pyrrolopyridone derivative (compound 67C) (PDB id: 5I1Q)^76^. (A) Density distributions of protein-ligand polar contacts using a 4 Å heavy-atom cutoff: direct contacts between protein and ligand oxygen atoms (left) and water-mediated (O-bridging) contacts involving protein and ligand oxygen atoms (right). Atom-level assignments are reported in Table S3.. Atom-level assignments for the corresponding contacts are reported in Table S3. (B) RMSD evolution of protein C*_α_* atoms (black) and ligand heavy atoms (red) over the 100-ns conventional MD simulation. (C) Average coordination number (CN) of O-bridging water molecules involved in mediating oxygen atoms of protein–ligand interactions (calculated within a 4 Å cutoff). (D) Radial distribution function, g(r), between the ligand heteroatoms and the oxygen atoms of the surrounding water molecules. Source data are provided in Supplementary Data 1.

The density distributions (Fig. 3A) identify recurrent direct and water-mediated polar contacts at the semi-buried interface. In the O-bridging motif, in which both the protein and ligand contact atoms are oxygen atoms, PHE1536, PRO1531, SER1579, ASN1583, and ASN1533 contribute prominent residue-associated features, with TYR1540 providing an additional recurrent hydration anchor (Fig. 3A and Fig. S4). The hydration network is further reinforced by N-, O/N, and N/O water-mediated polar-contact motifs, where the motif labels denote protein-element/ligand-element heavy-atom pairs (see Fig. S5-S7). Because the figure legends use compact residue-level labels, the corresponding atom-level assignments are reported in Tables S3. For instance, the dominant ASN1583 contact corresponds to an extended N· · ·O heavy-atom polar contact between ligand O2 and the ASN1583 side-chain ND2 atom, with a mean distance of 3.80 Å and an occupancy of approximately 0.92 (Table S3). We therefore describe this contact as an extended heavy-atom polar-contact peak rather than as a canonical strong direct hydrogen bond. ASN1533 and HSD1530 also contribute through the backbone and side-chain atoms specified in Table S3. Water-mediated contacts involving a protein oxygen and a ligand nitrogen are additionally associated with PHE1536 and PRO1531 (O/N-bridging, Fig. S7). Together, these analyses delineate a heterogeneous interfacial network, in which some residues participate in both direct and water-mediated contacts, whereas others contribute predominantly through solvent-mediated interactions. Atom-resolved assignments for all five complexes are provided in the SI.

The RMSD profiles in Fig. 3B demonstrate that the protein C*_α_* RMSD fluctuates around 4–5 Å, whereas the ligand remains near 1 Å. Thus, the ligand preserves a well-defined bound pose despite larger-scale protein motions during the trajectory. The O-bridging coordination number (CN) analysis (Fig. 3C) reveals a dynamic hydration profile, with the continuous CN predominantly sampling values between approximately two and three. To extend this observation, we computed individual CNs for each water-mediated protein-ligand pair (Fig. S4-S7), which exhibited intermittent fluctuations consistent with recurrent reorganization of water-mediated contacts. Because such aggregate and pairwise CN descriptors do not by themselves identify which crystallographic water site is occupied or whether a given site preserves its protein–water–ligand geometry, we further performed the site-resolved W1/W2 analysis described below.

Previous studies showed that water displacement in bromodomain complexes involves a delicate interplay between ligand binding and hydration network reorganization.^9,17,76^ We therefore examined two crystallographic hydration sites, W1 and W2, located between the ligand polar region and neighboring protein residues in the TAF1(2)–compound 67C complex (Fig. 4A).

**Figure 4.**
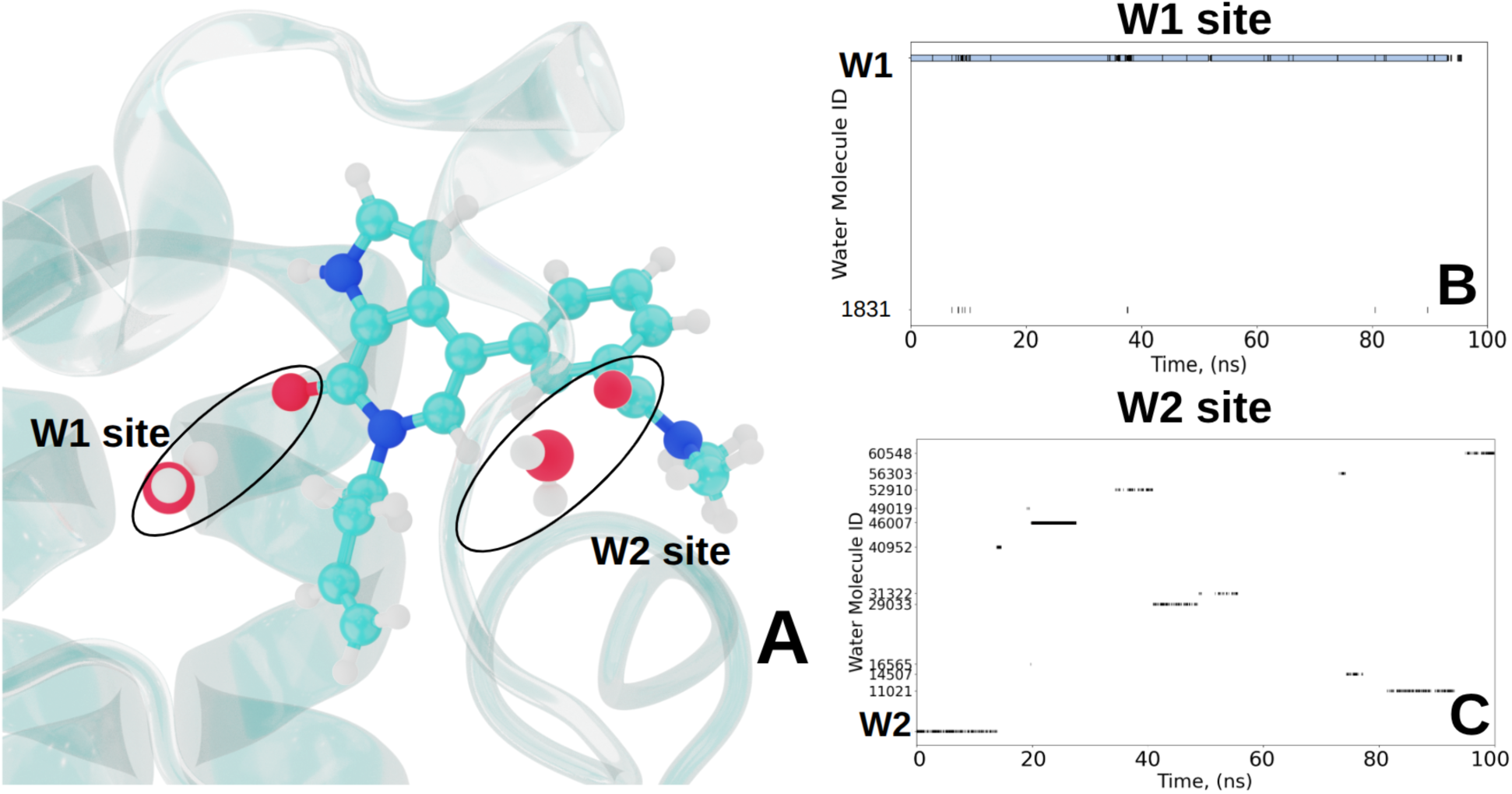
Site-specific water occupancy and identity in the TAF1(2) binding pocket (PDB id: 5I1Q). (A) Structural overview of the selected crystallographic hydration sites W1 and W2 (red spheres), located between the ligand polar region and neighboring protein residues. (B) Time-resolved nearest-water identity for the W1 site. The blue horizontal trace corresponds to the original crystallographic water, which remains associated with the W1 region for most of the trajectory before the site becomes depleted after approximately 93 ns. (C) Time-resolved nearest-water identity for W2, showing recurrent occupancy by multiple distinct water molecules. Source data are provided in Supplementary Data 1.

First, we assessed whether variation in the total ligand dipole moment covaried with the aggregate CN of bridging waters. No significant correlation was detected (Fig. S8), arguing against a simple correspondence between the global ligand dipole and the hydration fluctuations measured by this descriptor. This observation motivated a site-resolved analysis of W1 and W2.

To distinguish close localization at the crystallographic position from broader occupation of the surrounding hydration region, we applied two distance criteria around each reference water oxygen. A water located within 2.0 Å was assigned to the crystallographic core, whereas the 3.5 Å exchange shell captured waters occupying the broader site region during molecular exchange. The two sites displayed distinct behavior. W1 remained occupied by the original crystallographic water molecule for most of the trajectory, with near-continuous core occupancy until approximately 93 ns. This water molecule then left both the 2.0 Å core and the broader 3.5 Å shell, after which the W1 site remained largely depleted without stable occupation by a replacement water. By contrast, the original crystallographic water at W2 was not retained. The W2 region was recurrently occupied by different water molecules, with a transient depletion followed by rehydration by exchange waters later in the trajectory. Thus, W1 behaves as a persistent site that undergoes late depletion, whereas W2 behaves as a dynamically exchanging hydration site.

We next examined whether occupation of each site preserved the corresponding protein–water– ligand bridge geometry (Figs. S2 and S3). Before its depletion, the W1 water remains primarily anchored to the protein through the TYR1540 hydroxyl group. Its ligand-side interaction, however, does not preserve the crystallographic bridge geometry over the analyzed late-trajectory interval. Instead, the nearest ligand polar partners are predominantly the carbonyl oxygen and the adjacent ring nitrogen of the pyrrolopyridone moiety. W1 therefore contributes to the broader O-bridging coordination number through a redistributed ligand-side contact rather than through continuous preservation of the crystallographic bridge. Decomposition of the aggregate O-bridging CN into W1, W2, and remaining-water contributions further shows that loss of W1 removes a recurrent site-specific component but does not produce a one-to-one decrease in the total CN (Fig. 3C), because waters occupying W2 and other regions of the pocket continue to contribute independently. The results therefore distinguish three related but non-equivalent properties: occupation of a hydration site, persistence of a particular water molecule, and preservation of the complete protein–water– ligand bridge geometry.

Providing a broader perspective, the radial distribution function (RDF) between ligand heteroatoms and water oxygen atoms shows a pronounced first peak at ∼3.3 Å (Fig. 3D), consistent with first-shell ligand–water polar contacts,, followed by a secondary rise that reflects the presence of the structured solvation shell around the semi-buried ligand. Together, these findings show that the TAF1(2) hydration network combines persistent local organization with molecular water exchange. This observation is consistent with observations by Aldeghi et al.^17^, who showed that structured waters in bromodomain binding sites may be retained or displaced depending on the bound ligand.

#### Binding Affinity Calculation

To evaluate the binding affinity of the TAF1(2)–pyrrolopyridone compound, we determined the optimal set of protein and ligand atoms for the DBC restraint following the procedure described above. For this complex, the DBC upper boundary was set to 0.8 Å, and the selected protein and ligand atoms used in the calculation shown in Fig. 1A-B and in Fig. S1.

The detailed contributions to the standard binding free energy are summarized in Table 2. Both van der Waals and electrostatic terms exhibit large magnitudes in the site (bound) and bulk (unbound) calculations (see Fig. 5 (A-D)), however, it is their differences that determine the net binding affinity. The van der Waals contribution favors binding by −12.43 kcal·mol*^−^*^1^ (i.e., 37.85 − 25.42 kcal·mol*^−^*^1^), whereas the electrostatic term provides an additional stabilization of −8.34 kcal·mol*^−^*^1^. These results demonstrate that both van der Waals and electrostatic interactions contribute favorably to binding, with van der Waals interactions providing the dominant stabilization. After including the standard-state volume and the DBC restraint corrections, the resulting standard binding free energy is −9.79 ± 0.96 kcal·mol*^−^*^1^ which is in close agreement with experimental value (ΔG_exp_ =−10.0 kcal·mol*^−^*^1^, Table 2).^76^ It should be noted that this level of accuracy was achieved within a relatively short simulation time required to reach the convergence, highlighting the efficiency of our approach (see Fig. S9-S11).

**Figure 5.**
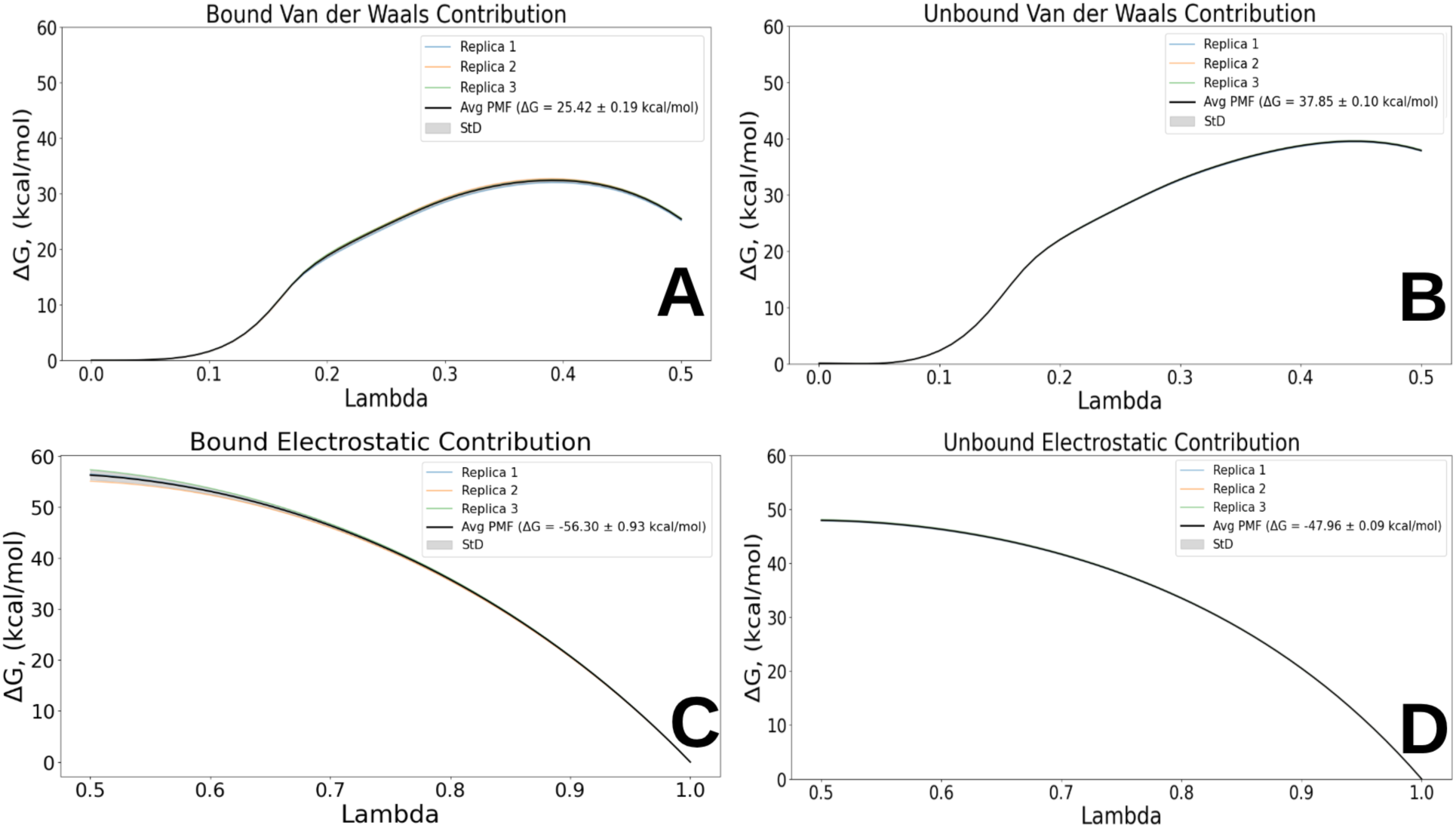
Potential-of-mean-force (PMF) profiles for the binding-site van der Waals (A), bulk-solvent van der Waals (B), binding-site electrostatic (C), and bulk-solvent electrostatic (D) decoupling calculations for the TAF1(2)–compound 67C complex (PDB id: 5I1Q). Each profile is shown for three independent replicate calculations, each comprising four walkers sharing the evolving bias. Source data are provided in Supplementary Data 1.

The critical role of dynamic binding pocket resolvation sampling in this complex was explicitly highlighted by Ge et al.^9^, who noted that extensive ligand motion within the binding site can hinder the insertion or rearrangement of water molecules. Such behavior poses a particular challenge for binding affinity calculations, as it can lead to poor convergence and unreliable free-energy estimates. These observations underscore the importance of accurately modeling ligand-induced water displacement to achieve robust and reproducible results.

To evaluate how effectively Lambda-ABF-OPES samples the solvent response during alchemical decoupling, we monitored the continuous CN of O-bridging water molecules mediating protein– ligand oxygen contacts across all three replicas using a 4 Å cutoff. This cutoff captures the first-shell water-mediated contact region around polar protein and ligand atoms while remaining permissive enough to include distorted or exchanging bridge geometries in the dynamic binding pocket. Comparing the alchemical sampling (Fig. S12-S13) to the corresponding distribution derived from conventional MD (Fig. 3C), we observe that the average CN converges to a mean continuous coordination number of approximately two across both electrostatic and van der Waals transformations. This value closely matches the equilibrium hydration level of the bound state, indicating that the enhanced sampling protocol recovers the expected structural hydration level despite continuous perturbation of the interaction potential. During the van der Waals transformation, increased water coordination is sampled predominantly at low *λ* values (0.0–0.2), where electrostatic interactions are absent and the ligand excluded volume is strongly reduced. Water molecules can consequently enter configurations that are inaccessible in the fully coupled state. As the van der Waals interactions are restored, the CN returns toward the range characteristic of the coupled-state reference. These observations indicate that the solvent population responds reversibly to modulation of the ligand steric coupling, without requiring an explicitly biased water coordinate.

To determine whether this agreement extended beyond the aggregate number of waters, we decomposed the CN into residue-associated contact motifs classified by the protein/ligand element pair (O/O, N/O, O/N, and N/N; Figs. S14–S19). Both binding-site transformations sample the dominant residue-associated motifs observed in conventional MD, including those involving TYR1540, ASN1533, PRO1531, and HSD1530 (Fig. 3A). This correspondence supports recovery of the principal interfacial contact topology rather than only its average hydration number. The N/O motif, involving a protein nitrogen and ligand oxygen, remains weakly populated at low *λ* during the van der Waals transformation (Fig. S15), consistent with its low density in the conventional-MD contact maps (Figs. S5–S7).

Although the decomposition of bridging motifs (Fig. S12–S19) provides a comprehensive overview of the pocket-solvation landscape, these structural descriptors do not independently establish the absolute thermodynamic stability of individual hydration sites. To characterize the site-specific and *λ*-dependent hydration response more precisely, we focused on the prominent anchor residue TYR1540 (Fig. 3A and Figs. S4 and S7), which exhibits persistent solvent engagement throughout the simulations.

We reconstructed *λ*-conditioned hydration probability distributions by tracking the TYR1540 hydroxyl oxygen using the distance-weighted continuous-CN formalism adapted from Ansari et al.^55^. Unlike discrete counting methods, this approach evaluates hydration using a smooth switching function and avoids integer-quantization artifacts. The resulting maps describe the relative populations of hydration states sampled at different values of *λ* and are not interpreted as independent absolute hydration free-energy surfaces.

The probability distributions in Fig. 6A–B resolve distinct *λ*-dependent hydration regimes across the two alchemical legs using the aggregated sampling of four walkers. During the van der Waals transformation, the distribution broadens markedly at low *λ*, with a higher-coordination population extending toward CN≈ 4, consistent with transient solvent entry as ligand steric interactions are removed. In contrast, the electrostatic transformation exhibits a narrower distribution centered near CN≈ 3, consistent with stronger localization of the local solvent network when the electrostatic interactions are present.

**Figure 6.**
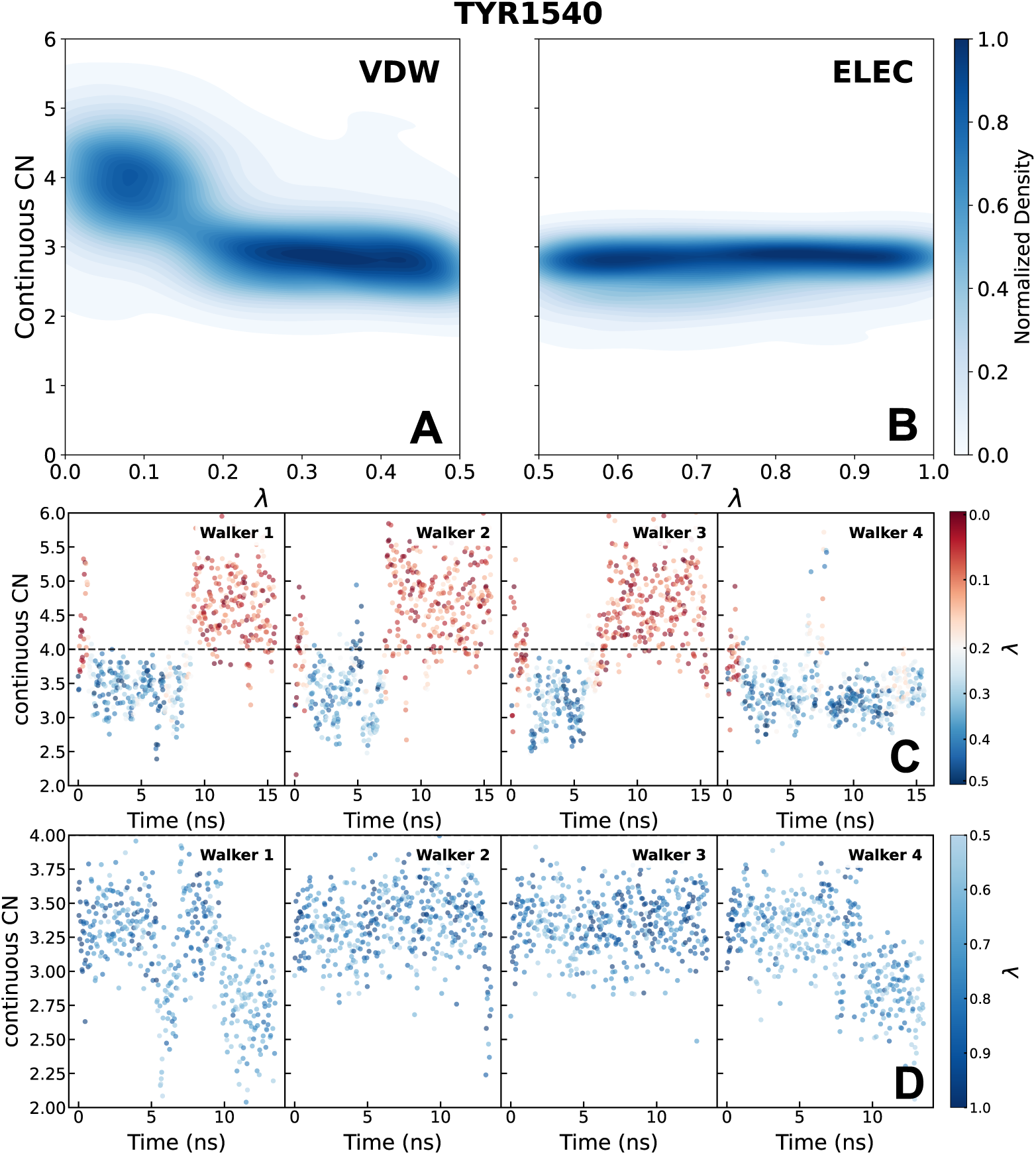
Continuous coordination-number analysis of site-specific hydration around TYR1540. (A,B) Two-dimensional probability-density maps of the continuous 4 Å water coordination number around the TYR1540 hydroxyl oxygen as a function of the alchemical coordinate *λ*. (A) During the van der Waals leg (*λ* : 0.5 → 0.0), the distribution broadens toward higher coordination at *λ <* 0.2, where ligand excluded volume is reduced. (B) During the electrostatic leg (*λ* : 1.0 → 0.5), the distribution remains more narrowly localized. Darker regions indicate more frequently sampled *λ*-conditioned hydration regimes and are not interpreted as independently validated hydration free-energy minima. (C,D) Coordination-number time series across four walkers for the van der Waals and electrostatic transformations, respectively. The traces characterize the reproducibility and structural response of the local hydration environment along the alchemical coordinate. Source data are provided in Supplementary Data 1.

This coordination regime is consistent with our conventional MD analysis, which identified TYR1540 as a participant for multiple solvent bridges. Specifically, it engages in one O-bridging interaction with the ligand oxygen (Fig. S4) and two additional O/N-bridging contacts with the ligand nitrogen (Fig. S7), cumulatively accounting for the observed coordination regime centered near CN≈ 3, supporting structural recovery of the expected local hydration topology within Lambda-ABF-OPES.

To assess whether the observed hydration regimes arise from reproducible solvent exchange rather than from isolated walker-specific events, we examined the continuous CN time series for all four walkers during the van der Waals and electrostatic transformations (Fig. 6C–D). During the van der Waals leg, the walkers repeatedly sample increased TYR1540 coordination at low *λ* ∈ [0, 0.2], where ligand excluded volume is reduced and solvent can enter the expanded pocket. Upon recoupling of the ligand van der Waals interactions, the CN returns toward the more compact bound-state hydration regime.

Although the exact timing and amplitude of these excursions differ among walkers, their repeated occurrence supports a reversible structural response of the hydration shell to steric decoupling. The CN time series therefore provide dynamical evidence that hydration exchange is sampled along the alchemical coordinate.

During the electrostatic leg (Fig. 6D), the TYR1540 coordination number remains comparatively confined around the tri-coordinated regime despite substantial fluctuations of *λ*. This behavior is consistent with stronger electrostatic stabilization of the local solvent network. The overlap of the CN traces across walkers indicates that this electrostatically stabilized hydration motif is sampled consistently, without persistent walker-specific dewetting or overhydration.

Complementary analysis of auxiliary bridging residues ASN1533, ASN1583, and VAL1532 (Fig. S60) reveals related *λ*-dependent hydration populations. Together with the TYR1540 results, these analyses show that the interfacial solvent network contains persistent structural motifs while remaining capable of exchanging individual water molecules during the alchemical transformation.

Because the CN distributions, time series, occupancies, and density maps primarily describe the structural populations of water, we performed an additional matched hydration-suppression control to test whether the observed TYR1540 rehydration response is coupled to the alchemical free-energy profile. The complete protocol, including the restraint definition, force-strength tests, sampling diagnostics, and PMF analysis, is provided in a dedicated subsection of the SI and summarized in Fig. S20. Because the control introduces an artificial *λ*-conditioned perturbation, the resulting PMF changes are interpreted as a sensitivity test rather than as absolute hydration free energies or corrections to the production binding affinity.

Suppressing the low-*λ* TYR1540 hydration transition shifted the binding-site van der Waals free-energy contribution relative to the unrestrained calculation while preserving complete *λ* sampling and comparable DBC behavior. Together with the agreement between the conventional-MD and alchemical hydration analyses, this result shows that Lambda-ABF-OPES effectively samples the exchange-mediated solvent response of the semi-buried 5I1Q pocket along the alchemical coordinate.

### 1I05: Buried Ligand Binding

#### Conventional MD

Major urinary protein I (MUP-I) is a member of the lipocalin family, abundantly present in male mouse urine, where it functions as a pheromone-binding protein.^75,112,113^ Structurally, MUP-I adopts the canonical lipocalin fold: an eight-stranded antiparallel *β*-barrel capped by a short *α*-helix, enclosing a deeply buried hydrophobic cavity (Fig. 7A, B). Its natural ligand, 6-hydroxy-6-methyl-3-heptanone, is a small ketone bearing a polar hydroxyl group central to pheromone signaling.^75,114^ Isothermal titration calorimetry (ITC) measurements reported by Timm et al.^75^ indicate that ligand binding is predominantly enthalpy-driven (Δ*H* = −13 kcal·mol*^−^*^1^), consistent with the extensive noncovalent interactions observed in crystallography, but partially offset by an entropic penalty (−*T* Δ*S* = +7 kcal·mol*^−^*^1^).

**Figure 7.**
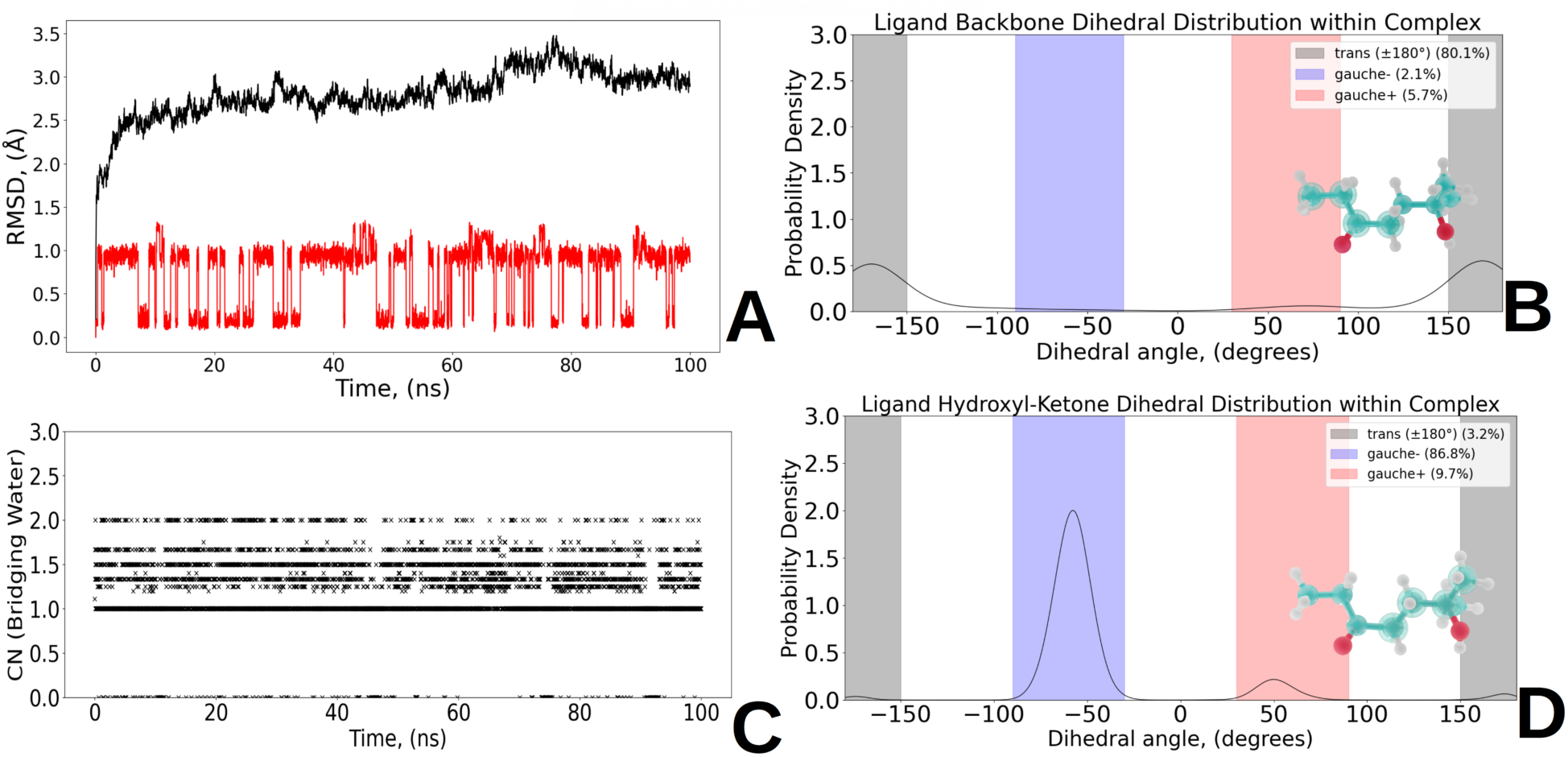
Conventional-MD analysis of the MUP-I–ligand complex (PDB id: 1I05). (A) RMSD of the protein C*_α_* atoms (black) and ligand heavy atoms (red) over the 100-ns trajectory. (B) Probability distributions of the ligand backbone dihedral angles. (C) Time evolution of the continuous coordination number (CN) of waters bridging the protein and ligand. A water was considered bridging when it was simultaneously located within 4.0 Å of at least one ligand oxygen atom and at least one protein oxygen or nitrogen atom. The profile is dominated by a mono-hydrated state near CN≈ 1, with transient excursions toward higher coordination. The mean bridging CN is 1.3. (D) Probability distribution of the ligand hydroxyl–ketone dihedral, showing the trans, gauche+, and gauche− populations.Source data are provided in Supplementary Data 1.

To assess the conformational stability of the MUP-I–ligand complex, we performed a 100-ns conventional MD simulation. The C*_α_* RMSD fluctuated around ∼3.0 Å, indicating that the lipocalin fold remained structurally stable over the trajectory (Fig. 7A). In contrast, the ligand heavy-atom RMSD fluctuated between 0.5 and 1.5 Å, indicating local conformational flexibility within the binding pocket while the ligand preserved its bound pose. To determine which internal motions remained accessible upon binding and whether particular ligand conformers were preferentially populated, we performed a torsional analysis of the ligand backbone and the hydroxyl– ketone dihedral. The backbone dihedrals predominantly adopted trans conformations (∼80%; Fig. 7B), whereas the hydroxyl–ketone dihedral favored the gauche− conformation (∼87%), with minor gauche+ (∼10%) and trans (∼3%) populations (Fig. 7D).

Compared to the TAF1(2)-related complex (i.e., PDB id 5I1Q) discussed above, where continuous water displacement was observed during the simulation, the MUP-I binding site exhibited markedly different behavior. Here, the ligand is deeply buried within the cavity, and a water molecule identified in the crystal structure remained as a persistent bridging water throughout the conventional MD trajectory.

To characterize the observed hydration network, we decomposed the water-mediated protein– ligand contacts according to the interacting protein and ligand atoms (Figs. S21–S22). The ligand is enclosed predominantly by nonpolar cavity residues, which provide extensive van der Waals contacts.^75^ The principal polar contacts are mediated by the two buried waters, which connect the ligand oxygen atoms with TYR138 and LEU58. Atom-level decomposition further assigns these contacts to the TYR138 side-chain hydroxyl and backbone oxygen and to the LEU58 backbone nitrogen and oxygen atoms (Table S4). TYR102 caps the cavity entrance and limits exposure to bulk solvent. Consistent with the crystallographic analysis of Timm et al.,^75^ PHE56 does not participate directly in the polar-contact network but contributes, together with PHE89 and LEU123, to the hydrophobic enclosure of the ligand.

To probe the organization of this buried hydration network, we analyzed correlations among fluctuations in the dipole magnitudes of W1, W2, and their local interaction partners (Fig. 8A–F). The Pearson coefficients quantify linear covariation between these observables and reveal distinct correlation patterns within the hydration network.

**Figure 8.**
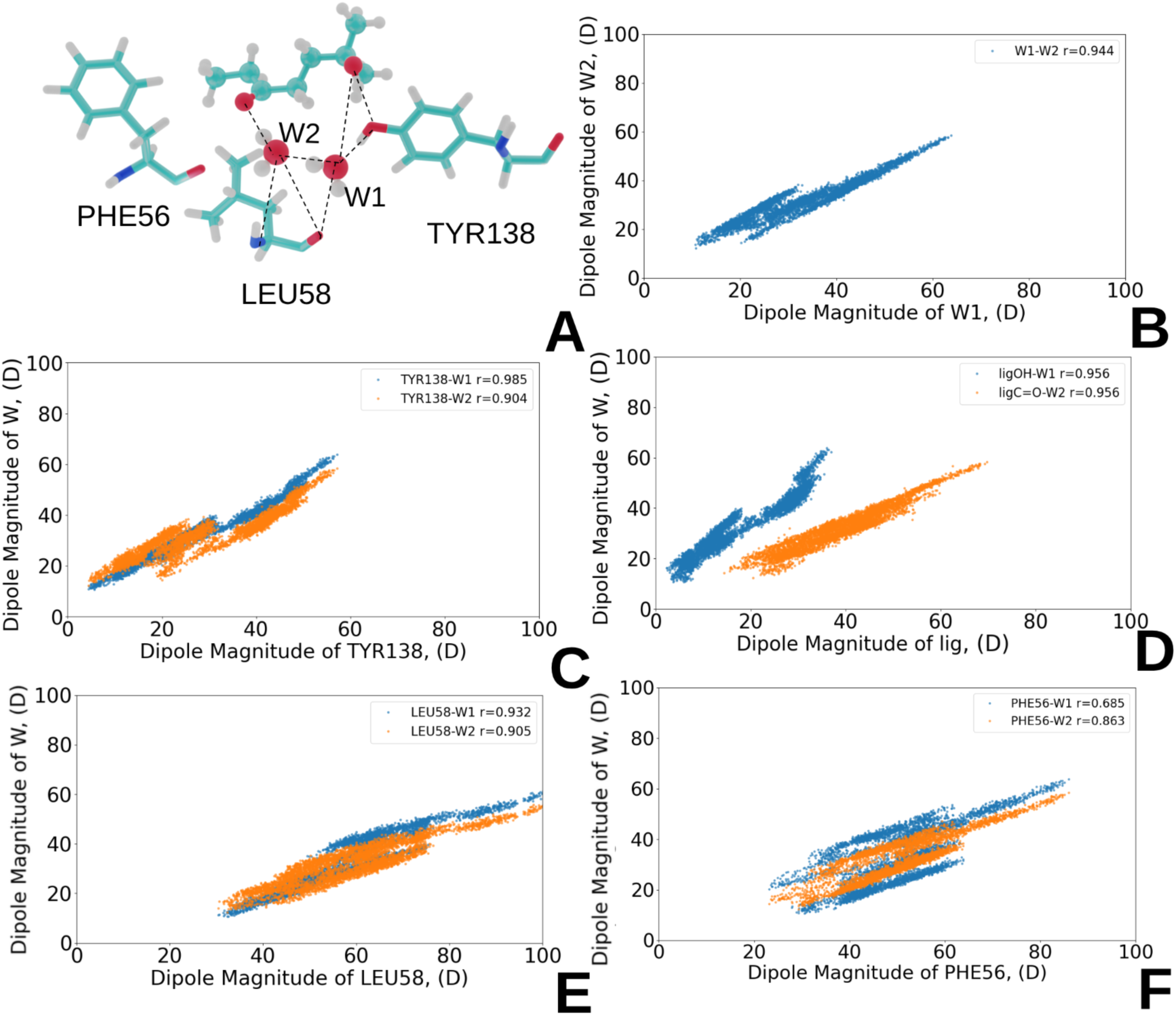
Dipole-magnitude correlations within the buried water-mediated polar-contact network. (A) Structural representation of the network. (B) Correlations between W1 and W2. (C) Correlations of the TYR138 oxygen dipole magnitude with W1 and W2. (D) Correlations of the ligand hydroxyl and carbonyl oxygen dipole magnitudes with W1 and W2. (E) Correlations of the LEU58 oxygen dipole magnitude with W1 and W2. (F) Correlations of the PHE56 oxygen dipole magnitude with W1 and W2. Pearson coefficients, *r*, quantify the linear correlation between the corresponding dipole-magnitude fluctuations.Source data are provided in Supplementary Data 1.

Specifically, W1 displays particularly strong correlations with TYR138 (*r* ≈ 0.99) and LEU58 (*r* ≈ 0.93). Its mean dipole moment, *µ* = 2.8 D with *σ* = 0.2 D (Fig. S23), is slightly larger than the corresponding bulk-water value of approximately 2.7 D. This difference is consistent with the strongly polar and hydrogen-bonded local environment of W1. More generally, *ab initio* MD simulations have shown that water dipole moments increase with hydrogen-bond coordination and local tetrahedral order, approaching ice-like values in highly ordered environments.^115^ The local organization around W1 therefore provides a plausible origin for its increased dipole moment, although tetrahedral order was not evaluated explicitly here.

The persistent occupancy of W1 is also consistent with the NMR relaxation measurements of Denisov et al.^116^, which showed that deeply buried protein waters can be protected from exchange by conformational free-energy barriers in the surrounding protein matrix. In the present trajectory, the strong local dipole correlations, increased polarization, and persistent occupancy of W1 collectively support its assignment as a tightly anchored structural water.

In contrast, W2 exhibits a different correlation pattern. Its correlation with the nearby hydrophobic residue PHE56 is comparatively weak (Fig. 8F), and its mean dipole moment, *µ* = 2.5 D (Fig. S23), is lower than the corresponding bulk-water value. Nevertheless, its dipole fluctuations are strongly correlated with those of both the ligand (*r* ≈ 0.96, Fig. 8D) and W1 (*r* ≈ 0.94, Fig. 8B). These observations are consistent with W1 being anchored primarily by polar protein groups, whereas the organization of W2 depends more strongly on its local relationship with the ligand and W1, together with geometric confinement within the cavity.

#### Binding Affinity Calculation

The 1I05 complex combines two features that complicate alchemical free-energy calculations: internal ligand flexibility and a deeply buried water-mediated interaction network. Previous investigations by Fu et al.^46^ highlighted these difficulties, showing that incomplete exploration of orientational states during the alchemical annihilation stage prevented convergence of free energy estimates. To mitigate this, slow-growth strategy^117^ with carefully tuned restraints were employed to stabilize the binding mode, yet additional complications arose from two buried crystallographic water molecules. Maintaining the experimentally observed hydrogen-bonding network required artificial restraints on these molecules during alchemical transformations, a strategy that, while effective, remains ad hoc and complex-specific, limiting its broader applicability.

In the present study, Lambda-ABF-OPES addresses these sampling challenges without restraining individual water molecules. By combining enhanced sampling of the alchemical coordinate with a DBC restraint that confines the ligand to the bound-state region, the protocol allows the ligand torsional ensemble, cavity, and solvent environment to reorganize repeatedly during decoupling.

Analysis of the dihedral probability distributions shows that while backbone torsions remain centered on the trans conformer under both electrostatic and van der Waals decoupling (Figs. S24-S25), the ketone–hydroxyl torsion displays a broader response (Fig. 9A, C). During van der Waals decoupling, this dihedral remains dominated by the gauche− basin but also samples minor trans and gauche+ populations. During electrostatic decoupling, its distribution more closely resembles the conventional-MD bound-state ensemble, with gauche− remaining predominant. Lambda-ABF-OPES therefore samples additional internal ligand conformations during decoupling without altering the dominant bound-state preference.

**Figure 9.**
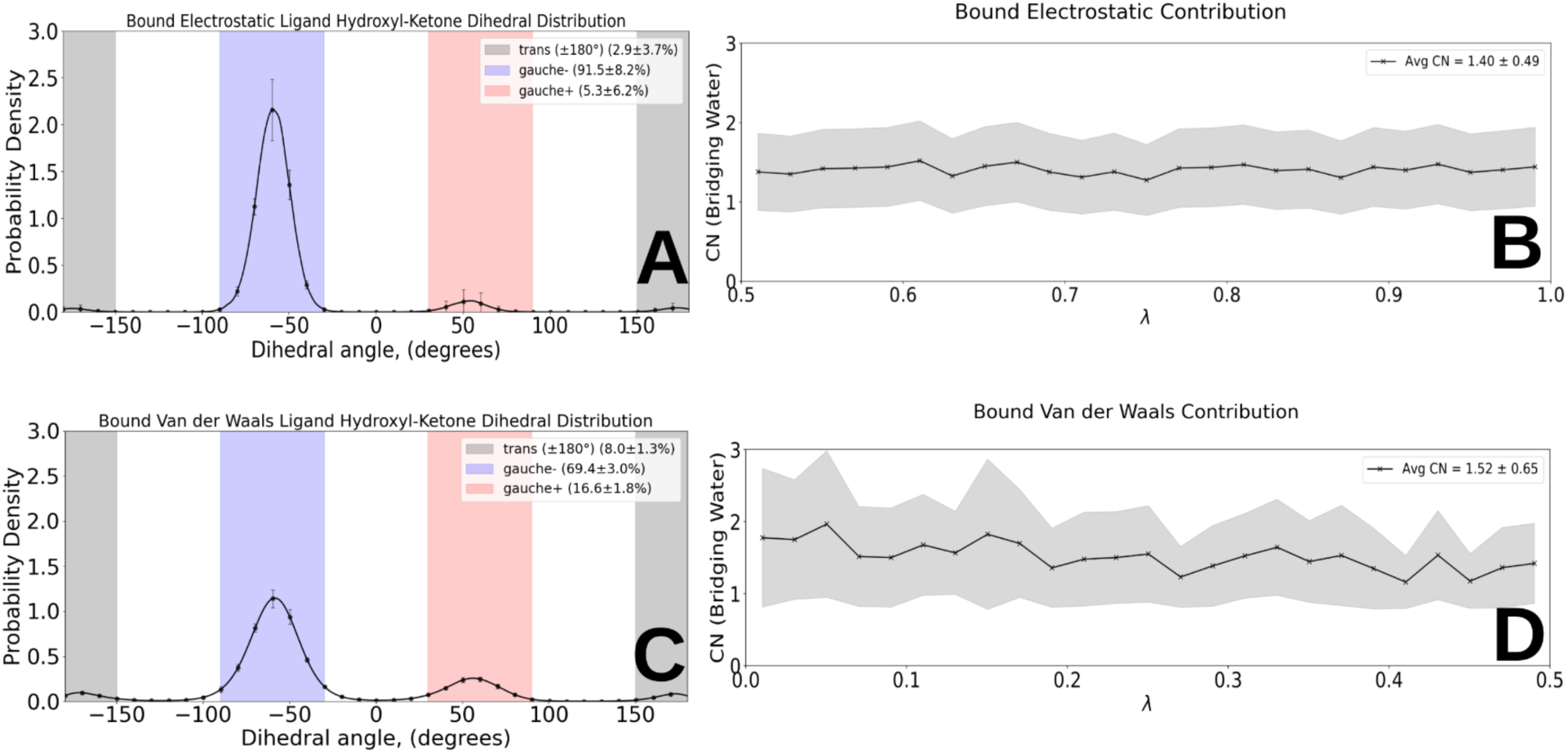
Ligand conformational and bridging-hydration analyses during the binding-site alchemical transformations. (A) Probability-density distribution of the ligand hydroxyl–ketone dihedral during the electrostatic leg. The shaded regions identify the trans (±180*^◦^*), gauche−, and gauche+ states, with their populations reported in the legend. The state populations reported in panels A and C are arithmetic means, and the accompanying uncertainties represent one standard deviation (±1*σ*). (B) Mean continuous CN of bridging waters as a function of the electrostatic coupling parameter *λ*. (C) Corresponding hydroxyl–ketone dihedral distribution during the van der Waals leg. (D) Mean bridging-water CN as a function of the van der Waals coupling parameter. A water was classified as bridging when its oxygen was simultaneously located within 4.0 Å of at least one ligand heteroatom and one protein oxygen or nitrogen atom. In panels B and D, the black trace represents the arithmetic mean of the continuous CN values sampled within each *λ* bin, and the gray band represents one standard deviation (±1*σ*) about that mean. Bridged hydration remains populated throughout both transformations, including the weakly coupled van der Waals states. Source data are provided in Supplementary Data 1.

The buried hydration network is sampled concurrently without restraints on individual waters. The average bridging CN is 1.52 ± 0.65 during the van der Waals leg and 1.40 ± 0.49 during the electrostatic leg (Fig. 9B,D). These values indicate persistent bridged hydration, with intermittent sampling of higher-coordination configurations, throughout both transformations.

The resulting binding free energy (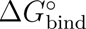 = −6.46 ± 0.35 kcal·mol*^−^*^1^) is in close agreement with experimental affinity data (ΔG_exp_ = −6.0 kcal·mol*^−^*^1^, Table 2), underscoring both the accuracy and robustness of Lambda-ABF-OPES for flexible, buried ligands such as 6-hydroxy-6-methyl-3-heptanone in MUP-I. More broadly, this case study illustrates the framework ability to extend alchemical free energy calculations to other complexes where ligand flexibility, cavity relaxation, and buried hydration have previously hindered convergence.

Each independent Lambda-ABF-OPES replicate required 117 ns of accumulated sampling, comprising 97 ns in the condensed phase and 20 ns for the gas-phase restraint calculation (Table 2). It should be noted that the bound-state VDW leg was the slowest contribution to converge, consistent with the need to sample steric decoupling together with ligand rearrangement and local cavity relaxation. In the present Lambda-ABF-OPES workflow, water exchange and hydrogen-bond reformation occurred spontaneously during the simulation, eliminating the need for artificial water restraints. This outcome demonstrates not only the efficiency of Lambda-ABF-OPES, but also its applicability to buried complexes in which alchemical decoupling must preserve ligand confinement, pocket stability, and local hydration. For 1I05 specifically, the conserved hydration motif is already present in conventional MD. The added value of the present protocol is therefore its ability to maintain this buried water-mediated electrostatic network while sampling the coupled ligand–cavity–solvent relaxation associated with alchemical decoupling.

To characterize the local solvent response along the alchemical coordinate, we analyzed the *λ*-dependent hydration profiles of TYR138 and PHE56 using the continuous-CN formalism applied above to 5I1Q (Fig. 10). Around TYR138, the van der Waals leg contains a weak secondary population near CN≈ 2, reflecting occasional sampling of higher coordination when the ligand electrostatic interactions are absent. During the electrostatic leg, the distribution is more narrowly concentrated near CN≈ 1. This behavior is consistent with a persistent hydration motif at TYR138 and with the conventional-MD dipole analysis, which identified strong correlated fluctuations between W1 and the tyrosine hydroxyl group (Fig. 8). Together, these results show that hydration near TYR138 remains localized as the ligand electrostatic interactions are scaled.

**Figure 10.**
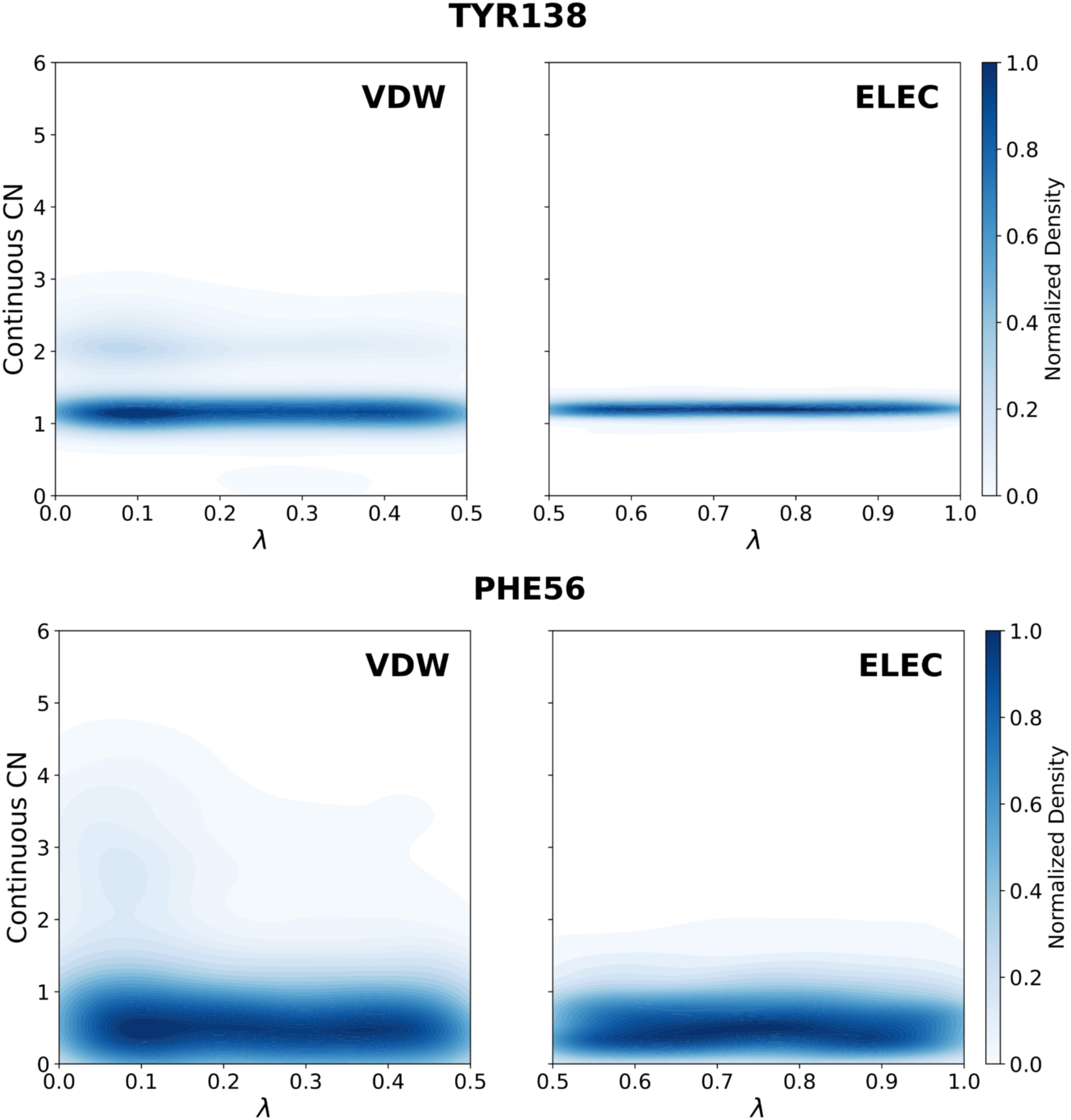
Lambda-dependent hydration profiles of the MUP-I binding pocket. 2D probability density maps of the continuous CN for the oxygen atom of the TYR138 hydroxyl group (top) and the backbone nitrogen atom of PHE56 (bottom) as a function of the alchemical coupling parameter (*λ*) collected over all replicas. (Left) The VDW decoupling leg (*λ* : 0.5 → 0.0). (Right) The electrostatic decoupling leg (*λ* : 1.0 → 0.5). Darker regions correspond to high-probability basins representing more frequently sampled *λ*-conditioned hydration regimes. Source data are provided in Supplementary Data 1.

Hydration near PHE56 displays a different response (Fig. 10, bottom). During the electrostatic leg, the distribution remains concentrated at low coordination, CN≤ 1, consistent with comparatively labile hydration near this hydrophobic region. However, during the van der Waals leg, the distribution broadens toward higher CN as the ligand excluded volume is reduced and additional water configurations become accessible. This behavior is consistent with hydration near the W2/PHE56 region depending more strongly on geometric confinement than on direct directional interactions with the protein. The two profiles therefore resolve distinct local responses within the same buried pocket: persistent, polar-anchored hydration near TYR138 and a broader, more exchange-prone hydration distribution near PHE56.

As a complementary structural-localization check, we monitored the occupancy of the crystallographic W1 and W2 regions during the bound-state Lambda-ABF-OPES calculations (Fig. S32). The resulting traces show that sampled waters remain localized near the buried crystallographic hydration regions during both alchemical legs. Together, the site occupancies, continuous-CN distributions, and bridging analyses show that Lambda-ABF-OPES efficiently samples the persistent buried hydration motif of 1I05 while allowing distinct local solvent responses to electrostatic and steric decoupling, without applying restraints to individual water molecules.

The present benchmark should be interpreted as a validation of hydration recovery and exchange for systems with experimentally defined water-mediated motifs, rather than as a universal test of *de novo* hydration-network formation. Such systems provide a controlled structural reference for assessing whether Lambda-ABF-OPES can preserve, recover, and exchange buried or semi-buried waters during ligand decoupling. Continuous alchemical scaling is expected to be most effective when water entry is controlled primarily by the ligand steric or electrostatic footprint. When hydration is instead coupled to slow protein conformational gating, such as loop opening, side-chain rearrangement, or cryptic-pocket formation, additional orthogonal sampling of the relevant protein motion may be required.

### Remaining Complexes

#### 5I40

In addition to the TAF1(2)–ligand complex (PDB id: 5I1Q), we examined the BRD9–pyrrolopyridinone complex (PDB id: 5I40)^76^ to explore differences in binding-site hydration and ligand recognition between these bromodomain subtypes. Unlike TAF1(2), BRD9 exhibits variations in binding-cavity size, shape, and flexibility, as well as amino acid substitutions that influence ligand access and hydration networks. The ligand, 6-methyl-1,6-dihydro-7H-pyrrolo[2,3-c]pyridin-7-one, acts as an acetyl-lysine mimetic, forming two-point hydrogen bonds with ASN100 via the pyridone carbonyl and pyrrole nitrogen (Fig. 1D), while its aromatic core engages in *π*–*π* stacking with the TYR106 gatekeeper.^76^

Throughout the 100-ns conventional MD simulation, the overall geometry of the protein–ligand complex was preserved. In contrast, the hydration sites, including those bridging ASN100 and TYR106, exhibited pronounced dynamism. Density distribution plots reveal a broad spatial distribution of bridging water positions (Figs. S33–S36), indicating that the water molecules undergo transient solvent exchange rather than remaining localized as persistent structural anchors. A conserved water site has also been reported to participate in a tetracoordinated hydrogen-bonding network involving ASN95, TYR57, and MET65 based on static structural analysis.^118^ However, while the crystallographic configuration highlights potential hydration motifs, it does not necessarily reflect their persistence under dynamic conditions and was not persistently maintained in our conventional MD simulations.

Regarding binding-affinity calculations, the corresponding PMFs converged (Figs. S37–S38), and the cumulative CN of bridging water molecules evaluated during conventional MD (Fig. S39) is in good agreement with the CN profiles obtained from the van der Waals and electrostatic alchemical transformations (Figs. S40–S41). This correspondence supports the sampling of solvent exchange and rehydration during the alchemical calculations and shows that Lambda-ABF-OPES accommodates a dynamically reorganizing hydration shell in the BRD9 pocket.

To characterize the site-resolved solvent response associated with this dynamic hydration, we computed continuous hydration profiles for ASN100 and TYR106 (Fig. S61). For ASN100 (Fig. S61, top), the electrostatic leg reveals negligible hydration density. However, upon steric decoupling (*λ* → 0), a distinct hydration population emerges near CN≈ 1.5, consistent with resolvation of this recognition region as the ligand excluded volume is reduced. In contrast, TYR106 (Fig. S61, bottom) displays a broad, high-occupancy probability distribution across the transformation. Consistent with its location at the pocket entrance, this profile reflects a broad and heterogeneous solvent population that remains accessible over a wide range of ligand coupling states. The contrast between ligand-dependent hydration at ASN100 and the broader solvent response around TYR106 illustrates how distinct regions of the BRD9 pocket respond differently to ligand decoupling, with ligand recognition involving local solvent displacement rather than stabilization by a persistent bridging-water network.

#### Hsp90-related complexes

The N-terminal domain of Hsp90 is an ATP-binding chaperone module essential for the maturation and stability of many oncogenic proteins. Its binding pocket is characterized by a polar subsite flanked by a hydrophobic cavity. The two Hsp90–ligand complexes, 2XAB and 2XJG, differ by a single modification of the resorcinol capping group: the hydroxyl substituent in 2XAB is replaced by a methoxy group in 2XJG.^77^ Despite this minimal change, the crystallographic structures differ slightly in resolution, which may affect model construction and subsequent simulation. 2XAB was refined at 1.9 Å, providing somewhat greater confidence in side-chain orientations and ordered solvent placement than 2XJG, which was resolved at 2.25 Å (Table 1).

Initially, unresolved histidine-rich segments in disordered regions and in the binding site were built for both 2XAB and 2XJG and assigned as neutral *δ*-tautomers (HSD), consistent with their local environment and potential hydrogen-bond network. The simulations using these full models (i.e., with disordered regions) revealed that inclusion of the disordered segments had very different effects in the two complexes. For 2XAB, the loops had a minor impact on the computed binding free energy providing a good agreement with the experimental data (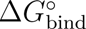 = −11.94 ± 1.01 kcal·mol*^−^*^1^ vs. ΔG_exp_ = −12.6 kcal·mol*^−^*^1^, Table 2), indicating that the full model is stable and can be used for accurate binding affinity evaluation. In contrast, for 2XJG, the disordered loops were highly mobile and introduced large-amplitude fluctuations that dominated the conformational ensemble, yielding a binding free energy approximately 16 kcal·mol*^−^*^1^ less favorable than that obtained with the truncated model (Table S10). This high intrinsic mobility makes the full 2XJG model unsuitable for reliable binding free–energy calculations. Based on these observations, truncated models (omitting the disordered segments) were then generated for both complexes and tested. Here, histidine protonation within the binding site was also carefully assigned, as misassignment can alter local electrostatic potentials, reorient ordered waters, and change hydrogen-bond networks, routinely affecting computed ΔG by several *k_B_T*. Protonation states were determined by visual inspection of potential inter- and intramolecular hydrogen-bonding networks: three neutral *ɛ*-tautomers HSE77, HSE189, HSE210, and one neutral *δ*-tautomer HSD154. The results showed that the truncated 2XJG model restored stability and gave physically meaningful binding energetics (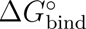 = −12.05 ± 0.45 kcal·mol*^−^*^1^ vs. ΔG_exp_ = −10.7 kcal·mol*^−^*^1^, Table 2, Figs. S47-S48, S58-S59), while for 2XAB, the truncation had a minor impact on the computed binding free energy (∼1.8 kcal·mol*^−^*^1^ less favorable), meaning that the full model including the disordered loops remains preferable for 2XAB to accurately capture the binding environment. All subsequent analyses were therefore performed using the truncated model for 2XJG and the full model for 2XAB, ensuring accurate representation of binding energetics without artefactual penalties.

To further characterize binding interactions, we analyzed density distributions of direct protein– ligand polar contacts and water-mediated polar contacts (Figs. S42–S45 for 2XAB and Figs. S51– S54 for 2XJG). For 2XAB, the distributions are generally more pronounced and numerous than in 2XJG, consistent with the capacity of the exposed ligand hydroxyl group to participate in water-mediated contacts. Several direct interaction patterns are shared between the two complexes, including contacts with PHE138 and MET98, and many direct polar contacts are additionally accompanied by bridging solvent molecules.

However, the organization and *λ*-dependent response of these solvent networks differ significantly between the two systems. While 2XAB accommodates a distinct hydration network, the methoxy group in 2XJG imposes a frustrated hydration landscape, where the competition between solvent displacement and retention hinders rapid equilibration. This observation aligns with Ge et al.^9^, who reported that the local solvent network in 2XJG is prone to kinetic trapping and requires extensive sampling to converge. Specifically, they found that even with advanced sampling strategies like BLUES^119^ (combining NCMC with classical MD) and GCMC,^120–123^ the 2XJG complex required millions of force evaluations to resolve the occupancy of these slow-exchanging sites.

We evaluated solvent-structure descriptors in both complexes and found that the principal re-hydration patterns were sampled during the alchemical transformations. Comparative plots of the continuous CN of bridging waters during conventional MD (Figs. S46 and S55) and its dependence on the alchemical coupling parameter *λ* are provided in Figs. S49–S50 for 2XAB and Figs. S56– S57 for 2XJG. Near the fully coupled state, the CN distributions obtained from the electrostatic and van der Waals calculations overlap with the range observed in conventional MD, supporting the structural consistency of the sampled hydration patterns. Convergence of the corresponding PMFs is shown in Figs. S47–S48 and Figs. S58–S59 for 2XAB and 2XJG, respectively.

To dissect the microscopic origins of these differential hydration patterns, we analyzed continuous hydration landscapes for the conserved residue GLY97 and the ligand-proximal residue THR184 (Figs. S62-S63), both of which participate in water-mediated contacts with ligand oxygen atoms (Fig. 1E-F). For GLY97, both complexes exhibit a persistent hydration population centered near CN≈1 across the alchemical transformations. This behavior indicates that the GLY97 hydration region remains populated despite the difference between the two ligands and contributes to the local polar network. The hydration response near THR184 is more ligand-dependent. In 2XJG (Fig. S63, bottom), replacement of the hydroxyl group by the bulkier methoxy substituent shifts the distribution toward a narrower, lower-coordination population near CN≈0.5. This reduction is consistent with steric restriction of water access by the methoxy group and illustrates how a localized ligand modification alters rehydration of the protein–ligand interface.

## Conclusion and Perspectives

Protein–ligand binding is not governed by a static network of pairwise interactions, but by the collective reorganization of the ligand, the receptor, and the surrounding solvent. Water is therefore not merely a structural component of the binding site or a passive background to molecular recognition. Its displacement, recruitment, and reorganization contribute directly to the thermodynamic balance between the bound and unbound states. Accurately describing this response remains one of the central challenges of absolute binding free-energy calculations, particularly in buried and semi-buried cavities where solvent exchange is coupled to ligand and pocket relaxation.

Here, we show that Lambda-ABF-OPES, combined with the polarizable AMOEBA force field, samples structural signatures of this coupled response across five water-mediated protein–ligand complexes while yielding absolute binding free energies in good agreement with experiment. The central result is not simply that water returns to a vacated binding site during ligand decoupling. Rather, the calculations show that the relevant hydration environments are sampled as part of the collective response of the ligand, receptor, and solvent to the progressive removal and restoration of ligand interactions. This occurs without restraining individual water molecules or introducing system-specific water collective variables.

The systems examined here further illustrate that there is no single microscopic signature of a correctly hydrated binding site. Some recognition motifs rely on strongly localized water-mediated contacts, whereas others are maintained through recurrent occupation by waters whose molecular identities change over time. In both cases, the relevant thermodynamic object is not an individual crystallographic water molecule, but the ensemble of solvent configurations compatible with the local protein–ligand environment. Accordingly, preservation of a particular water identity is, by itself, neither necessary nor sufficient for an accurate binding free-energy calculation. What matters is whether the simulation samples the solvent-mediated states that contribute to the equilibrium balance between coupled and decoupled configurations.

This perspective also clarifies the respective roles of the physical model and the sampling strategy. AMOEBA provides a polarizable energetic description of the interactions within the protein– ligand–water environment, whereas continuous alchemical sampling promotes relaxation of the steric, conformational, and solvent degrees of freedom coupled to ligand decoupling. The agreement obtained here should therefore be attributed to their combined action rather than to polarization, enhanced sampling, or hydration recovery alone. More generally, the results emphasize that improving the energetic model cannot compensate for incomplete configurational sampling, just as extensive sampling cannot correct an inadequate physical description of the molecular interactions.

The present benchmark was deliberately constructed from complexes containing experimentally resolved hydration motifs, thereby providing well-defined structural references against which solvent localization and reorganization could be assessed. The conclusions should therefore be interpreted within this domain. The study supports the application of the combined framework to the buried and semi-buried small-molecule complexes examined here, but it does not establish universal transferability to all forms of molecular recognition. In particular, when solvent penetration is controlled by slow loop opening, side-chain gating, domain motion, cryptic-pocket formation, or membrane reorganization, modulation of the ligand interactions along *λ* may not be sufficient to sample the relevant hydration transitions. Such systems may require additional enhanced sampling of the orthogonal conformational coordinates that control solvent access to the binding region.

Similar caution applies to shallow or highly solvent-exposed binding interfaces, flexible ligands, peptides, and protein–protein complexes. In these regimes, the principal challenge may shift from local pocket resolvation toward the sampling of multiple orientations, conformations, and binding modes. Extension of the framework to these systems will therefore require dedicated validation and, where necessary, restraint definitions or enhanced-sampling strategies adapted to their broader configurational ensembles.

The use of a fully polarizable force field remained computationally tractable within the GPU-accelerated Tinker-HP implementation, with convergence of the individual free-energy contributions generally reached within tens of nanoseconds per independent replicate for the systems studied here. Continued developments in adaptive sampling, automated restraint construction, and neural-network molecular models may further extend the accessible chemical and conformational space.^124–126^ Such advances should, however, preserve the statistical-mechanical rigor and explicit convergence controls required for quantitative free-energy prediction.

Beyond affinity estimation, an atomistic free-energy framework connects a thermodynamic observable to the molecular reorganizations from which it emerges. This connection is particularly valuable for water-mediated recognition, where apparently similar binding affinities may arise from different balances of direct interactions, solvent displacement, bridging hydration, and conformational relaxation. The broader implication of this work is therefore that binding-site water should neither be frozen into a predefined structural role nor treated only as an average occupancy. It should be sampled as an integral component of the molecular ensemble whose collective response determines binding.

## Acknowledgments

This work has received funding from the European Research Council (ERC) under the European Union’s Horizon 2020 research and innovation program (grant agreement No 810367), project EMC2 (JPP). We thank the Grand Équipement de Calcul Intensif (GENCI), Institut du Développement et des Ressources en Informatique (IDRIS), and Centre Informatique de l’Enseignement Supérieur (CINES), France, for their support in term of computational ressources through grant no. A0180716167 and A0150712052.

## Author contributions

MB, NA performed the computations. All authors analyzed the data. MB and NA wrote the paper with inputs from all other co-authors. LL and JPP designed and supervised the research.

## Data availability

The numerical source data underlying the graphs in Figs. 3–10 are provided with this paper as Supplementary Data 1 (`Supplementary_Data_1_Source_Data.zip`). The Tinker-HP and Colvars input files, initial and equilibrated structures, and supporting workflow files used in this study are openly available in the Zenodo repository at https://doi.org/10.5281/zenodo.22208958 ^127^, within the `Water_Networks_complexes` directory.

## Code availability

The custom Python scripts used to analyse the simulation data and generate the reported figures are provided in the `Code` directory of Supplementary Data 1 and are openly available through the GitHub repository at https://github.com/ansarinarjes/ABFE-L-ABF-OPES. A permanent, versioned snapshot of the code and associated workflow materials is archived in Zenodo at https://doi.org/10.5281/zenodo.22208958127. No access restrictions apply.

## Supporting Information Available

Supplementary Information: Computational details of the simulations and additional analyses (PDF).

Supplementary Data 1: Numerical source data underlying the graphs presented in Figs. 3–10. Plain-text files are organized by figure and panel. The accompanying README defines the columns, units, systems, alchemical legs, and replicate identifiers.

