## Supplementary Information for "Enhanced Sampling Enables Binding Site Water Reorganization in Polarizable Absolute Binding Free Energy Calculations"

### Source data and analysis code

The numerical source data underlying the graphs in main-text Figs. 3–10 are provided separately with this article as Supplementary Data 1 (`Supplementary_Data_1_Source_Data.zip`). The files are organized by main-text figure and panel, and the accompanying README defines all columns, units, systems, alchemical legs, and replicate identifiers. The custom Python scripts associated with these analyses are provided in the `Code` directory of Supplementary Data 1. The code and associated workflow materials are also available through https://github.com/ansarinarjes/ABFE-L-ABF-OPES, with a permanent snapshot archived at https://doi.org/10.5281/zenodo.22208958.

**Table S1. Summary of cubic simulation box side lengths, total number of water molecules, and ion composition for each simulated protein–ligand complex.**

| PDB id | Box side length (Å) | No. of Water Molecules | $Na^+$ ions | $Cl^-$ ions |
|---|---|---|---|---|
| 5I1Q | 84.830 | 19738 | 68 | 56 |
| 1I05 | 80.141 | 16250 | 59 | 46 |
| 2XJG | 93.806 | 26416 | 90 | 75 |
| 2XAB | 82.639 | 17753 | 65 | 50 |
| 5I40 | 72.090 | 12057 | 34 | 39 |

All systems were prepared in cubic condensed-phase boxes with a minimum solute–box boundary distance of 15 Å. Simulations used the AMOEBA-BIO-2018 polarizable force-field, which combines the AMOEBA 2013 protein parameters with the AMOEBA 2018 nucleic-acid parameters [1,2]. Water, [3] sodium and chloride ions were assigned the corresponding standard AMOEBA parameters.

### RMSF Profiles and DBC Anchor Atom Selection

The Root Mean Square Fluctuation (RMSF) profiles provide crucial insights into the positional flexibility of the protein backbones during the molecular dynamics simulations. Figure S1 shows the RMSF of all $C_\alpha$ atoms of the five studied protein systems (panels a–e) alongside their corresponding structural binding pockets.

The $C_\alpha$ atoms selected as Anchor atoms for the DBC mapping are explicitly highlighted with red markers. As detailed in the main text, these specific anchor atoms were filtered based on a strict spatial cutoff within 6 Å of the bound ligand in the initial frame. Crucially, as shown across all five simulated systems in Figure S1, every single selected DBC anchor atom consistently maintains an RMSF value below the threshold of 0.7 Å (denoted by the horizontal red dashed line), indicating exceptionally low thermal fluctuation and high structural rigidity within the binding core. The 3D structural panels to the right of each plot illustrate the spatial distribution of these rigid DBC anchor atoms (represented as yellow spheres) surrounding their respective ligands within the binding pocket.

**Table S2. System-specific DBC and center-of-mass restraint parameters used in the absolute binding free-energy calculations. $R_{\mathrm{DBC}}^{\mathrm{site}}$ is the flat-bottom DBC upper wall used in the bound complex; $R_{\mathrm{DBC}}^{\mathrm{gas}}$ is the corresponding DBC upper wall used in the gas-phase restraint-release calculation; and $R_{\mathrm{COM}}$ is the spherical center-of-mass restraint radius entering the standard-state volume correction. All radii are reported in Å and all force constants in kcal mol$^{-1}$ Å$^{-2}$.**

| PDB | $R_{\mathrm{DBC}}^{\mathrm{site}}$ | $R_{\mathrm{DBC}}^{\mathrm{gas}}$ | $R_{\mathrm{COM}}$ | $k_{\mathrm{DBC}}$ | $k_{\mathrm{COM}}$ |
|---|---|---|---|---|---|
| 5I1Q | 0.800 | 0.800 | 1.800 | 100.0 | 100.0 |
| 1I05 | 0.700 | 0.700 | 1.700 | 100.0 | 100.0 |
| 5I40 | 0.800 | 0.800 | 1.800 | 100.0 | 100.0 |
| 2XAB | 1.200 | 1.200 | 1.200 | 100.0 | 100.0 |
| 2XJG | 0.700 | 0.700 | 1.700 | 100.0 | 100.0 |

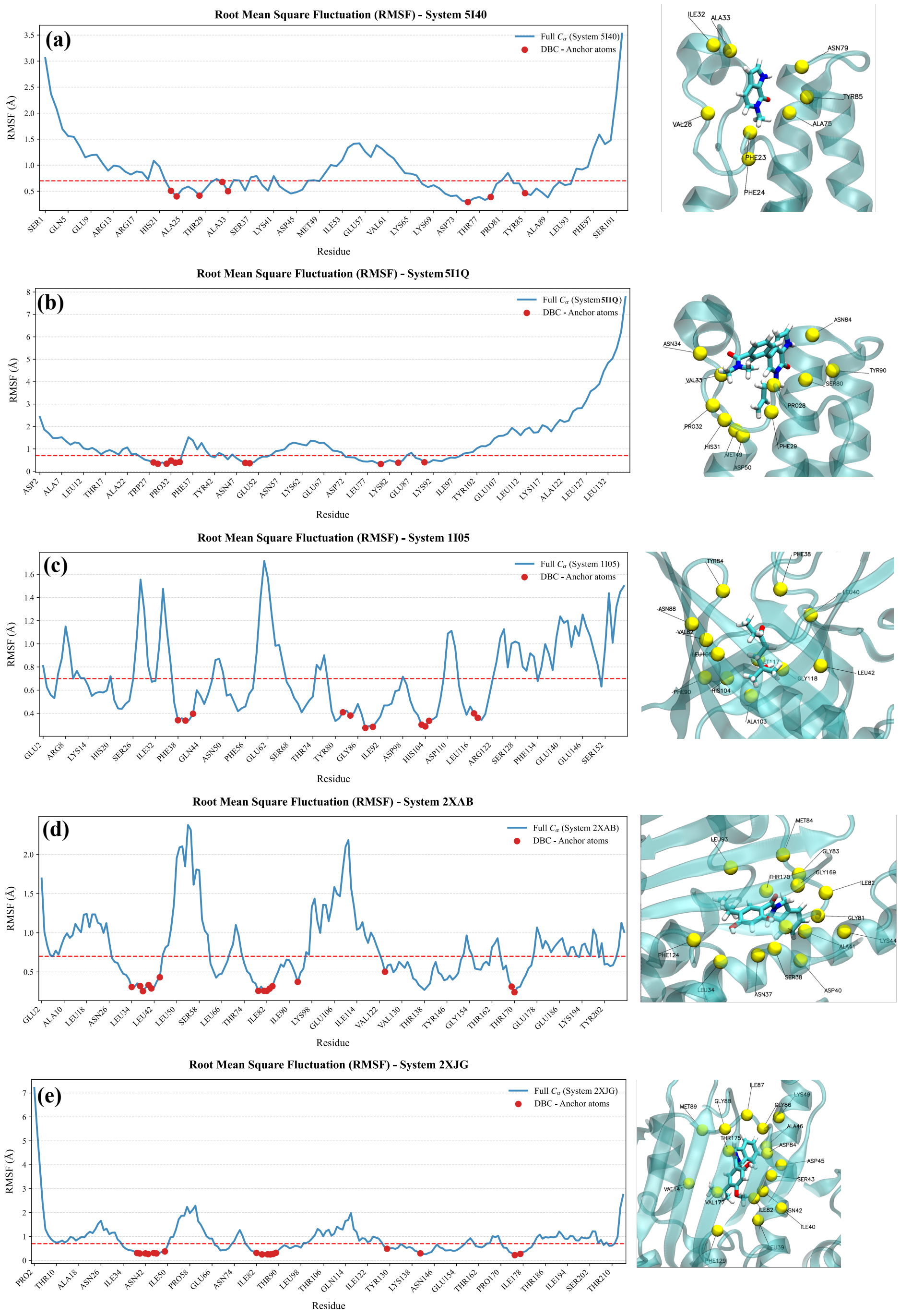


Figure S1. Root Mean Square Fluctuation (RMSF) profiles of $C_\alpha$ atoms across the five studied protein systems: (a) System 5I40, (b) System 5I1Q , (c) System 1I05, (d) System 2XAB, and (e) System 2XJG. The continuous line represents the full $C_\alpha$ trajectory, while red dots isolate the selected DBC anchor atoms located within 6 Å of the ligand. The horizontal red dashed line marks the 0.7 Å structural stability threshold. Right panels show the corresponding 3D visualization of the binding pockets with the localized anchor atoms displayed as spheres.

#### Example of Colvars File for $\Delta G^{*}_{\text{site}}$ Contribution Evaluation

```
colvarsTrajFrequency 1000
colvarsRestartFrequency 1000

colvar {
  name l
  extendedLagrangian on
  extendedLangevinDamping 1000
  extendedtemp 300
  extendedmass  150000

  lowerBoundary 0.5 # to evaluate electrostatic contribution
  upperBoundary 1.0 # to evaluate van der Waals, then lowerBoundary 0.0/ upperBoundary 0.5
  reflectingLowerBoundary
  reflectingUpperBoundary
  width 0.01
  alchLambda {
  }
  outputTotalForce
  outputAppliedforce
  subtractappliedforce
}


abf {
  colvars l
  fullSamples 10000
  historyFreq 1000
  shared
  sharedFreq 1000
}

opes_metad {
  name opes_l
  colvars l
  newHillFrequency 300
  barrier 2
  adaptiveSigma on
  neighborList on
  printTrajectoryFrequency 1000
  pmf on
```

```
  pmfColvars l
  pmfHistoryFrequency 1000
  outputEnergy on
  explore on
}

colvar {
    # A "distance to bound configuration" (DBC) coordinate for ligand binding restraints

    name DBC

    rmsd {
        # Reference coordinates (for ligand RMSD computation)
         refPositionsFile complex_ref.xyz # taken as the last frame of conventional MD in xyz-coordinates
        # refPositionsCol O
        # refPositionsColValue 1
            atoms {
            # Define ligand atoms used for RMSD calculation
            # "auto-updating" keyword updates atom IDs when applying cfg or changing
 molecule
            # auto-updating selection: "serial 2300 2301 2302 2303 2305 2306 2308 2309

2311 2313 2315 2316 2318 2319 2322 2327 2330 2331 2333 2335 2336"
            atomNumbers 2300 2301 2302 2303 2305 2306 2308 2309 2311 2313 2315 2316

2318 2319 2322 2327 2330 2331 2333 2335 2336

            # Moving frame of reference is defined below
            centerToReference yes
            rotateToReference yes
            fittingGroup {
                # Define binding site atoms used for fitting
            # "auto-updating" keyword updates atom IDs when applying cfg or changing molecule
                        # auto-updating selection: "serial 465 489 523 540 557 571 836 850 1356 1419 1501"
            atomNumbers 465 489 523 540 557 571 836 850 1356 1419 1501
}
           refPositionsFile complex_ref.xyz # taken as the last frame of conventional MD in xyz-coordinates
        # refPositionsCol O
        # refPositionsColValue 1

        }
```

```
    }
}
harmonicWalls {
colvars DBC
upperWalls 0.8
forceConstant 100.0
}
colvar {
  name cnuml
  coordNum {
    group1 {
      atomnumbersRange  2296-2341  # ligand
      }
    group2 {
      atomnumbersRange  61556-61679  #all ions
      }
  }
}

scriptedColvarForces
sourceTclFile script.tcl
```

#### Example of Colvars File for $\Delta G^{*}_{\mathbf{bulk}}$ Contribution Evaluation

```
colvarsTrajFrequency 1000
colvarsRestartFrequency 1000

colvar {
  name l
  extendedLagrangian on
  extendedLangevinDamping 1000
  extendedtemp 300
  extendedmass  150000

  lowerBoundary 0.5 # to evaluate electrostatic contribution
  upperBoundary 1.0 # to evaluate van der Waals, then lowerBoundary 0.0/ upperBoundary 0.5
  reflectingLowerBoundary
  reflectingUpperBoundary
  width 0.01
  alchLambda {
```

```
  }
  outputTotalForce
  outputAppliedforce
  subtractappliedforce
}


abf {
  colvars l
  fullSamples 10000
  historyFreq 1000
  shared
  sharedFreq 1000
}

opes_metad {
  name opes_l
  colvars l
  newHillFrequency 300
  barrier 2
  adaptiveSigma on
  neighborList on
  printTrajectoryFrequency 1000
  pmf on
  pmfColvars l
  pmfHistoryFrequency 1000
  outputEnergy on
  explore on
}


colvar {
  name cnuml
#  width 0.1
  coordNum {
    group1 {
      atomnumbersRange  1-46  # ligand
      }
    group2 {
      atomnumbersRange 59261-59372  #ions
}
```

```
  }
}

scriptedColvarForces
sourceTclFile script.tcl
```

##### script.tcl mentioned in *.colvars code

```
proc calc_colvar_forces { ts } {
  #  puts "Running calc_colvar_forces at timestep $ts"
  set k 10

  set l [cv colvar l value]


  set c [cv colvar cnuml value]
  if { $l < 0.95 } {
    set Fc [expr -$k*($c)]
    cv colvar cnuml addforce $Fc
  }
  if { $l > 0.05 } {
    set Fc [expr -$k*($c)]
    cv colvar cnuml addforce $Fc
  }
}
```

#### Example of Colvars File for $\triangle G_{DBC}$ Contribution Evaluation

```
colvarsTrajFrequency 1000
colvarsRestartFrequency 1000

colvar {
    name DBC
```

```
    rmsd {
        refPositionsFile dbc_gas_ref.xyz
        atoms {
            atomNumbers 5 7 11 12 14 15 16 17 18 21 22 28 30 31 34 35 37 39 40 41
            # same atoms as used in the bound contribution calculations

            # Moving frame of reference is defined below
            centerToReference no
            rotateToReference no
        }
    }
}

colvar {
    name d
    # distance of  centre of mass to the centre of the refrence
    distance {
        group1 {atomNumbers 5 7 11 12 14 15 16 17 18 21 22 28 30 31 34 35 37 39 40 41}
        # same atoms as used in the bound contribution calculations
        group2 {
            dummyAtom (0.03522,-0.53846,0.13517)
        }
    }
}

harmonicWalls {
    # COM Restraints
    colvars d
    upperWalls 1.8 # defined DBC + 1
    forceConstant 100
}

harmonicwalls {
```

```
    colvars DBC
    upperWalls 0.8 # same as the defined DBC
    forceConstant 100.0
    decoupling on
    lambdaExponent 5.0
    targetnumstages 20
    targetnumsteps 1000000
    outputenergy
}
```

### PMF Convergence Analysis

Convergence of the potential of mean force (PMF) was assessed from generalized from four walkers history-based PMF outputs (**.all.hist.pmf*) generated during simulation. For each replica, the free energy value at $\lambda$ =0.5 was extracted at every iteration. This produced a time series of $\Delta$G values that reflect how the estimated free energy evolves with simulation length. For each replica, $\Delta$G values at the selected coordinate were plotted as a function of iteration (frames), yielding a convergence trajectory.

# 5I1Q

#### Site-resolved hydration-distance analysis for the 5I1Q conventional MD trajectory

To address whether the water-density and occupancy maps correspond to persistent protein–water–ligand contacts, we performed a site-resolved distance analysis for the 5I1Q conventional MD trajectory. The W1 and W2 hydration sites were defined from the crystallographic water oxygen positions WAT1875 and WAT1867, respectively. For each trajectory frame, the protein was aligned to the crystallographic reference using C$\alpha$ atoms, and the nearest water oxygen to each crystallographic site center was identified. Two occupancy definitions were used: a strict crystallographic-core cutoff of $2.0$ Å and a broader exchange-shell cutoff of $3.5$ Å.

For each assigned site water, we computed direct PBC-corrected distances to the corresponding protein and ligand polar atoms. For W1, the protein-side anchor was TYR1540 OH. Because the original crystallographic ligand O1 geometry was not preserved during the 80–100 ns interval, we additionally monitored distances to the ligand O2 and N3 atoms, which form the nearest ligand polar region during this interval. For W2, we monitored the distance to the closest protein anchor among HSD1530 O, PRO1531 O, and VAL1532 N, together with distances to ligand O2 and N3. In parallel, we decomposed the oxygen-based O-bridging coordination number into W1, W2, and remaining-water contributions using the same $4.0$ Å cutoff as in the main CN analysis.

The resulting analysis shows that W1/WAT1875 remains tightly occupied until approximately 93 ns, followed by a marked depletion of the crystallographic site. The W1 strict-core occupancy decreases from $99.2\%$ during 80–90 ns and $99.3\%$ during 90–93 ns to $13.8\%$ during 93–95 ns and $8.0\%$ during 95–100 ns. Water-identity tracking shows that this depletion is not a replacement event: the original WAT1875 water remains the nearest water when the site is occupied, whereas stable replacement-water occupancy is negligible. The W2/WAT1867 site shows a different behavior. The original WAT1867 water does not persist, but the site remains recurrently hydrated by exchange waters, with transient depletion around 93–95 ns followed by rehydration during 95–100 ns.

The ligand-distance analysis also reveals that the W1 site does not preserve the original crystallographic O1-mediated bridge geometry in the analyzed interval. Instead, the W1 water remains mainly protein-anchored through TYR1540 OH, while its nearest ligand polar partners are O2 and N3. Therefore, W1 contributes to the broader O-bridging CN through a redistributed polar environment rather than through a persistent O1-mediated crystallographic bridge. Consistently, the W1-specific O-bridging CN contribution decreases strongly after W1 depletion, whereas the remaining global CN is carried mainly by other transient waters. W2 does not act as a direct compensatory replacement for W1 in the O-bridging metric.

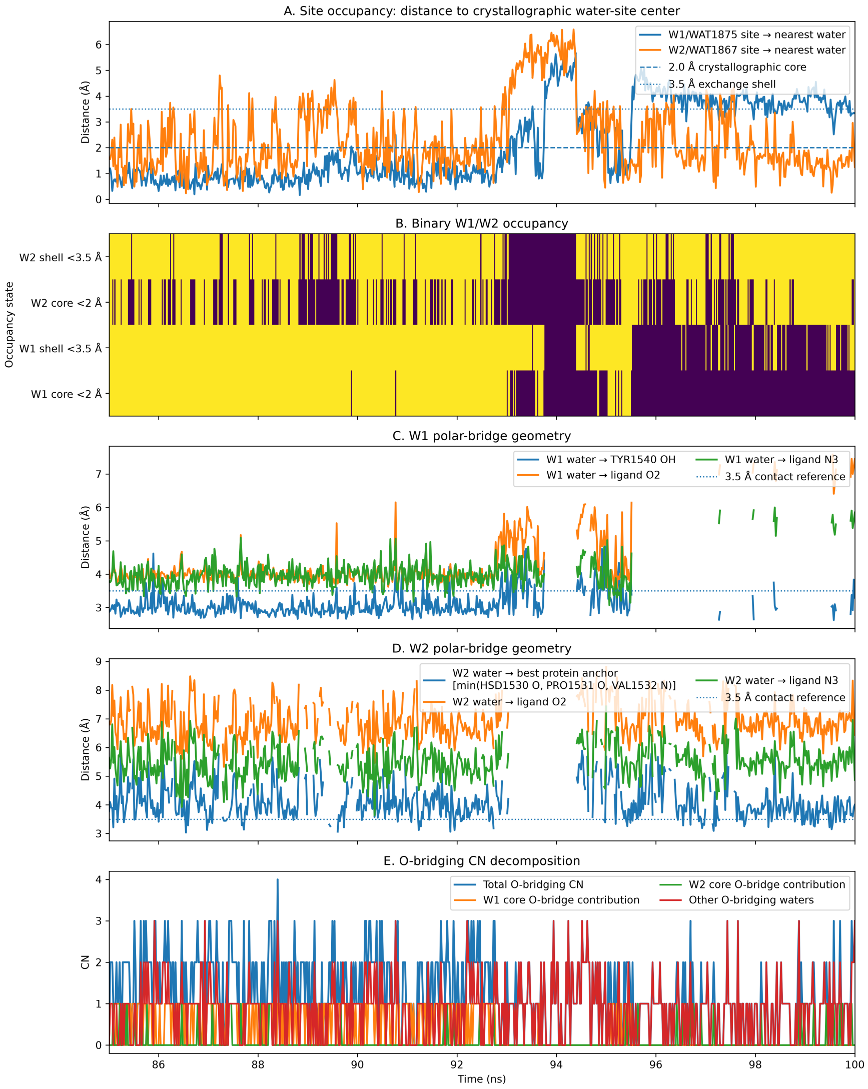


Figure S2. Site-resolved W1/W2 hydration-distance analysis for the 5I1Q conventional MD trajectory. W1 and W2 were defined from the crystallographic water oxygen positions WAT1875 and WAT1867, respectively. (A) Distance between each crystallographic water-site center and the nearest water oxygen over the 85–100 ns interval. The dashed and dotted horizontal lines indicate the 2.0 Å crystallographic-core cutoff and the 3.5 Å exchange-shell cutoff, respectively. (B) Binary occupancy map for the W1 and W2 core and exchange-shell definitions. The W1 site remains occupied until approximately 93 ns and then becomes depleted, whereas W2 behaves as an exchange-hydrated site with a transient dehydration interval followed by rehydration. (C) Polar-distance analysis for the W1-assigned water. The W1 water remains primarily anchored on the protein side through TYR1540 OH before depletion, while the nearest ligand polar contacts involve the pyrrolopyridone carbonyl oxygen O2 and the adjacent ring nitrogen N3. (D) Polar-distance analysis for the W2-assigned water. The W2 water was compared with the nearest protein anchor among HSD1530 O, PRO1531 O, and VAL1532 N, and with ligand O2/N3. The discontinuities in panels C and D arise because distances are reported only when the corresponding crystallographic site is occupied within the 3.5 Å exchange shell. (E) Decomposition of the oxygen-based O-bridging coordination number into W1, W2, and other-water contributions. The W1-specific contribution decreases after W1 depletion, while the residual CN is mainly carried by other transient bridging waters.

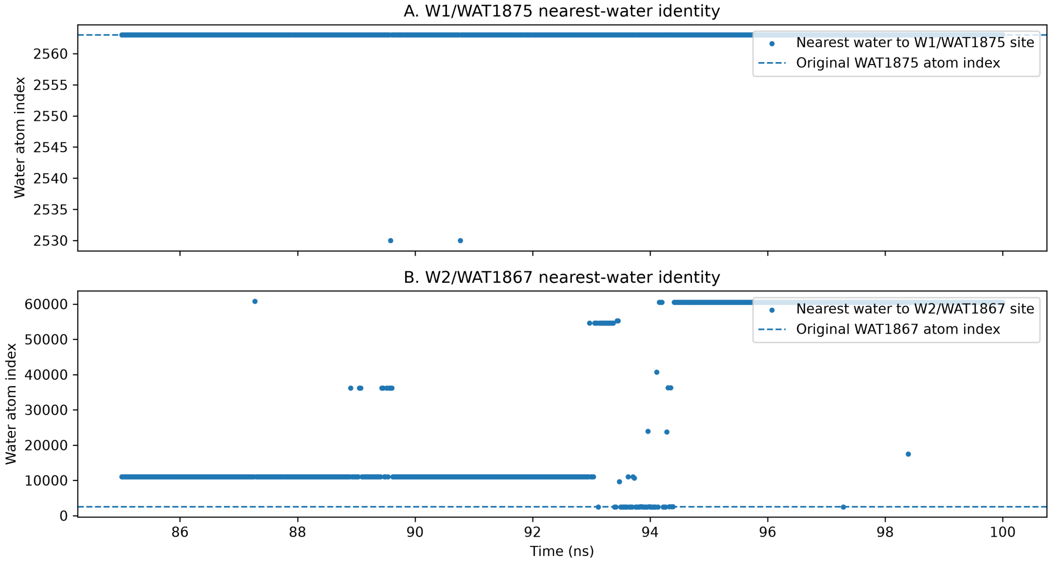


Figure S3. Water-identity tracking for the W1/WAT1875 and W2/WAT1867 crystallographic hydration sites in the 5I1Q conventional MD trajectory. Each point reports the atom index of the nearest water oxygen to the corresponding crystallographic site center. The dashed horizontal line indicates the original crystallographic water atom index. For W1, the original WAT1875 water remains the nearest water when the site is occupied, showing that the late-time depletion is not a stable replacement event. For W2, the original WAT1867 water is not retained; the site is instead populated by exchange waters.

The labels O/O, N/O, O/N, and N/N denote the heavy-atom element classes used for contact classification, with the first element corresponding to the protein atom and the second to the ligand atom. Residue labels in the figures are used as compact notation. The corresponding atom-level assignments, including protein atom name and backbone versus side-chain origin, are reported in Table S3

**Table S3. Atom-level direct and water-mediated polar contacts for 5i1q. Motif labels are reported as protein-element/ligand-element. Direct contacts use a 4.0 Å heavy-atom cutoff. Water-mediated contacts require both ligand–water and water–protein distances to be within 4.0 Å, and ligand–protein proximity ≤ 6.0 Å.**

| contact type | motif | contact label | atom origin | occ. fraction | mean events | direct dist Å | lig–wat dist Å | wat–prot dist Å | total path Å | unique water O |
|---|---|---|---|---|---|---|---|---|---|---|
| direct | N/O | 67C1701:O2–ASN1583:ND2 (side chain) | side chain | 0.916 | 1.00 | 3.80 | | | | |
| direct | O/N | 67C1701:N2–VAL1532:O (backbone) | backbone | 0.421 | 1.00 | 3.66 | | | | |
| direct | N/O | 67C1701:O1–ASN1533:N (backbone) | backbone | 0.315 | 1.00 | 3.40 | | | | |
| direct | N/N | 67C1701:N1–ASN1533:N (backbone) | backbone | 0.308 | 1.00 | 3.39 | | | | |
| direct | N/O | 67C1701:O1–HSD1530:NE2 (side chain) | side chain | 0.301 | 1.00 | 3.25 | | | | |
| direct | N/N | 67C1701:N1–HSD1530:NE2 (side chain) | side chain | 0.296 | 1.00 | 3.26 | | | | |
| direct | O/N | 67C1701:N3–VAL1532:O (backbone) | backbone | 0.157 | 1.00 | 3.82 | | | | |
| direct | O/O | 67C1701:O2–VAL1532:O (backbone) | backbone | 0.095 | 1.00 | 3.76 | | | | |
| direct | N/N | 67C1701:N1–HSD1530:ND1 (side chain) | side chain | 0.056 | 1.00 | 3.67 | | | | |
| direct | N/O | 67C1701:O1–HSD1530:ND1 (side chain) | side chain | 0.052 | 1.00 | 3.68 | | | | |
| direct | N/N | 67C1701:N1–PRO1531:N (backbone) | backbone | 0.033 | 1.00 | 3.69 | | | | |
| direct | N/O | 67C1701:O1–PRO1531:N (backbone) | backbone | 0.032 | 1.00 | 3.68 | | | | |
| direct | N/O | 67C1701:O1–VAL1532:N (backbone) | backbone | 0.031 | 1.00 | 3.80 | | | | |
| direct | N/N | 67C1701:N1–VAL1532:N (backbone) | backbone | 0.029 | 1.00 | 3.79 | | | | |

*Continued on next page*

#### Table S3. Atom-level direct and water-mediated polar contacts for 5i1q (continued)

| contact type | motif | contact label | atom origin | occ. fraction | mean events | direct dist Å | lig–wat dist Å | wat–prot dist Å | total path Å | unique water O |
|---|---|---|---|---|---|---|---|---|---|---|
| direct | O/N | 67C1701:N3–PHE1528:O (backbone) | backbone | 0.022 | 1.00 | 3.87 | | | | |
| direct | O/O | 67C1701:O1–HSD1530:O (backbone) | backbone | 0.021 | 1.00 | 3.39 | | | | |
| direct | O/N | 67C1701:N1–HSD1530:O (backbone) | backbone | 0.021 | 1.00 | 3.36 | | | | |
| direct | N/O | 67C1701:O1–ASN1533:ND2 (side chain) | side chain | 0.017 | 1.00 | 3.58 | | | | |
| direct | N/N | 67C1701:N1–ASN1533:ND2 (side chain) | side chain | 0.016 | 1.00 | 3.56 | | | | |
| direct | O/N | 67C1701:N3–TYR1540:OH (side chain) | side chain | 0.014 | 1.00 | 3.77 | | | | |
| direct | N/N | 67C1701:N3–PHE1528:N (backbone) | backbone | 0.011 | 1.00 | 3.94 | | | | |
| direct | O/O | 67C1701:O1–PRO1531:O (backbone) | backbone | 0.011 | 1.00 | 3.70 | | | | |
| water-mediated | N/N | 67C1701:N1–W(O)–HSD1530:NE2 (side chain) | side chain | 0.612 | 1.68 | | 3.34 | 3.22 | 6.56 | 1045 |
| water-mediated | N/O | 67C1701:O1–W(O)–HSD1530:NE2 (side chain) | side chain | 0.606 | 1.72 | | 3.35 | 3.22 | 6.57 | 1064 |
| water-mediated | O/N | 67C1701:N3–W(O)–TYR1540:OH (side chain) | side chain | 0.601 | 1.00 | | 3.72 | 3.00 | 6.73 | 3 |
| water-mediated | O/O | 67C1701:O2–W(O)–TYR1540:OH (side chain) | side chain | 0.587 | 1.00 | | 3.86 | 3.00 | 6.85 | 1 |
| water-mediated | O/N | 67C1701:N3–W(O)–TYR1540:O (backbone) | backbone | 0.567 | 1.00 | | 3.72 | 3.51 | 7.23 | 3 |
| water-mediated | O/O | 67C1701:O2–W(O)–TYR1540:O (backbone) | backbone | 0.477 | 1.00 | | 3.86 | 3.48 | 7.34 | 1 |
| water-mediated | O/O | 67C1701:O2–W(O)–ASN1583:O (backbone) | backbone | 0.341 | 1.00 | | 3.86 | 3.71 | 7.57 | 1 |
| water-mediated | N/O | 67C1701:O2–W(O)–ASN1583:ND2 (side chain) | side chain | 0.313 | 1.00 | | 3.86 | 3.76 | 7.62 | 1 |
| water-mediated | O/N | 67C1701:N3–W(O)–ASN1583:O (backbone) | backbone | 0.273 | 1.00 | | 3.75 | 3.73 | 7.48 | 1 |
| water-mediated | N/O | 67C1701:O1–W(O)–ASN1533:ND2 (side chain) | side chain | 0.270 | 1.24 | | 3.37 | 3.50 | 6.87 | 933 |

*Continued on next page*

#### Table S3. Atom-level direct and water-mediated polar contacts for 5i1q (continued)

| contact type | motif | contact label | atom origin | occ. fraction | mean events | direct dist Å | lig–wat dist Å | wat–prot dist Å | total path Å | unique water O |
|---|---|---|---|---|---|---|---|---|---|---|
| water-mediated | N/N | 67C1701:N1–W(O)–ASN1533:ND2 (side chain) | side chain | 0.267 | 1.25 | | 3.35 | 3.51 | 6.85 | 914 |
| water-mediated | O/N | 67C1701:N1–W(O)–PRO1531:O (backbone) | backbone | 0.267 | 1.06 | | 3.25 | 3.10 | 6.35 | 335 |
| water-mediated | O/O | 67C1701:O1–W(O)–PRO1531:O (backbone) | backbone | 0.264 | 1.06 | | 3.28 | 3.11 | 6.39 | 337 |
| water-mediated | N/N | 67C1701:N1–W(O)–PRO1531:N (backbone) | backbone | 0.245 | 1.00 | | 3.30 | 3.65 | 6.95 | 242 |
| water-mediated | N/N | 67C1701:N3–W(O)–ASN1583:ND2 (side chain) | side chain | 0.243 | 1.00 | | 3.78 | 3.80 | 7.58 | 1 |
| water-mediated | N/O | 67C1701:O1–W(O)–PRO1531:N (backbone) | backbone | 0.238 | 1.01 | | 3.33 | 3.65 | 6.98 | 236 |
| water-mediated | N/O | 67C1701:O1–W(O)–HSD1530:ND1 (side chain) | side chain | 0.225 | 1.17 | | 3.29 | 3.69 | 6.98 | 638 |
| water-mediated | N/N | 67C1701:N1–W(O)–HSD1530:ND1 (side chain) | side chain | 0.212 | 1.19 | | 3.29 | 3.70 | 6.99 | 631 |
| water-mediated | N/N | 67C1701:N1–W(O)–ASN1533:N (backbone) | backbone | 0.155 | 1.01 | | 3.30 | 3.58 | 6.89 | 398 |
| water-mediated | N/O | 67C1701:O1–W(O)–ASN1533:N (backbone) | backbone | 0.154 | 1.01 | | 3.31 | 3.58 | 6.90 | 390 |
| water-mediated | O/O | 67C1701:O2–W(O)–SER1579:OG (side chain) | side chain | 0.100 | 1.00 | | 3.87 | 3.78 | 7.64 | 1 |
| water-mediated | N/O | 67C1701:O1–W(O)–HSD1530:N (backbone) | backbone | 0.095 | 1.07 | | 3.40 | 3.16 | 6.56 | 59 |
| water-mediated | N/N | 67C1701:N1–W(O)–HSD1530:N (backbone) | backbone | 0.090 | 1.06 | | 3.41 | 3.16 | 6.57 | 67 |
| water-mediated | O/O | 67C1701:O1–W(O)–ASN1533:O (backbone) | backbone | 0.090 | 1.12 | | 3.37 | 3.61 | 6.99 | 354 |
| water-mediated | N/N | 67C1701:N1–W(O)–VAL1532:N (backbone) | backbone | 0.088 | 1.00 | | 3.52 | 3.68 | 7.21 | 20 |
| water-mediated | N/O | 67C1701:O1–W(O)–VAL1532:N (backbone) | backbone | 0.086 | 1.00 | | 3.53 | 3.69 | 7.22 | 24 |
| water-mediated | O/N | 67C1701:N1–W(O)–ASN1533:O (backbone) | backbone | 0.074 | 1.15 | | 3.34 | 3.58 | 6.92 | 316 |
| water-mediated | O/O | 67C1701:O1–W(O)–ASN1533:OD1 (side chain) | side chain | 0.072 | 1.05 | | 3.33 | 3.60 | 6.93 | 275 |

*Continued on next page*

#### Table S3. Atom-level direct and water-mediated polar contacts for 5i1q (continued)

| contact type | motif | contact label | atom origin | occ. fraction | mean events | direct dist Å | lig–wat dist Å | wat–prot dist Å | total path Å | unique water O |
|---|---|---|---|---|---|---|---|---|---|---|
| water-mediated | O/N | 67C1701:N1–W(O)–ASN1533:OD1 (side chain) | side chain | 0.068 | 1.07 | | 3.33 | 3.60 | 6.93 | 270 |
| water-mediated | O/N | 67C1701:N1–W(O)–PHE1536:O (backbone) | backbone | 0.066 | 1.24 | | 3.34 | 3.43 | 6.77 | 253 |
| water-mediated | N/O | 67C1701:O1–W(O)–HSD1529:NE2 (side chain) | side chain | 0.062 | 1.22 | | 3.31 | 3.10 | 6.41 | 103 |
| water-mediated | O/O | 67C1701:O1–W(O)–PHE1536:O (backbone) | backbone | 0.060 | 1.26 | | 3.37 | 3.43 | 6.79 | 250 |
| water-mediated | N/N | 67C1701:N1–W(O)–HSD1529:NE2 (side chain) | side chain | 0.053 | 1.18 | | 3.36 | 3.07 | 6.44 | 90 |
| water-mediated | O/N | 67C1701:N3–W(O)–SER1579:OG (side chain) | side chain | 0.038 | 1.00 | | 3.70 | 3.73 | 7.43 | 1 |
| water-mediated | O/N | 67C1701:N1–W(O)–VAL1532:O (backbone) | backbone | 0.033 | 1.00 | | 3.45 | 3.18 | 6.63 | 11 |
| water-mediated | O/O | 67C1701:O1–W(O)–VAL1532:O (backbone) | backbone | 0.029 | 1.00 | | 3.44 | 3.25 | 6.69 | 9 |
| water-mediated | O/N | 67C1701:N2–W(O)–TYR1540:OH (side chain) | side chain | 0.022 | 1.00 | | 3.92 | 3.03 | 6.95 | 1 |
| water-mediated | O/N | 67C1701:N2–W(O)–TYR1540:O (backbone) | backbone | 0.019 | 1.00 | | 3.91 | 3.51 | 7.42 | 1 |
| water-mediated | O/O | 67C1701:O1–W(O)–HSD1530:O (backbone) | backbone | 0.019 | 1.44 | | 3.41 | 3.36 | 6.77 | 69 |
| water-mediated | O/N | 67C1701:N1–W(O)–HSD1530:O (backbone) | backbone | 0.017 | 1.45 | | 3.40 | 3.39 | 6.79 | 63 |

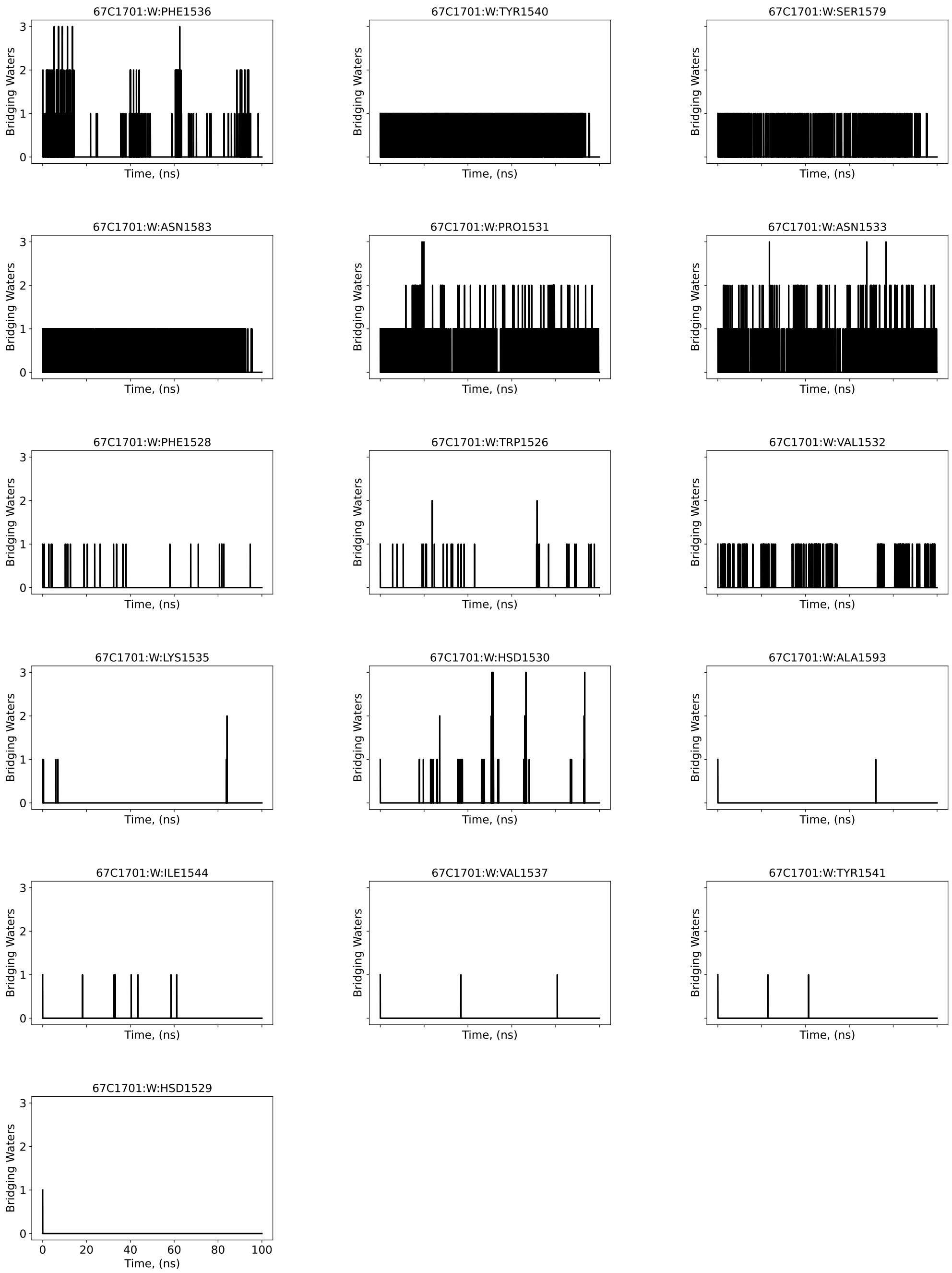


Figure S4. 4-Å -coordination number of O-bridging water molecules that mediate individual oxygen protein-ligand atom pairs along 100-ns conventional MD for the TAF1(2) complex (PDB: 5I1Q).

#### Water-site tracking around ligand generalized dipole

Water dynamics near the ligand were analyzed within VMD.[4] All ligand atoms and all water oxygen atoms within 20 Å of the ligand were initially selected. For each trajectory frame, the

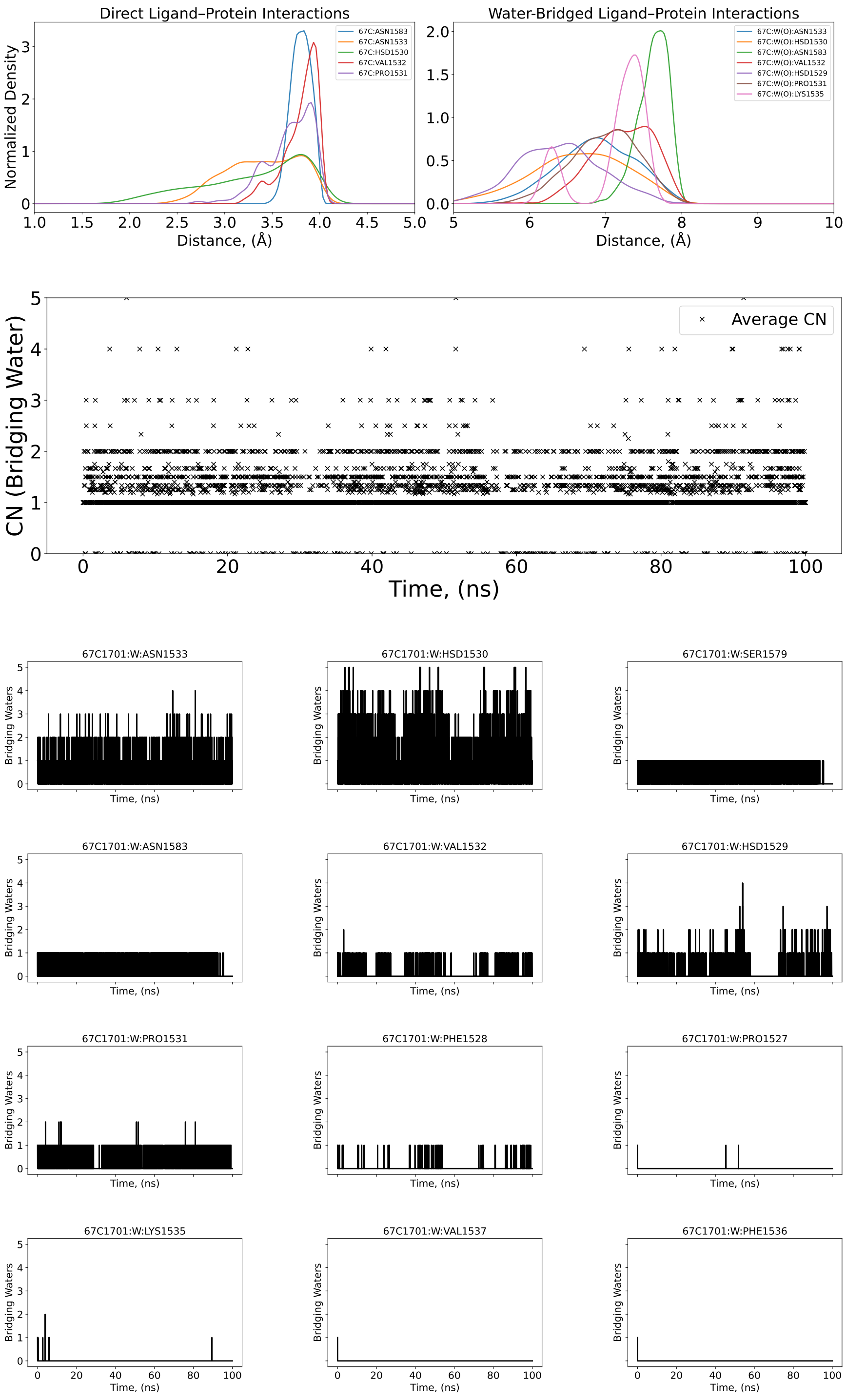


Figure S5. 4-Å–density distribution, averaged and individual coordination number of bridging oxygen water molecules (N/O-bridging) that mediate individual nitrogen atoms of the protein and oxygen atoms of the ligand along a 100-ns conventional MD for the TAF1(2) complex (PDB: 5I1Q).

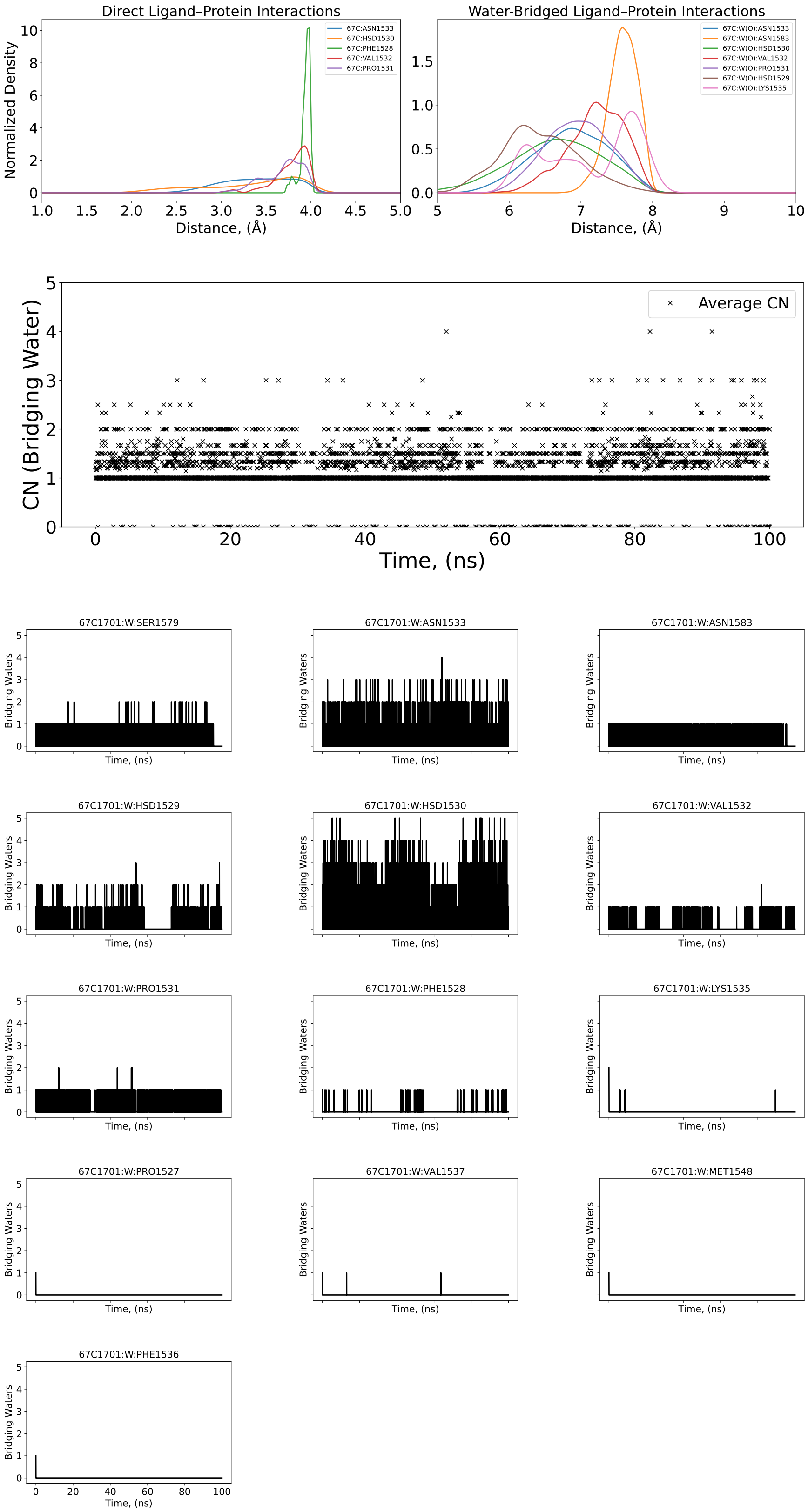


Figure S6. 4-Å–density distribution, averaged and individual coordination number of bridging oxygen water molecules (N-bridging) that mediate individual nitrogen atoms of the protein and the ligand along a 100-ns conventional MD for the TAF1(2) complex (PDB: 5I1Q).

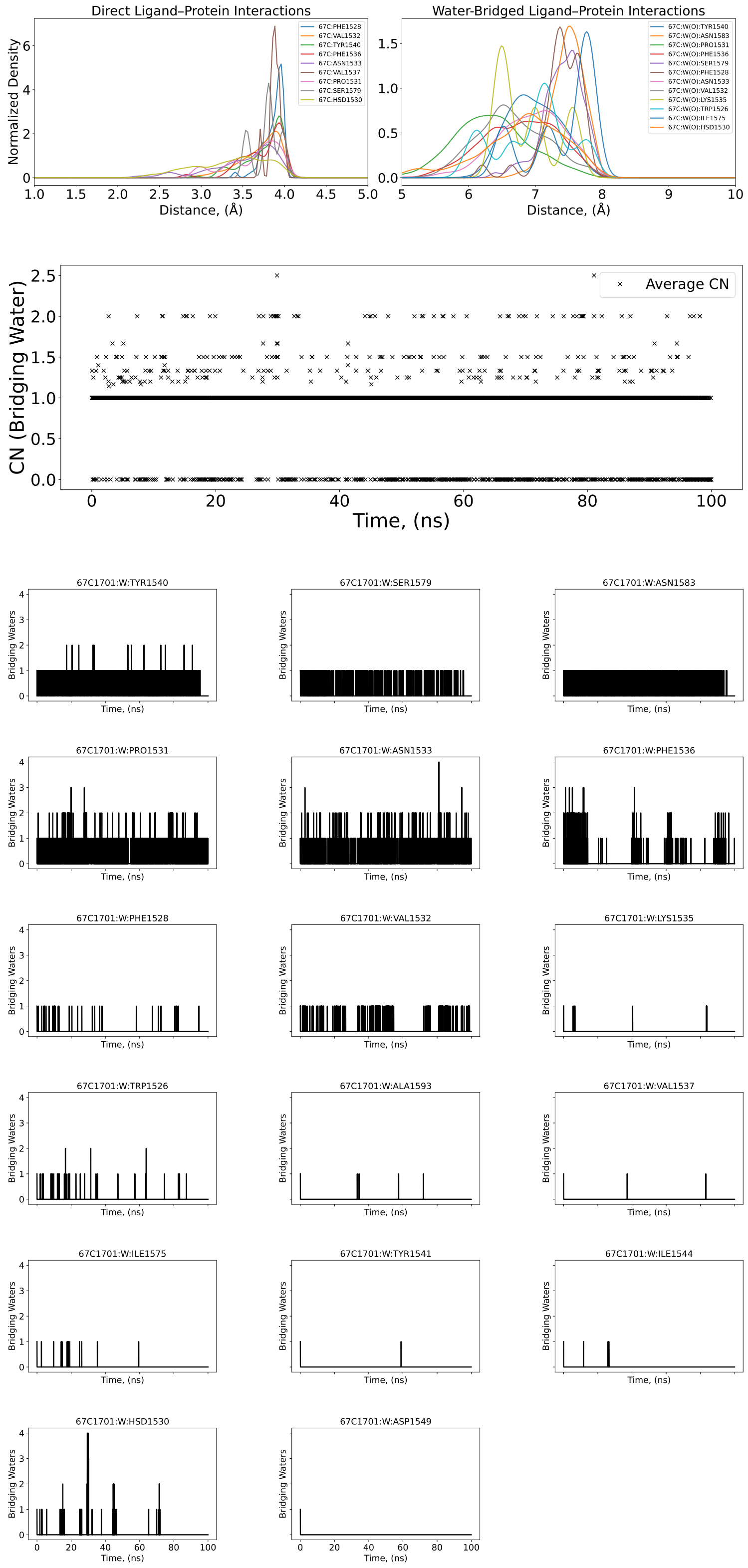


Figure S7. 4-Å–density distribution, averaged and individual coordination number of bridging oxygen water molecules (O/N-bridging) that mediate individual oxygen atoms of the protein and the nitrogen of the ligand along a 100-ns conventional MD for the TAF1(2) complex (PDB: 5I1Q).

minimum distance between each water and the ligand was computed. Water molecules within 10 Å of the ligand were considered site occupants. A binary site-occupancy time series (1 = occupied, 0 = empty) were used for subsequent analysis. Ligand dipole magnitudes were obtained from MD simulation.

To probe coupling between ligand dipole fluctuations and water-site occupancy, we computed the lagged Pearson correlation function. Occupancy and dipole time series were cross-correlated over 4 ns. Positive lag values correspond to dipole changes preceding occupancy changes, while negative lags correspond to occupancy changes preceding dipole changes. Correlations were computed using the standard Pearson coefficient, excluding cases where variance was zero.

The cross-correlation function between ligand dipole magnitude and water-site occupancy is shown in Figure S8. Correlation values remain small across all lag times, with coefficients fluctuating around zero ($|r| < 0.05$). No significant peak was observed at zero lag or at shifted lag times. These results indicate that ligand dipole fluctuations and water-site occupancy are largely uncorrelated on the nanosecond timescale. Time-dependent fluctuations in local water-site occupancy do not exhibit a measurable correlation with the ligand dipole over the analyzed lag interval, which instead is likely dominated by intramolecular electronic fluctuations.

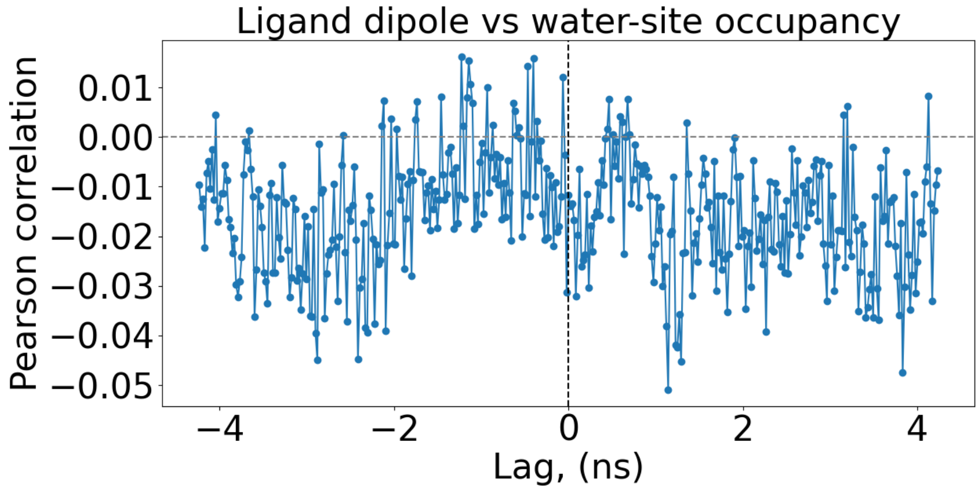


Figure S8. Cross-correlation between ligand dipole magnitude and water-site occupancy. The Pearson correlation coefficient is plotted as a function of lag time (±4 ns). Correlation values remain close to zero, indicating no significant coupling between dipole fluctuations and water occupancy dynamics at the hydration site for the TAF1(2) complex (PDB: 5I1Q).

#### Binding Affinity PMF and Convergence Details

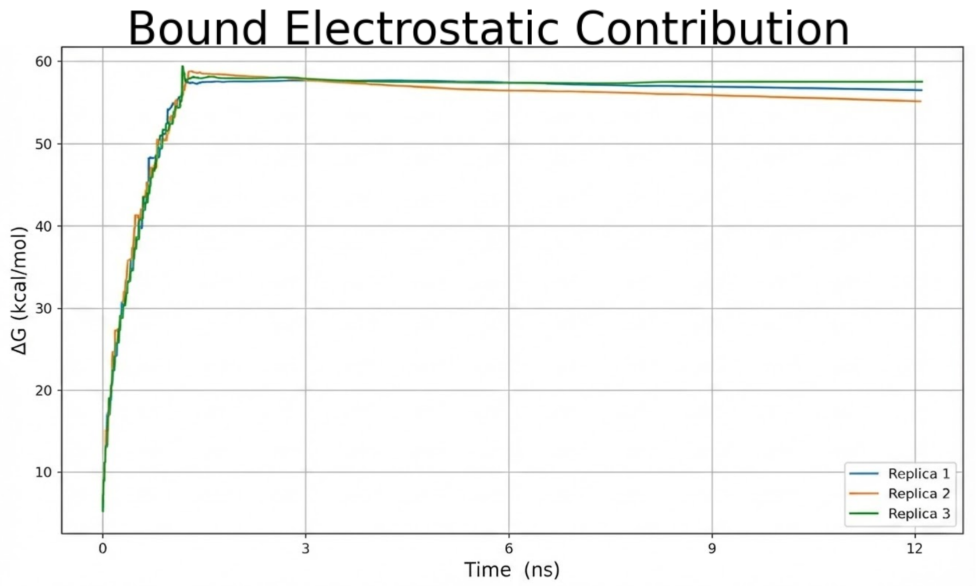


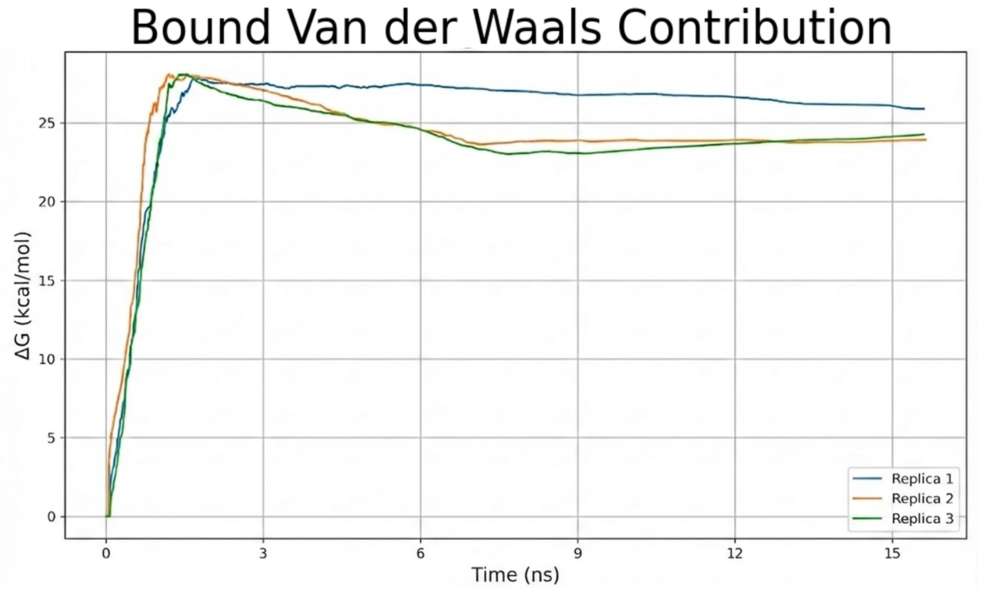


Figure S9. Convergence of the Bound energetic contributions for the TAF1(2) complex (PDB: 5I1Q). The plots display the Free Energy (Δ G) evolution as a function of simulation frames for the Electrostatic (top) and van der Waals (bottom) alchemical legs across three independent replicas. The tight overlap of the trajectories and their stabilization at the end of the production run confirm that the binding affinity calculation has reached thermodynamic equilibrium and is statistically robust.

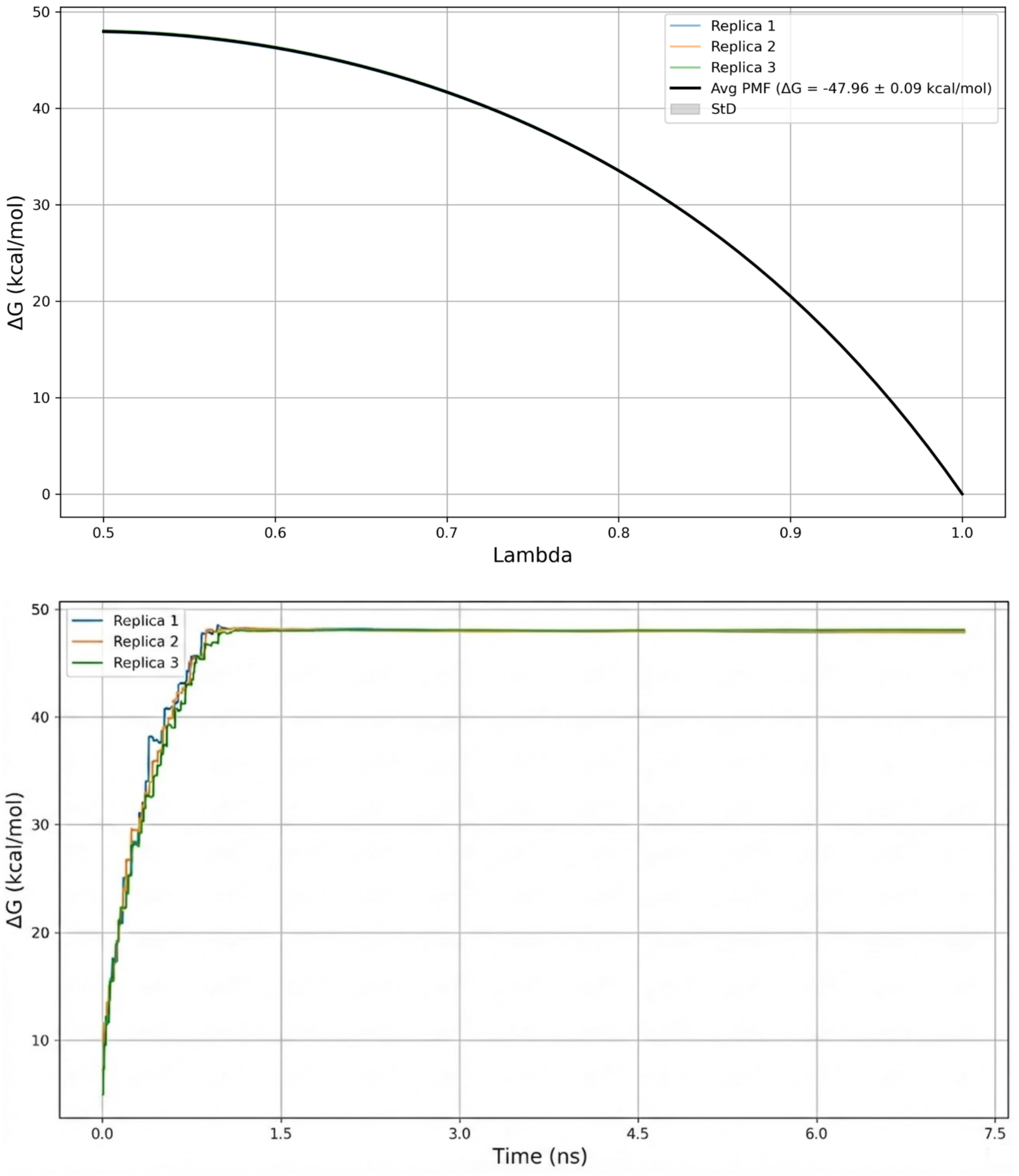


Figure S10. Averaged PMF and convergence plots of the Unbound (Solvent) Electrostatic contribution for the TAF1(2) complex (PDB: 5I1Q). The overlap of the PMF profiles across replicas confirms the precise sampling of the ligand decoupling from the bulk solvent environment.

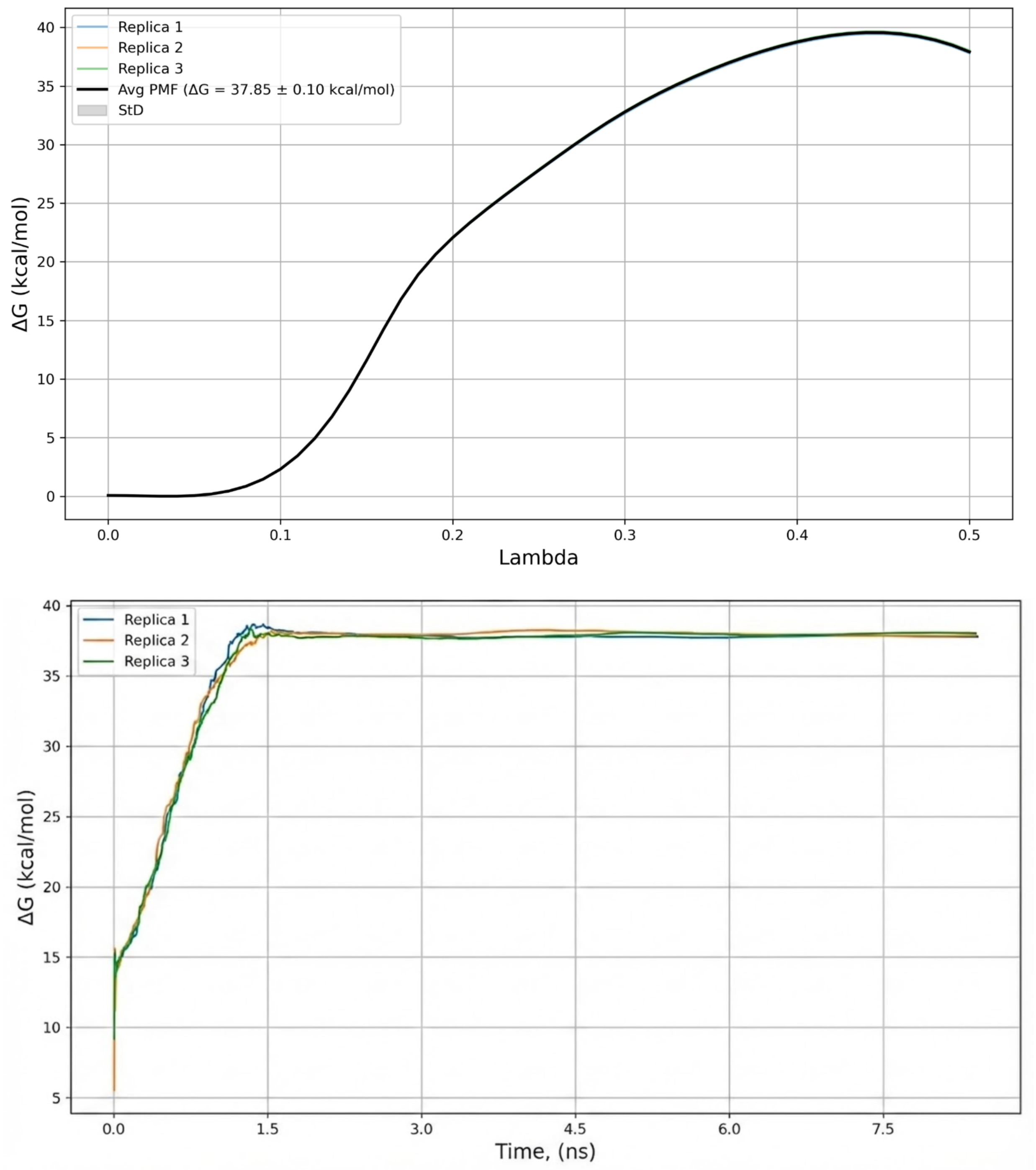


Figure S11. Averaged PMF and convergence plots of the Unbound (Solvent) van der Waals contribution for the TAF1(2) complex (PDB: 5I1Q). The convergence of the decoupling profiles demonstrates reliable sampling of cavity formation energy in the bulk solvent.

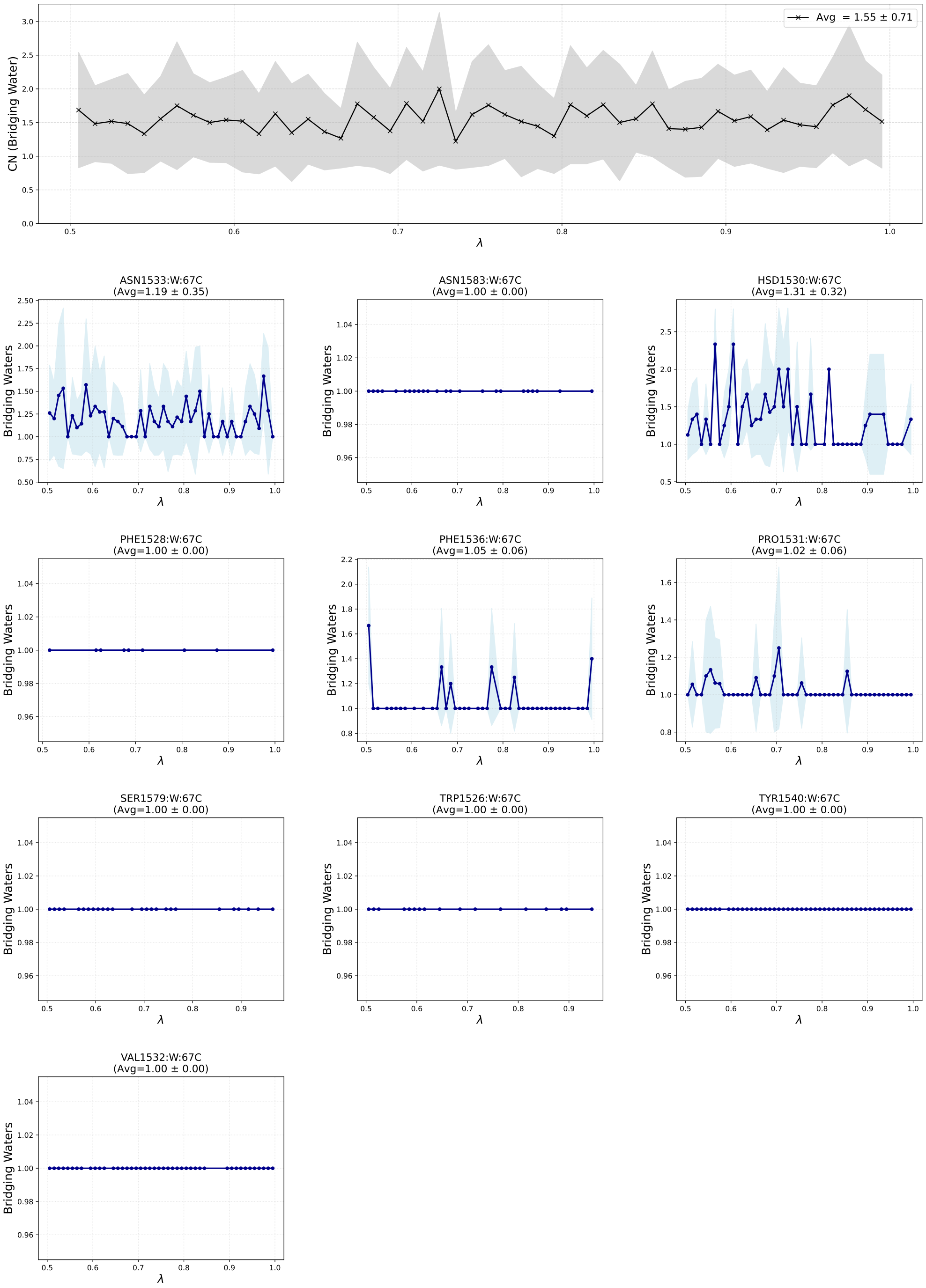


Figure S12. Water-mediated interaction profiles for the O-bridging motif for electrostatic bound contribution in Lambda-ABF-OPES binding affinity calculations for PDB id 5I1Q. (Top Panel) Global analysis of the total hydration stoichiometry within the binding pocket. The black trace represents the conditional average count of water molecules participating in bridging interactions across all detected residue pairs. The gray shaded region indicates the standard deviation ($\pm 1\sigma$) across the ensemble, reflecting fluctuations in solvent multiplicity. (Bottom Panels) Site-specific hydration profiles for key anchoring residues. Each panel displays the average number of water molecules bridging the specific residue–ligand pair as a function of the alchemical coordinate $\lambda$. Error bands represent the fluctuation in water count during bridging events.

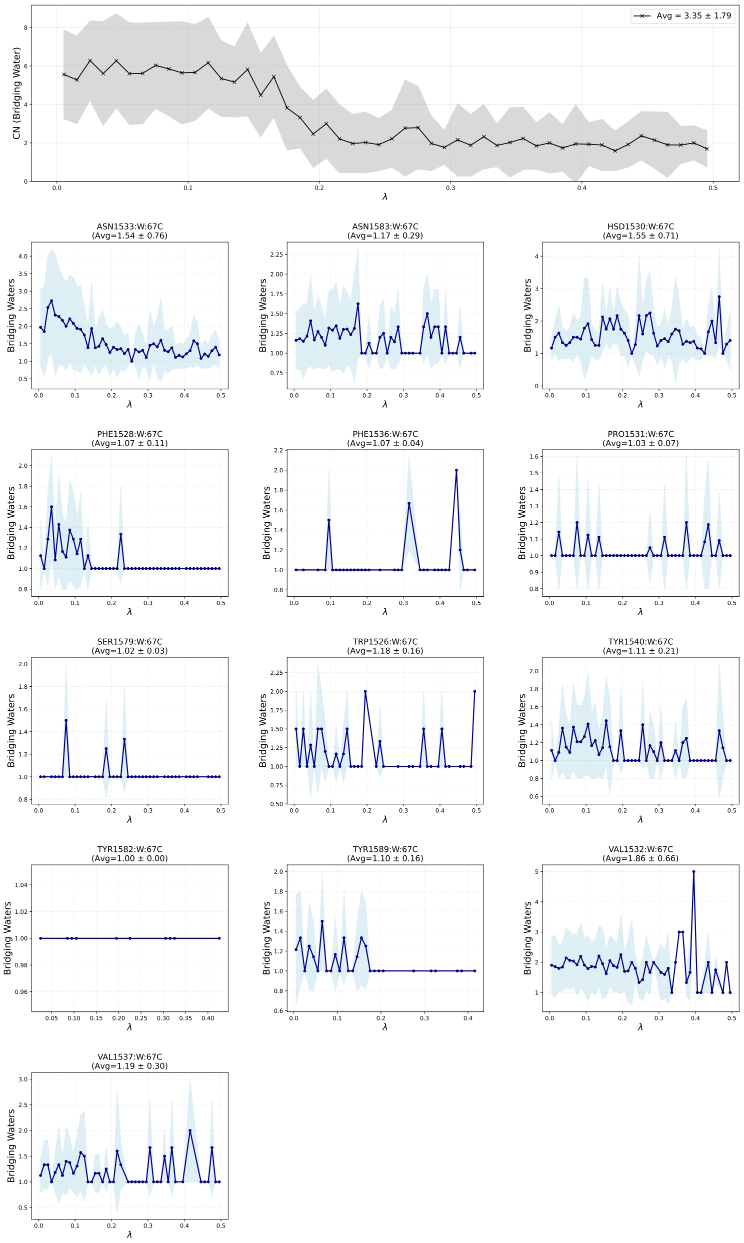


Figure S13. Water-mediated interaction profiles for the O-bridging motif for van der Waals bound contribution in Lambda-ABF-OPES binding affinity calculations for PDB id 5I1Q. (Top Panel) Global analysis of the total hydration stoichiometry within the binding pocket. The black trace represents the conditional average count of water molecules participating in bridging interactions across all detected residue pairs. The gray shaded region indicates the standard deviation ($\pm 1\sigma$) across the ensemble, reflecting fluctuations in solvent multiplicity. (Bottom Panels) Site-specific hydration profiles for key anchoring residues. Each panel displays the average number of water molecules bridging the specific residue–ligand pair as a function of the alchemical coordinate $\lambda$. Error bands represent the fluctuation in water count during bridging events.

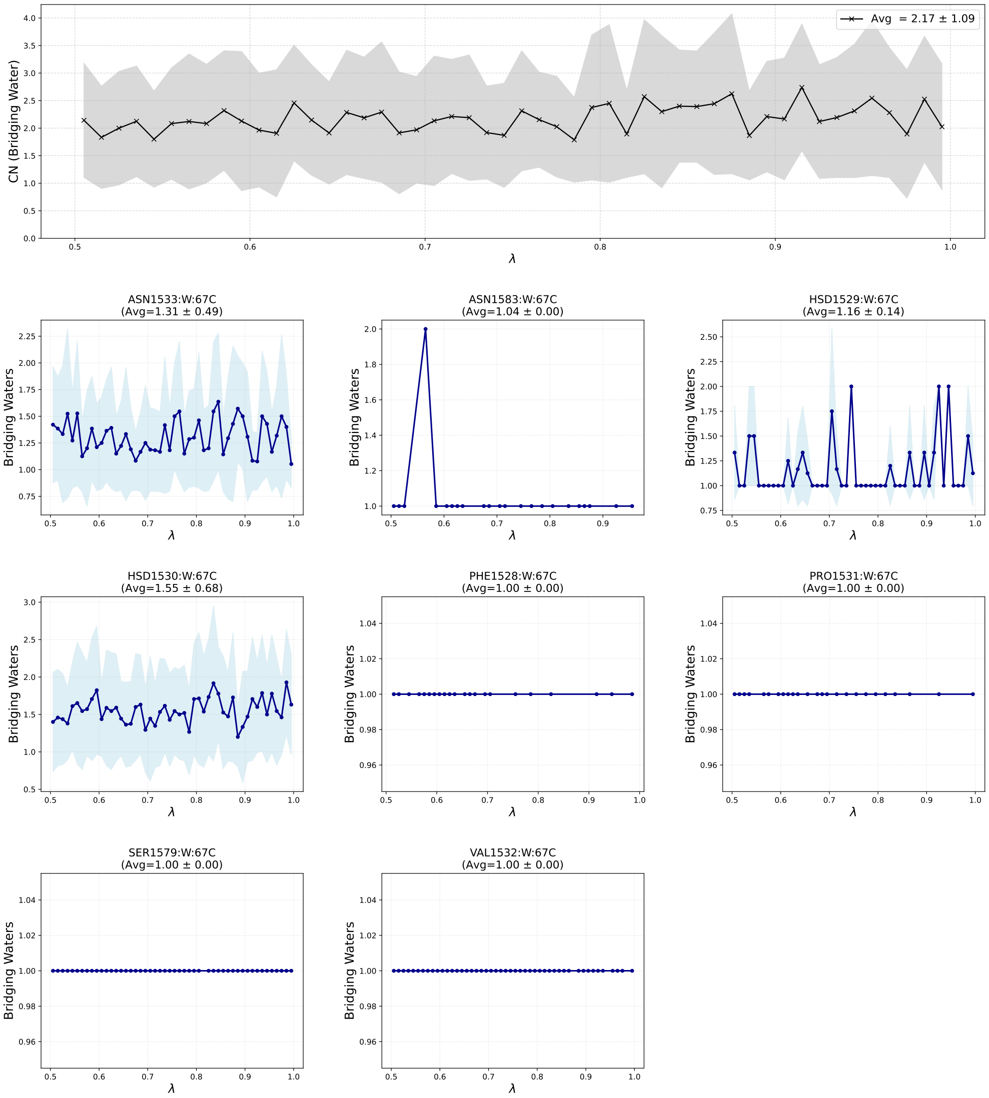


Figure S14. Water-mediated interaction profiles for the N/O-bridging motif for electrostatics bound contribution in Lambda-ABF-OPES binding affinity calculations for PDB id 5I1Q. (Top Panel) Global analysis of the total hydration stoichiometry within the binding pocket. The black trace represents the conditional average count of water molecules participating in bridging interactions across all detected residue pairs. The gray shaded region indicates the standard deviation ($\pm 1\sigma$) across the ensemble, reflecting fluctuations in solvent multiplicity. (Bottom Panels) Site-specific hydration profiles for key anchoring residues. Each panel displays the average number of water molecules bridging the specific residue–ligand pair as a function of the alchemical coordinate $\lambda$. Error bands represent the fluctuation in water count during bridging events.

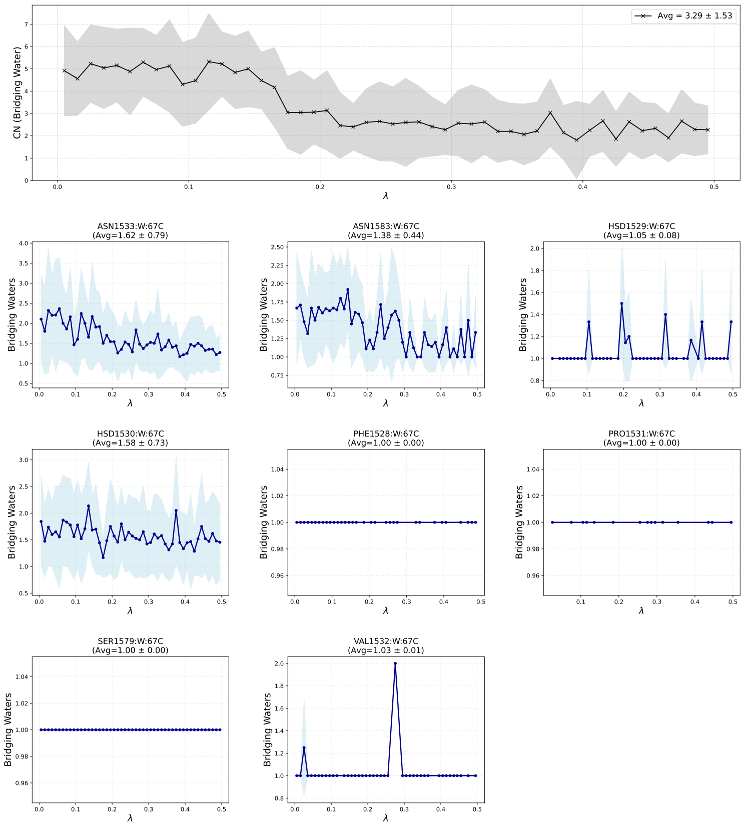


Figure S15. Water-mediated interaction profiles for the N/O-bridging motif for van der Waals bound contribution in Lambda-ABF-OPES binding affinity calculations for PDB id 5I1Q. (Top Panel) Global analysis of the total hydration stoichiometry within the binding pocket. The black trace represents the conditional average count of water molecules participating in bridging interactions across all detected residue pairs. The gray shaded region indicates the standard deviation ($\pm 1\sigma$) across the ensemble, reflecting fluctuations in solvent multiplicity. (Bottom Panels) Site-specific hydration profiles for key anchoring residues. Each panel displays the average number of water molecules bridging the specific residue–ligand pair as a function of the alchemical coordinate $\lambda$. Error bands represent the fluctuation in water count during bridging events.

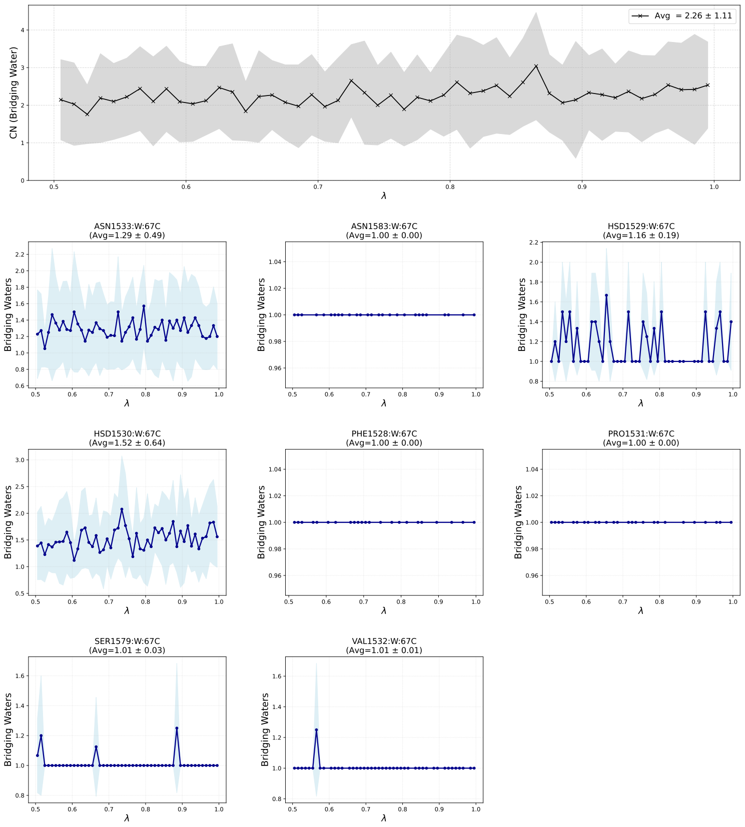


Figure S16. Water-mediated interaction profiles for the N-bridging motif for electrostatics bound contribution in Lambda-ABF-OPES binding affinity calculations for PDB id 5I1Q. (Top Panel) Global analysis of the total hydration stoichiometry within the binding pocket. The black trace represents the conditional average count of water molecules participating in bridging interactions across all detected residue pairs. The gray shaded region indicates the standard deviation ($\pm 1\sigma$) across the ensemble, reflecting fluctuations in solvent multiplicity. (Bottom Panels) Site-specific hydration profiles for key anchoring residues. Each panel displays the average number of water molecules bridging the specific residue–ligand pair as a function of the alchemical coordinate $\lambda$. Error bands represent the fluctuation in water count during bridging events.

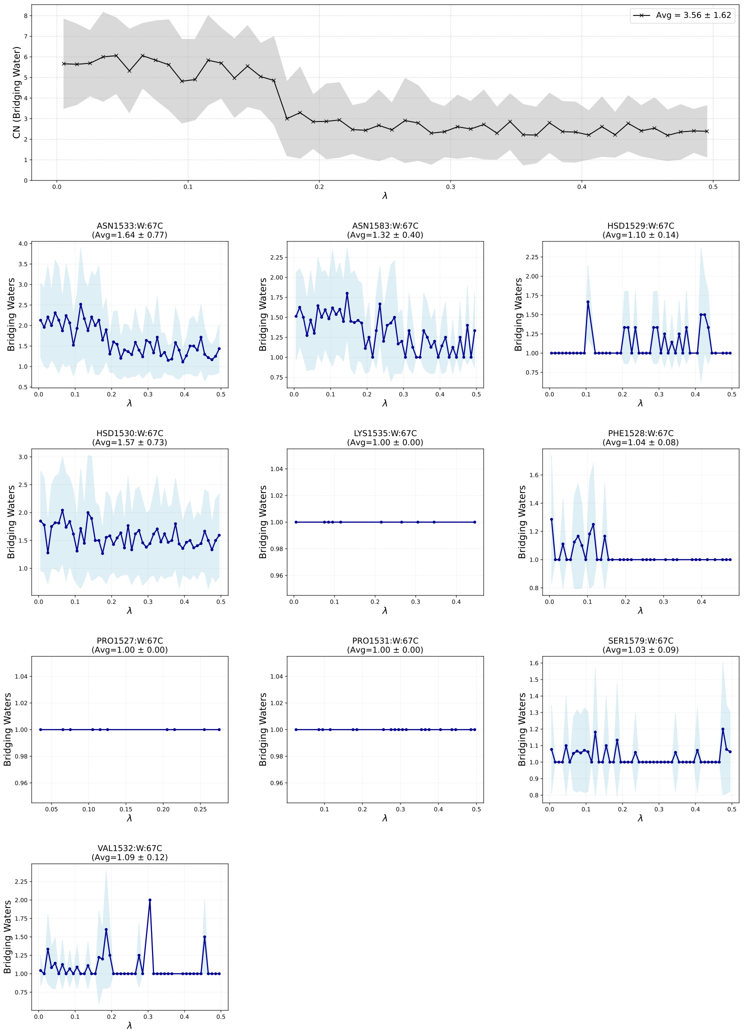


Figure S17. Water-mediated interaction profiles for the N-bridging motif for van der Waals bound contribution in Lambda-ABF-OPES binding affinity calculations for PDB id 5I1Q. (Top Panel) Global analysis of the total hydration stoichiometry within the binding pocket. The black trace represents the conditional average count of water molecules participating in bridging interactions across all detected residue pairs. The gray shaded region indicates the standard deviation ($\pm 1\sigma$) across the ensemble, reflecting fluctuations in solvent multiplicity. (Bottom Panels) Site-specific hydration profiles for key anchoring residues. Each panel displays the average number of water molecules bridging the specific residue–ligand pair as a function of the alchemical coordinate $\lambda$. Error bands represent the fluctuation in water count during bridging events.

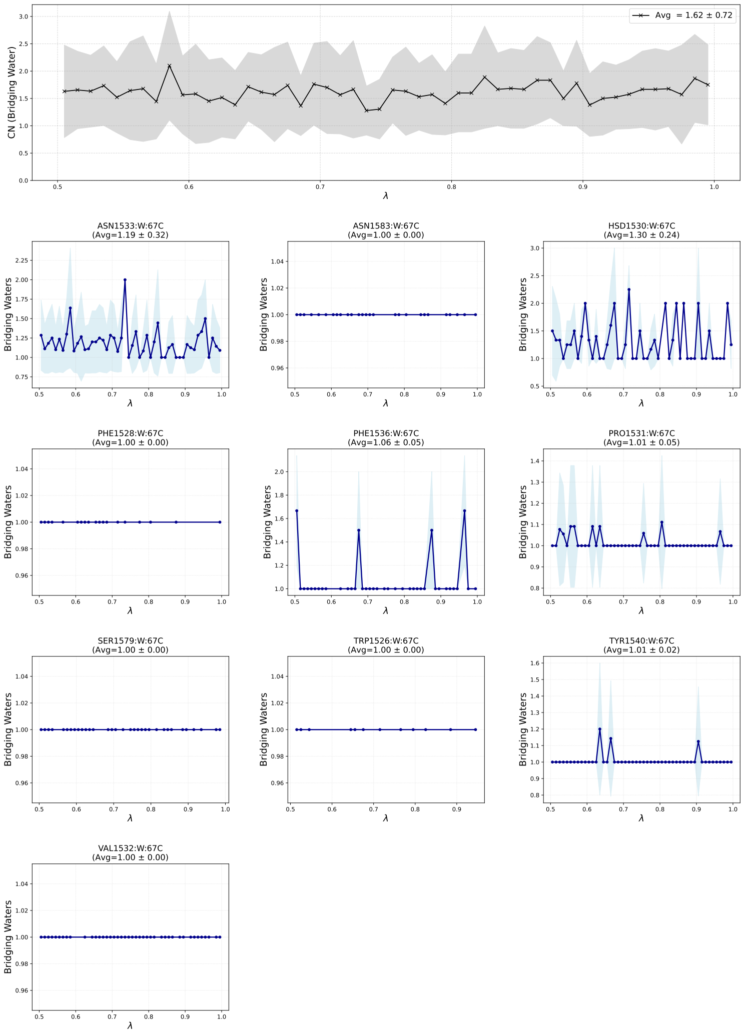


Figure S18. Water-mediated interaction profiles for the O/N-bridging motif for electrostatics bound contribution in Lambda-ABF-OPES binding affinity calculations for PDB id 5I1Q. (Top Panel) Global analysis of the total hydration stoichiometry within the binding pocket. The black trace represents the conditional average count of water molecules participating in bridging interactions across all detected residue pairs. The gray shaded region indicates the standard deviation ($\pm 1\sigma$) across the ensemble, reflecting fluctuations in solvent multiplicity. (Bottom Panels) Site-specific hydration profiles for key anchoring residues. Each panel displays the average number of water molecules bridging the specific residue–ligand pair as a function of the alchemical coordinate $\lambda$. Error bands represent the fluctuation in water count during bridging events.

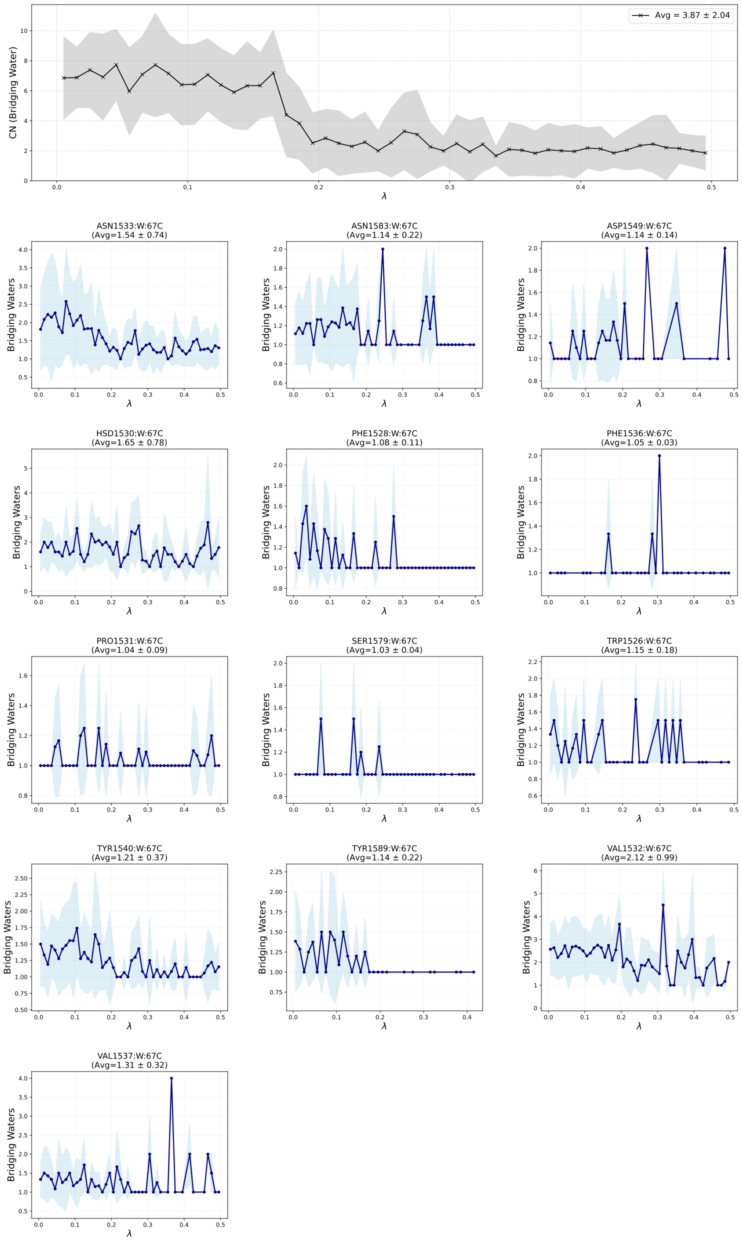


Figure S19. Water-mediated interaction profiles for the O/N-bridging motif for van der Waals bound contribution in Lambda-ABF-OPES binding affinity calculations for PDB id 5I1Q. (Top Panel) Global analysis of the total hydration stoichiometry within the binding pocket. The black trace represents the conditional average count of water molecules participating in bridging interactions across all detected residue pairs. The gray shaded region indicates the standard deviation ($\pm 1\sigma$) across the ensemble, reflecting fluctuations in solvent multiplicity. (Bottom Panels) Site-specific hydration profiles for key anchoring residues. Each panel displays the average number of water molecules bridging the specific residue–ligand pair as a function of the alchemical coordinate $\lambda$. Error bands represent the fluctuation in water count during bridging events.

#### Matched hydration-suppression control for TYR1540 in 5I1Q

Coordination numbers, site occupancies, and water-density maps establish where and how frequently water configurations are sampled, but they do not by themselves establish whether the corresponding hydration response is thermodynamically coupled to the alchemical transformation. We therefore performed a matched perturbational control in which the principal $\lambda$-dependent TYR1540 hydration transition was deliberately suppressed while retaining the remaining bound-state Lambda-ABF-OPES protocol.

##### Selection of the controlled hydration feature

The 5I1Q complex was selected because its bound-state van der Waals leg shows a pronounced increase in water coordination around the TYR1540 hydroxyl oxygen when $\lambda < 0.2$. This feature provides a clearly resolved hydration transition that can be selectively perturbed. The control retained exactly the same TYR1540 coordination-number definition used in the main-text structural analysis.

For an instantaneous molecular configuration $\mathbf{q}$, the TYR1540 hydration coordinate was defined as

$$c_{\mathrm{TYR1540}}(\mathbf{q}) = \sum_{w \in W} s(r_w), \qquad s(r_w) = \frac{1}{1 + (r_w/r_0)^6}, \tag{1}$$

where $W$ is the set of all water oxygen atoms, $r_w$ is the instantaneous distance between water oxygen $w$ and the TYR1540 hydroxyl oxygen, and $r_0 = 4.0$ Å is the switching distance. A water close to the TYR1540 hydroxyl oxygen contributes approximately one, its contribution is $0.5$ at $r_w = r_0$, and its contribution decreases smoothly toward zero at larger distances. Consequently, $c_{\mathrm{TYR1540}}$ is a continuous, dimensionless hydration measure rather than an integer number of water molecules.

#### Conditional hydration-suppression force

In the restrained calculations, an additional generalized force was applied to $c_{\mathrm{TYR1540}}$ only in the low-$\lambda$ region where the unrestrained hydration increase occurs:

$$F_c(\lambda, c_{\mathrm{TYR1540}}) = \begin{cases} -k\, c_{\mathrm{TYR1540}}, & \lambda < 0.2, \\ 0, & \lambda \geq 0.2. \end{cases} \tag{2}$$

At a fixed value of $\lambda < 0.2$, this generalized force is equivalent to the derivative of a zero-centered harmonic potential along the coordination coordinate,

$$U_k(c_{\mathrm{TYR1540}}) = \frac{1}{2} k\, c_{\mathrm{TYR1540}}^2, \qquad F_c = -\frac{\partial U_k}{\partial c_{\mathrm{TYR1540}}} = -k\, c_{\mathrm{TYR1540}}. \tag{3}$$

Two force constants were examined: $k = 10\,\mathrm{kcal\,mol^{-1}}$, denoted K10, and $k = 20\,\mathrm{kcal\,mol^{-1}}$, denoted K20.. At the same instantaneous coordination number, K20 applies twice the generalized restoring force and twice the harmonic energy penalty of K10. For example, at $c_{\mathrm{TYR1540}} = 3$, the applied generalized forces are $-30$ and $-60$ for K10 and K20, respectively.

The force was introduced as a diagnostic suppression perturbation. It was not used as a restraint correction and was not added to the production binding-affinity estimate.

#### Matched simulation protocol

Three matched conditions were examined:

1. an unrestrained monitor, in which $c_{\mathrm{TYR1540}}$ was recorded without an additional force;
2. the K10 hydration-suppression control;
3. the K20 hydration-suppression control.

Four walkers were propagated for each condition. All starting coordinates, AMOEBA parameters, Lambda-ABF-OPES settings, alchemical boundaries, DBC restraints, and ligand–ion treatment were otherwise retained.

Each walker was propagated for $10\ \mathrm{ns}$ per walker. The three conditions were therefore compared at exactly matched sampling duration.

##### Analysis criteria

The control was evaluated in three stages. First, the TYR1540 coordination number was analyzed as a function of the instantaneous alchemical coordinate to verify that the low-$\lambda$ high-coordination branch was present in the unrestrained calculation and suppressed in the K10 and K20 calculations. Second, the implementation was checked by verifying that

$$\frac{F_c}{c_{\mathrm{TYR1540}}} = -10 \quad \text{or} \quad -20 \qquad \text{for } \lambda < 0.2, \tag{4}$$

for K10 and K20, respectively, and that $F_c = 0$ for $\lambda \geq 0.2$. Third, DBC was analyzed as a function of $\lambda$ to determine whether suppression of hydration was accompanied by gross displacement of the ligand from the restrained bound-state region.

##### Suppression of the TYR1540 hydration transition

As shown in Fig. S20a, the unrestrained calculation reproduces the pronounced high-coordination branch around TYR1540 at $\lambda < 0.2$. This branch is removed by K10, demonstrating that $k = 10$ is already sufficient to suppress the targeted hydration transition. K20 produces a still lower-coordination ensemble and therefore constitutes a stronger perturbation of the local solvent response.

The applied-force analysis confirmed the expected force ratios in the active interval and an exactly zero additional coordination force for $\lambda \geq 0.2$. All three conditions continued to visit the complete $\lambda \in [0, 0.5]$ interval.

The DBC distributions remain strongly overlapping among the unrestrained, K10, and K20 conditions (Fig. S20b). Thus, the disappearance of the high-coordination TYR1540 branch is not accompanied by an evident gross displacement of the ligand from the restrained binding-site region.

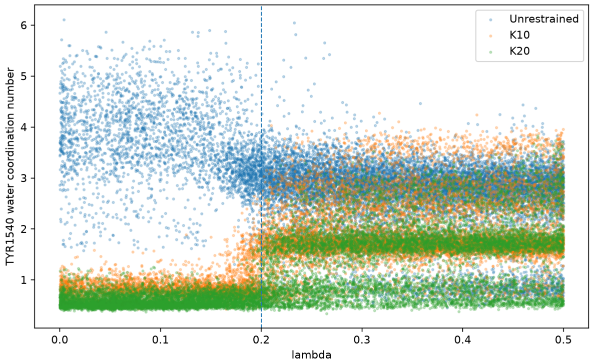


(a) TYR1540 water coordination number versus $\lambda$.

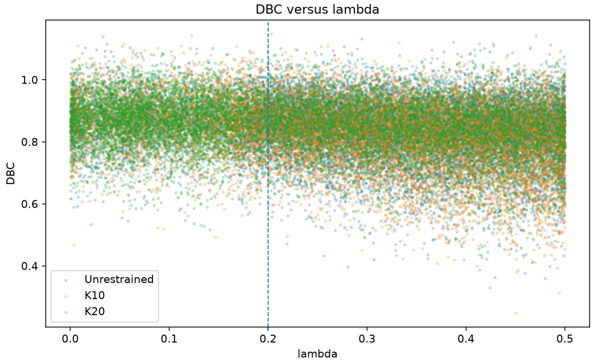


(b) DBC distribution versus $\lambda$.

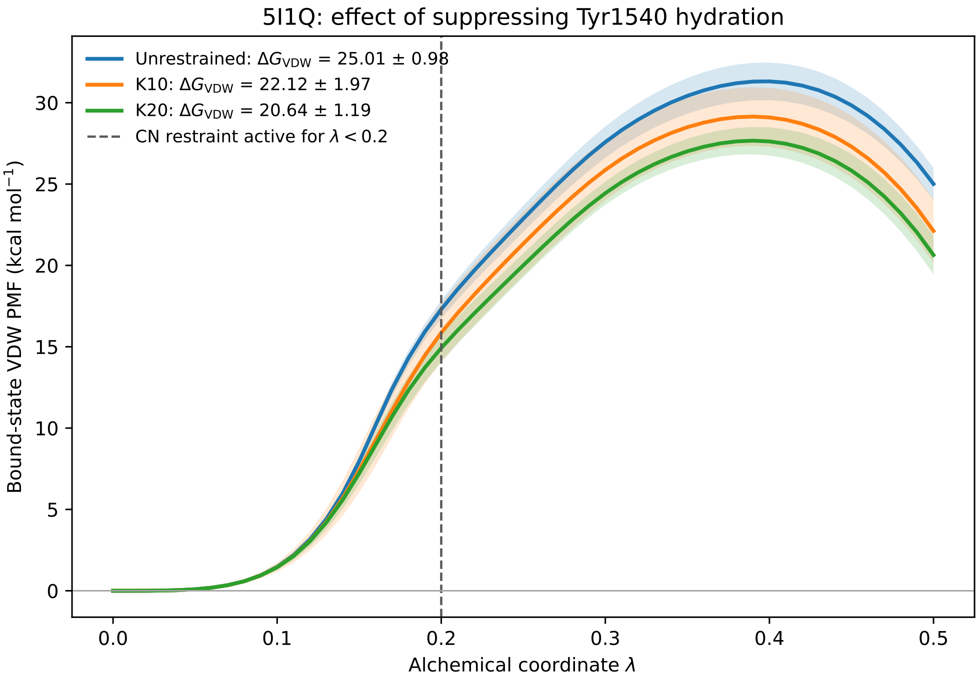


(c) Matched-duration bound-state van der Waals PMFs.

Figure S20. Matched hydration-suppression control for the TYR1540 hydration transition in the 5I1Q complex. The vertical dashed line marks $\lambda = 0.2$, below which the conditional coordination-number force was applied. **(a)** The unrestrained calculation samples a pronounced high-coordination branch below $\lambda = 0.2$, which is removed by K10 and K20. **(b)** The DBC distributions remain strongly overlapping, indicating no evident gross displacement of the ligand. **(c)** Bound-state van der Waals PMFs obtained from four matched-duration walkers per condition. Solid curves show the walker mean and shaded regions show the between-walker standard deviation.

#### Effect on the bound-state van der Waals PMF

The matched-duration PMF analysis yielded

$$\Delta G_{\mathrm{VDW}}^{\mathrm{unrestrained}} = 25.01 \pm 0.98\ \mathrm{kcal\,mol^{-1}}, \tag{5}$$

$$\Delta G_{\mathrm{VDW}}^{K10} = 22.12 \pm 1.97\ \mathrm{kcal\,mol^{-1}}, \tag{6}$$

$$\Delta G_{\mathrm{VDW}}^{K20} = 20.64 \pm 1.19\ \mathrm{kcal\,mol^{-1}}. \tag{7}$$

The mean shifts relative to the unrestrained calculation are therefore

$$\Delta\Delta G_{K10} = \Delta G_{\mathrm{VDW}}^{K10} - \Delta G_{\mathrm{VDW}}^{\mathrm{unrestrained}} = -2.89\ \mathrm{kcal\,mol^{-1}}, \tag{8}$$

$$\Delta\Delta G_{K20} = \Delta G_{\mathrm{VDW}}^{K20} - \Delta G_{\mathrm{VDW}}^{\mathrm{unrestrained}} = -4.37\ \mathrm{kcal\,mol^{-1}}. \tag{9}$$

Thus, suppression of the TYR1540 hydration transition decreases the calculated bound-state van der Waals contribution by approximately $2.9\ \mathrm{kcal\,mol^{-1}}$ with K10 and $4.4\ \mathrm{kcal\,mol^{-1}}$ with K20. These decreases correspond to approximately $12\%$ and $17\%$, respectively, relative to the unrestrained value.

The direction of this response is consistent with solvent entry stabilizing weakly coupled configurations near $\lambda = 0$. In the unrestrained calculation, removal of ligand van der Waals interactions permits increased TYR1540 hydration, which lowers the free energy of the low-$\lambda$ ensemble relative to the fully coupled state. Suppressing this solvent stabilization decreases the endpoint separation, producing the lower $\Delta G_{\mathrm{VDW}}$ values obtained for K10 and K20.

K20 decreases $\Delta G_{\mathrm{VDW}}$ by an additional $1.48\ \mathrm{kcal\,mol^{-1}}$ relative to K10. However, K10 already eliminates the targeted high-coordination branch. The additional change under K20 therefore reflects a stronger perturbation of the hydration ensemble rather than evidence that K10 was insufficient.

##### Interpretation and limitations

The combined analysis establishes the intended causal sequence: the unrestrained calculation samples the TYR1540 hydration transition, while K10 and K20 suppress this transition. The DBC behavior remains comparable; and the bound-state van der Waals PMF changes systematically in response to the hydration perturbation. These observations support thermodynamic coupling between local TYR1540 rehydration and ligand van der Waals decoupling.

The control does not establish that the unrestricted simulation reconstructs an independently validated absolute hydration free-energy landscape. The additional force defines an artificial, $\lambda$-conditioned perturbation of the bound-state Hamiltonian. Consequently, the PMF differences are not absolute water-insertion free energies, individual-water contributions, or corrections to the production binding affinity. They quantify the sensitivity of the alchemical PMF to suppression of the local TYR1540 hydration response.

# 1I05

The labels O/O, N/O, O/N, and N/N denote the heavy-atom element classes used for contact classification, with the first element corresponding to the protein atom and the second to the ligand atom. Residue labels in the figures are used as compact notation. The corresponding atom-level assignments, including protein atom name and backbone versus side-chain origin, are reported in Table S4.

**Table S4. Atom-level direct and water-mediated polar contacts for 1i05. Motif labels are reported as protein-element/ligand-element. Direct contacts use a $4.0$ Å heavy-atom cutoff. Water-mediated contacts require both ligand–water and water–protein distances to be within $4.0$ Å and ligand–protein proximity $\leq 6.0$ Å.**

| contact type | motif | contact label | atom origin | occ. fraction | mean events | direct dist Å | lig–wat dist Å | wat–prot dist Å | total path Å | unique water O |
|---|---|---|---|---|---|---|---|---|---|---|
| direct | O/O | LTL408:O2–PHE74:O (backbone) | backbone | 0.903 | 1.00 | 3.27 | | | | |
| direct | O/O | LTL408:O2–TYR138:O (backbone) | backbone | 0.512 | 1.00 | 3.77 | | | | |
| direct | O/O | LTL408:O1–PHE74:O (backbone) | backbone | 0.344 | 1.00 | 3.44 | | | | |
| direct | O/O | LTL408:O2–TYR138:OH (side chain) | side chain | 0.283 | 1.00 | 3.83 | | | | |
| direct | O/O | LTL408:O2–LEU134:O (backbone) | backbone | 0.102 | 1.00 | 3.77 | | | | |
| direct | O/O | LTL408:O1–LEU134:O (backbone) | backbone | 0.050 | 1.00 | 3.64 | | | | |
| direct | O/O | LTL408:O2–LEU123:O (backbone) | backbone | 0.048 | 1.00 | 3.44 | | | | |
| direct | O/O | LTL408:O1–TYR138:O (backbone) | backbone | 0.041 | 1.00 | 3.15 | | | | |
| direct | O/O | LTL408:O1–TYR138:OH (side chain) | side chain | 0.039 | 1.00 | 3.30 | | | | |
| direct | O/O | LTL408:O1–LEU123:O (backbone) | backbone | 0.031 | 1.00 | 3.41 | | | | |
| direct | O/O | LTL408:O1–LEU72:O (backbone) | backbone | 0.027 | 1.00 | 3.50 | | | | |
| direct | O/O | LTL408:O2–LEU58:O (backbone) | backbone | 0.012 | 1.00 | 3.74 | | | | |
| water-mediated | O/O | LTL408:O2–W(O)–TYR138:O (backbone) | backbone | 0.835 | 1.49 | | 3.64 | 2.81 | 6.45 | 2 |

*Continued on next page*

#### Table S4. Atom-level direct and water-mediated polar contacts for 1i05 (continued)

| contact type | motif | contact label | atom origin | occ. fraction | mean events | direct dist Å | lig–wat dist Å | wat–prot dist Å | total path Å | unique water O |
|---|---|---|---|---|---|---|---|---|---|---|
| water-mediated | O/O | LTL408:O2–W(O)–LEU134:O (backbone) | backbone | 0.636 | 1.03 | | 3.60 | 3.30 | 6.90 | 2 |
| water-mediated | O/O | LTL408:O2–W(O)–TYR138:OH (side chain) | side chain | 0.618 | 1.02 | | 3.68 | 2.77 | 6.45 | 2 |
| water-mediated | O/O | LTL408:O2–W(O)–LEU42:O (backbone) | backbone | 0.352 | 1.56 | | 3.63 | 3.27 | 6.90 | 2 |
| water-mediated | O/O | LTL408:O2–W(O)–LEU58:O (backbone) | backbone | 0.071 | 1.00 | | 3.60 | 3.17 | 6.78 | 2 |
| water-mediated | O/O | LTL408:O1–W(O)–TYR138:O (backbone) | backbone | 0.045 | 1.58 | | 3.11 | 2.59 | 5.69 | 2 |
| water-mediated | O/O | LTL408:O1–W(O)–TYR138:OH (side chain) | side chain | 0.043 | 1.03 | | 3.17 | 2.76 | 5.93 | 2 |
| water-mediated | O/O | LTL408:O1–W(O)–LEU134:O (backbone) | backbone | 0.034 | 1.06 | | 3.02 | 3.15 | 6.17 | 2 |
| water-mediated | N/O | LTL408:O2–W(O)–LEU58:N (backbone) | backbone | 0.032 | 1.21 | | 3.57 | 3.56 | 7.13 | 2 |
| water-mediated | O/O | LTL408:O1–W(O)–LEU42:O (backbone) | backbone | 0.027 | 1.65 | | 3.06 | 3.29 | 6.34 | 2 |
| water-mediated | O/O | LTL408:O1–W(O)–LEU58:O (backbone) | backbone | 0.018 | 1.00 | | 3.04 | 3.07 | 6.11 | 1 |
| water-mediated | N/O | LTL408:O1–W(O)–LEU58:N (backbone) | backbone | 0.014 | 1.25 | | 2.95 | 3.64 | 6.59 | 2 |
| water-mediated | O/O | LTL408:O2–W(O)–LEU123:O (backbone) | backbone | 0.013 | 1.00 | | 3.57 | 3.77 | 7.34 | 1 |
| water-mediated | O/O | LTL408:O2–W(O)–PHE74:O (backbone) | backbone | 0.011 | 1.00 | | 3.61 | 3.85 | 7.46 | 1 |

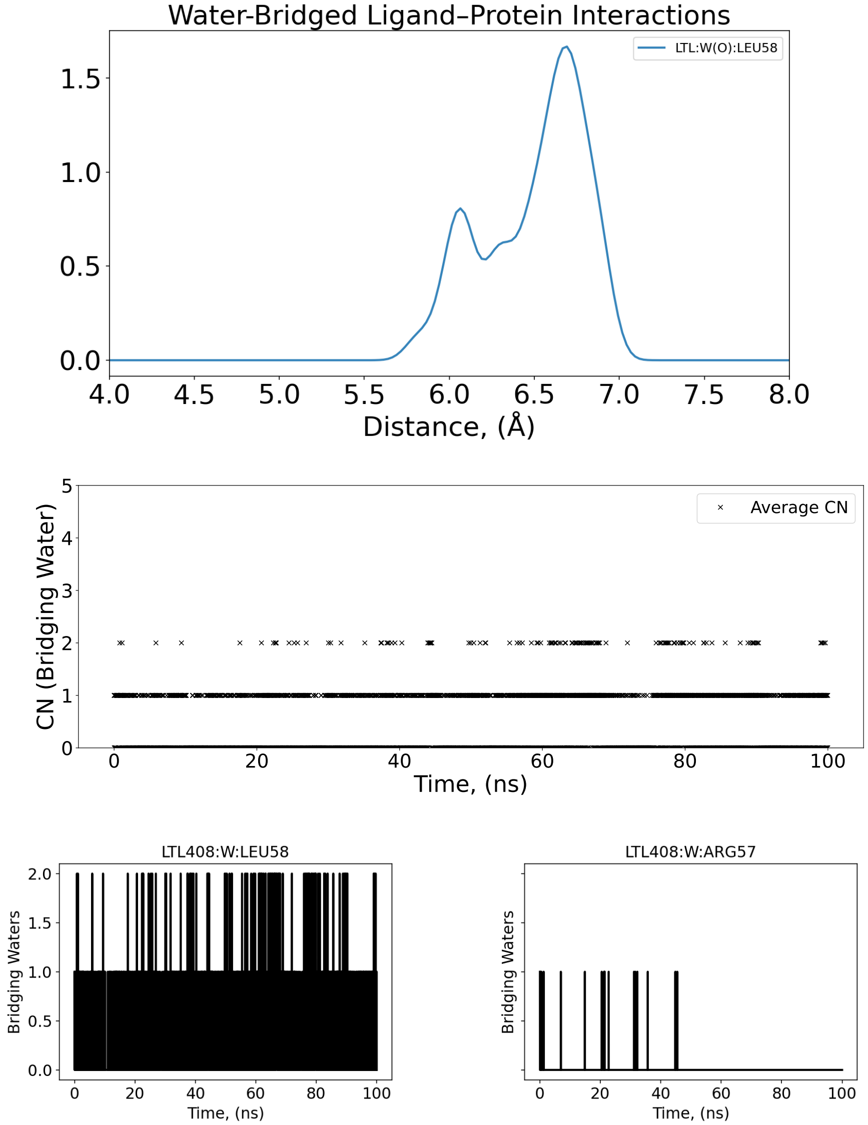


Figure S21. 4-Å–density distribution, averaged and individual coordination number of bridging oxygen water molecules (N/O-bridging) that mediate individual nitrogen atoms of the protein and oxygen atoms of the ligand along a 100-ns conventional MD for MUP-I (PDB: 1I05). The absence of significant density peaks shows that direct N/O bridging is not a primary stabilization mechanism in this deeply buried hydrophobic pocket.

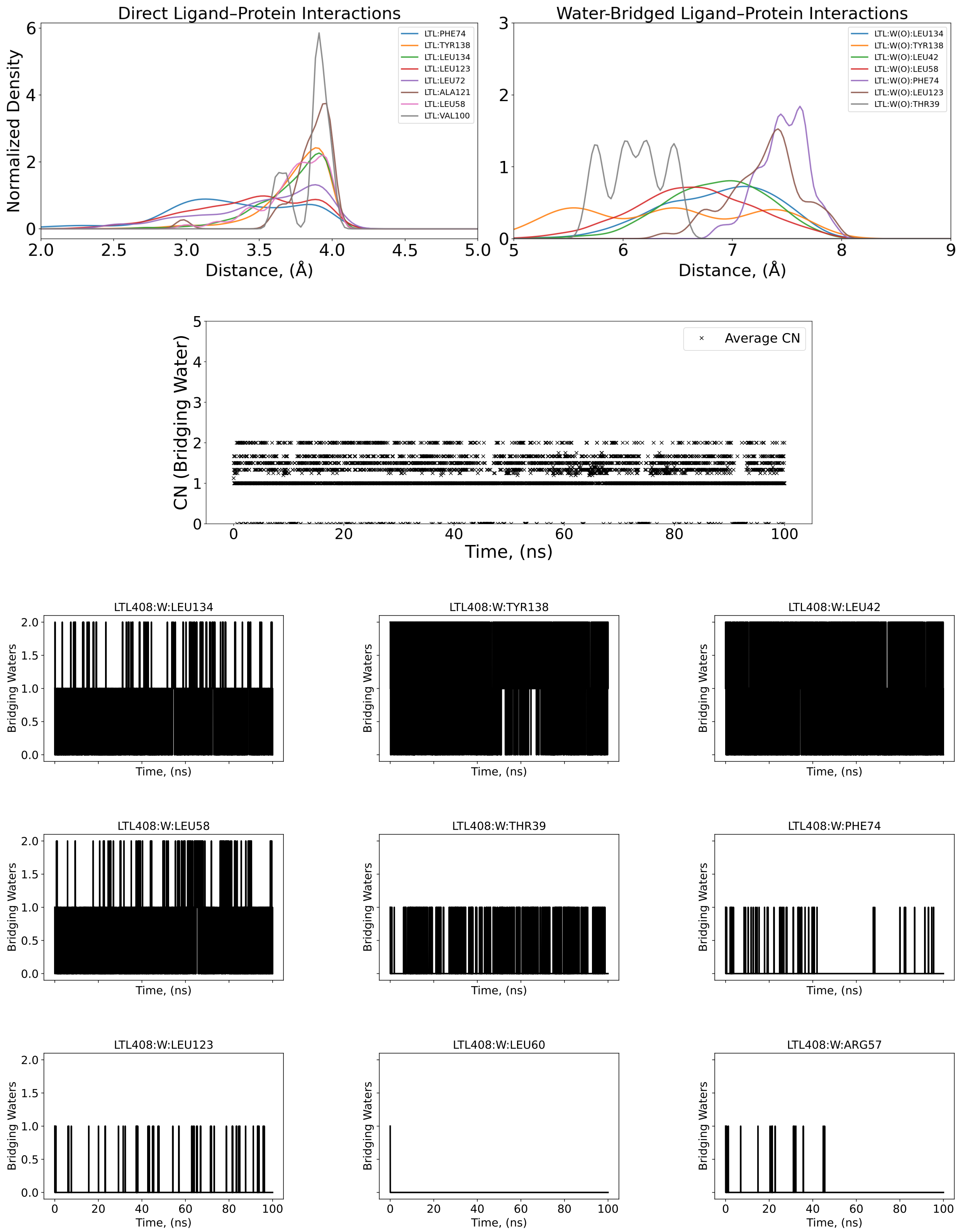


Figure S22. 4-Å–density distribution, averaged and individual coordination number of bridging oxygen water molecules (O-bridging) that mediate individual oxygen atoms of the protein and oxygen atoms of the ligand along a 100-ns conventional MD for MUP-I (PDB: 1I05).

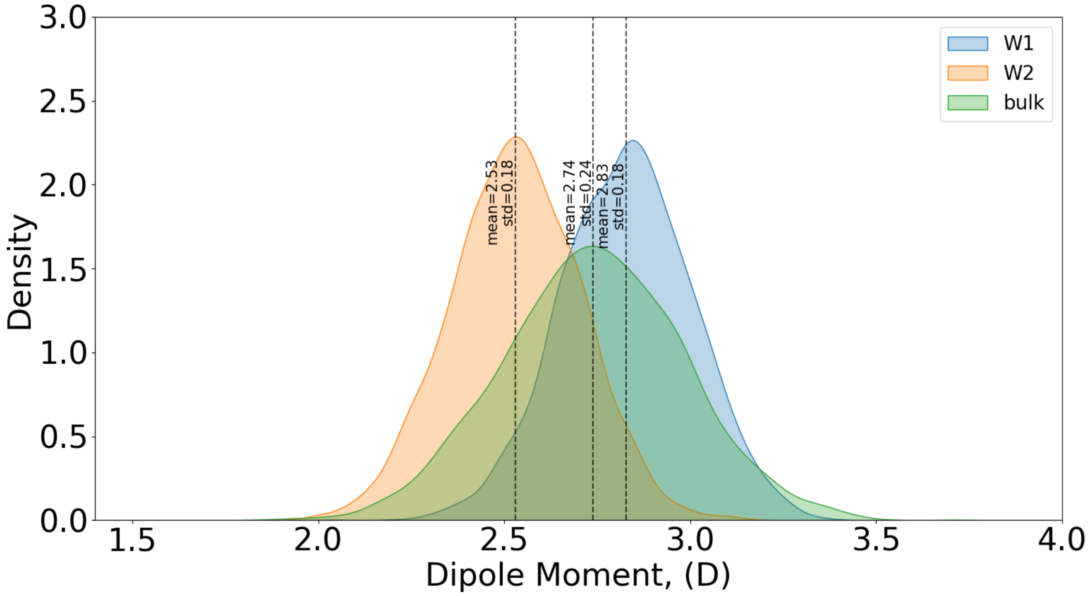


Figure S23. Dipole Moment Distributions for MUP-I complex (PDB: 1I05). Distribution of instantaneous dipole moments for water molecules residing in sites W1 (orange) and W2 (blue) compared to bulk water (green). The W2 distribution indicates that the trapped water molecules retain bulk-like electrostatic properties and is not strongly polarized by the protein environment.

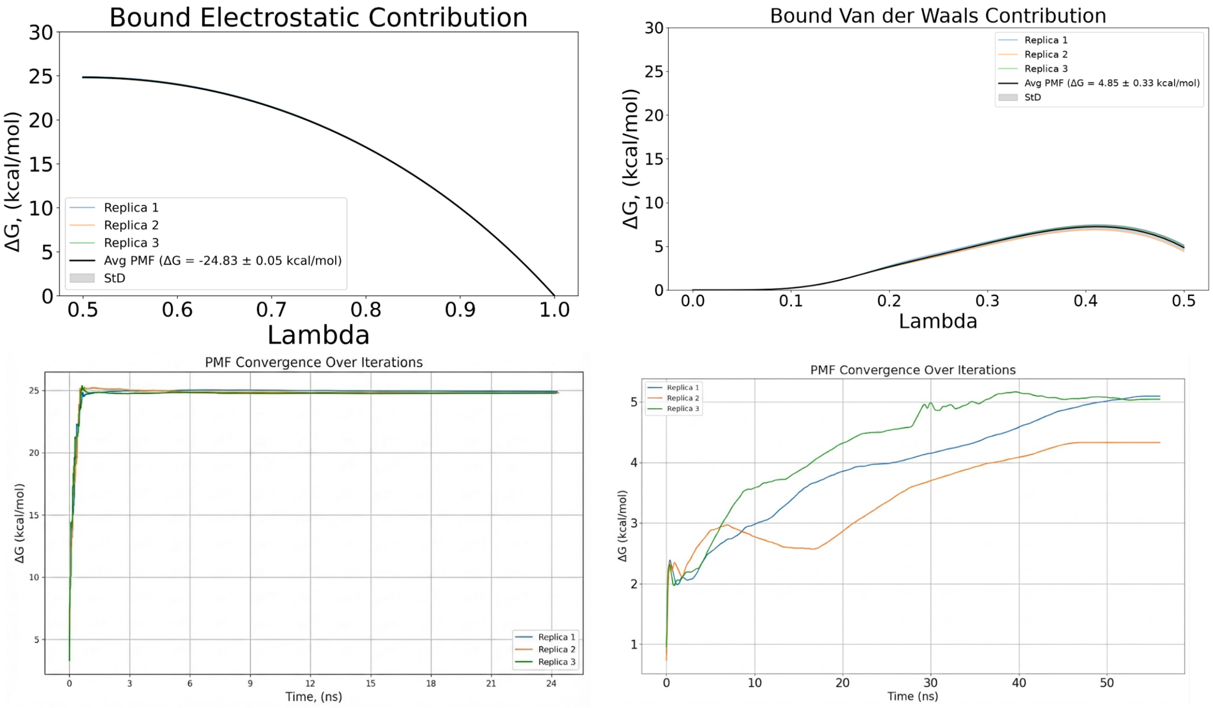


Figure S24. PMF and convergence plots of bound electrostatic and van der Waals energetic contribution for three replicas for MUP-I complex (PDB: 1I05).

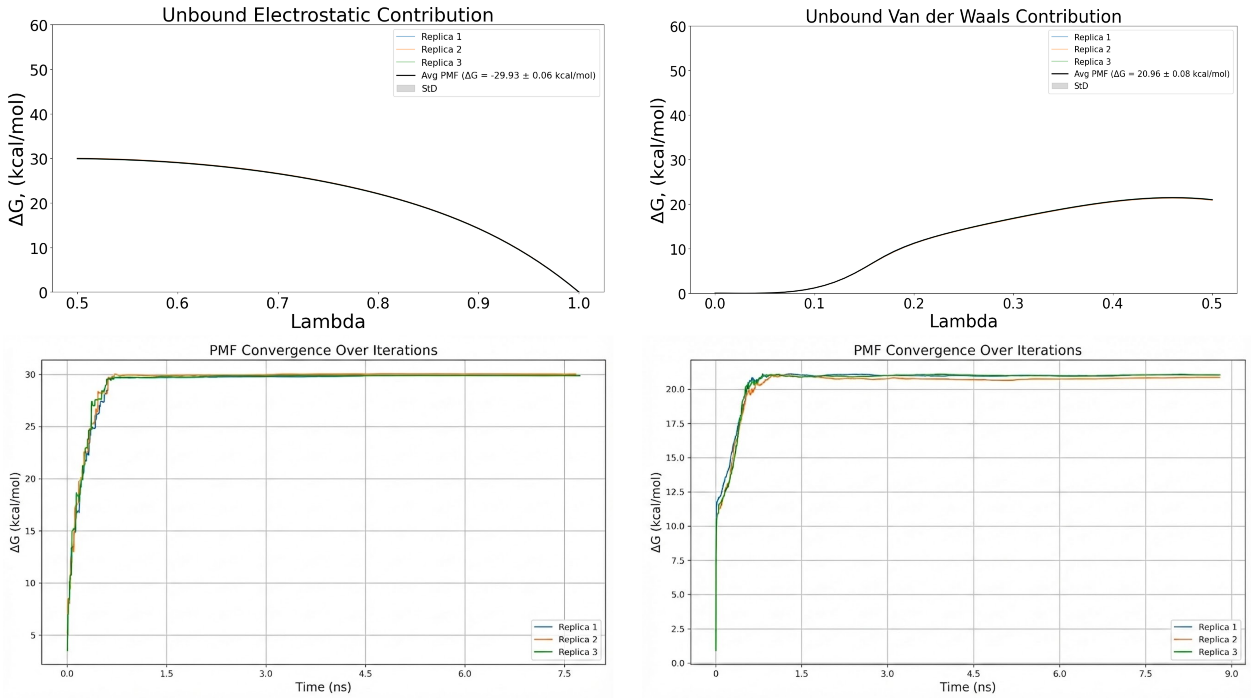


Figure S25. PMF and convergence plots of unbound electrostatic and van der Waals energetic contribution for three replicas for MUP-I complex (PDB: 1I05).

#### $\lambda$-Conditioned Hydration and Crystallographic Site Occupancy in the 1I05 Complex

To characterize both the extent and spatial localization of hydration sampled during the bound-state alchemical transformation, we performed complementary analyses of the TYR138-centered hydration environment and the crystallographic W1 and W2 water sites. The TYR138-centered analysis was applied to three sampling protocols, whereas the W1/W2 analysis was used as a site-localization diagnostic for the production Lambda-ABF-OPES trajectories.

**Sampling protocols and analyzed trajectories.** The hydration analysis was performed for three bound-state complex-leg protocols constructed using the same 1I05 molecular system, AMOEBA polarizable force field, alchemical leg definitions, and DBC restraint:

1. the production continuous Lambda-ABF-OPES calculation;
2. a continuous Lambda-ABF calculation in which the OPES contribution was removed, hereafter denoted *No-OPES*;
3. a narrow-window dynamic-$\lambda$ Lambda-ABF-OPES control, hereafter denoted the *narrow-window control*.

The production Lambda-ABF-OPES protocol allows the dynamical alchemical coordinate to diffuse continuously over the complete interval associated with each alchemical leg. The multiple-walker ABF contribution estimates and compensates the mean force along $\lambda$, while OPES provides an additional adaptive bias designed to enhance exploration of under-sampled regions of the alchemical coordinate.

The No-OPES calculation retains the AMOEBA force field, the continuous dynamical alchemical coordinate, the multiple-walker ABF contribution, and the DBC restraint, but excludes the OPES bias. It therefore isolates the incremental effect of OPES relative to the underlying continuous Lambda-ABF protocol without changing the molecular interaction model. The production

Lambda-ABF-OPES and No-OPES trajectories were compared over the same analyzed trajectory length.

For the narrow-window control, the alchemical path was divided into intervals of width

$$\Delta\lambda = 0.025. \tag{10}$$

The VDW leg was represented by intervals spanning $\lambda = 0.000$–$0.500$, and the electrostatic leg by intervals spanning $\lambda = 0.500$–$1.000$. Each interval was sampled independently for 2 ns. Reflecting lower and upper boundaries confined the extended $\lambda$ coordinate to the prescribed interval.

In extended-Lagrangian alchemical dynamics, the motion of the dynamical coordinate can be written schematically as

$$M_\lambda \ddot{\lambda} = F_\lambda - \gamma_\lambda \dot{\lambda} + \eta_\lambda(t), \tag{11}$$

where $M_\lambda$ is the extended mass, $F_\lambda$ is the effective force acting on the alchemical coordinate, $\gamma_\lambda$ is the friction coefficient, and $\eta_\lambda(t)$ is the corresponding thermostat noise. For the narrow-window control, we used

$$M_\lambda = 1.5 \times 10^9, \tag{12}$$

which strongly reduces the acceleration and mobility of the extended coordinate. Together, the large extended mass and the reflecting boundaries produce quasi-local sampling within each narrow $\lambda$ interval.

Because $\lambda$ remains a dynamical variable and continues to fluctuate within each prescribed interval, these simulations are referred to as narrow-window dynamic-$\lambda$ controls rather than exact fixed-$\lambda$ simulations. Analysis of the sampled $\lambda$ trajectories confirmed the intended confinement: the sampled values remained within their assigned boundaries, with mean within-window standard deviations of approximately 0.0073 in the VDW leg and 0.0072 in the electrostatic leg.

The narrow-window protocol therefore tests whether independent local sampling along the alchemical path recovers the same thermodynamic and structural ensemble as continuous exploration of the complete alchemical interval. These calculations constitute a mechanistic local-sampling control and are not interpreted as a total-computational-cost-matched comparison with the continuous Lambda-ABF-OPES and No-OPES trajectories.

**Free-energy profiles obtained with the three sampling protocols.** For the narrow-window calculations, the absolute PMF offset within an individual interval is arbitrary. We therefore retained the invariant free-energy difference across each interval:

$$\Delta G_i = G_i(\lambda_{\text{upper}}) - G_i(\lambda_{\text{lower}}), \tag{13}$$

and reconstructed the cumulative profile according to

$$G_{\text{windowed}}(\lambda_n) = \sum_{i=1}^{n} \Delta G_i. \tag{14}$$

All profiles were shifted to zero at the beginning of the corresponding alchemical leg. The resulting endpoint contributions are summarized in Table S5, and the full profiles are shown in Fig. S26.

**Table S5. Endpoint free-energy contributions obtained with the three sampling protocols for the bound-state complex leg of 1I05. All profiles were shifted to zero at the beginning of the corresponding alchemical leg.**

| Protocol | ELEC contribution ($\text{kcal mol}^{-1}$) | VDW contribution ($\text{kcal mol}^{-1}$) |
|---|---|---|
| Narrow-window control | $-24.95$ | $+2.22$ |
| Lambda-ABF-OPES | $-24.90$ | $+3.58$ |
| No-OPES | $-24.78$ | $+3.24$ |

The electrostatic contribution is essentially protocol-independent: the three endpoints differ by less than $0.2\ \text{kcal mol}^{-1}$. By contrast, the VDW contribution is more sensitive to the sampling protocol. The Lambda-ABF-OPES and No-OPES calculations yield $+3.58$ and $+3.24\ \text{kcal mol}^{-1}$, respectively, whereas the narrow-window control yields $+2.22\ \text{kcal mol}^{-1}$. The modest difference between Lambda-ABF-OPES and No-OPES indicates that OPES provides an incremental contribution rather than being the sole origin of the VDW response. The larger difference relative to the narrow-window control suggests that continuous exploration of the full alchemical interval

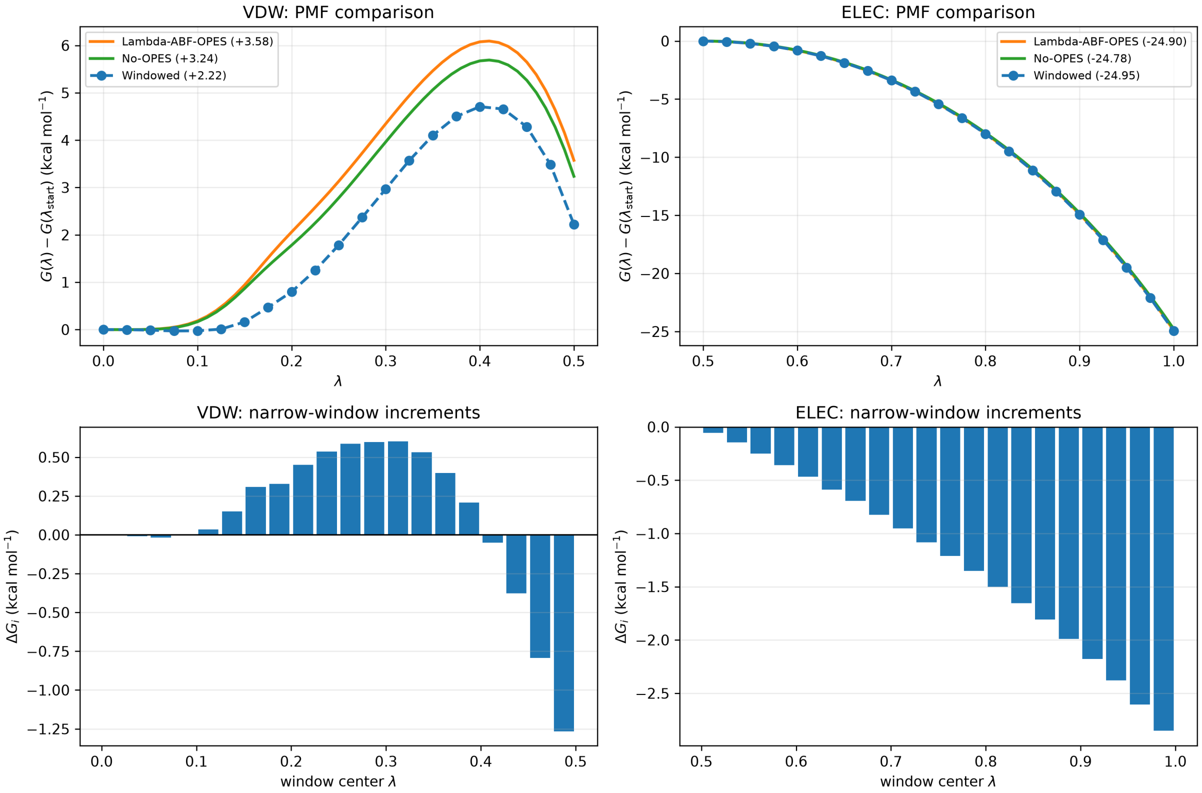


Figure S26. Comparison of the 1I05 bound-state PMFs obtained with the continuous Lambda-ABF-OPES calculation, the continuous No-OPES control, and the narrow-window dynamic-$\lambda$ control. The upper panels show the cumulative VDW and electrostatic PMFs, whereas the lower panels show the local narrow-window free-energy increments, $\Delta G_i$. The electrostatic profiles and endpoints are closely reproduced by all three protocols. The VDW leg displays greater protocol dependence, with the narrow-window control yielding a smaller endpoint contribution than the two continuous-$\lambda$ calculations.

facilitates relaxation of slower degrees of freedom coupled to steric decoupling.

Because each narrow window was sampled for 2 ns, this result is interpreted as evidence of protocol sensitivity over the investigated timescale. It does not imply that the narrow-window estimator is intrinsically biased or that substantially longer sampling could not recover the same endpoint.

The narrow-window VDW increments also change sign along the alchemical path, whereas the electrostatic contribution accumulates more smoothly. This difference indicates that the VDW transformation is coupled to a more complex combination of cavity formation, ligand relaxation, and solvent reorganization. We therefore examined whether the PMF differences were associated with changes in the restrained ligand-pose or ligand-conformational ensembles.

**Restrained ligand-pose and conformational controls.** The DBC coordinate was analyzed as a function of $\lambda$ for all three protocols (Fig. S27). The sampled DBC envelopes are broadly comparable, indicating that the protocol-dependent VDW differences do not arise from ligand escape or gross displacement outside the restrained bound-state region.

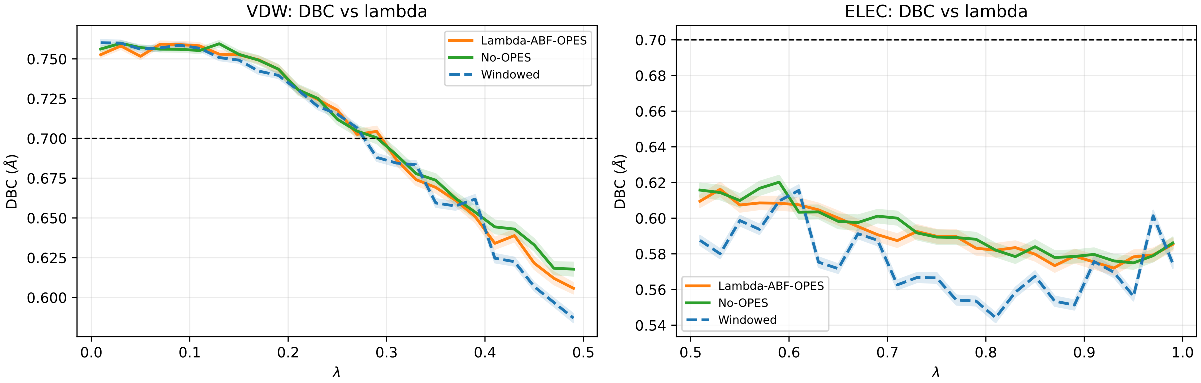


Figure S27. DBC coordinate as a function of the alchemical coordinate for Lambda-ABF-OPES, No-OPES, and the narrow-window control. The comparable DBC envelopes show that the protocol-dependent VDW PMF differences are not caused by gross ligand displacement or loss of the restrained bound-state ensemble.

A representative ligand torsion was also compared across the three protocols (Fig. S28). The torsional distributions remain multimodal and contain the same principal basins. Differences in their relative populations are present, but no protocol loses a major ligand conformational state. The VDW PMF difference therefore cannot be attributed simply to a qualitatively different ligand conformation.

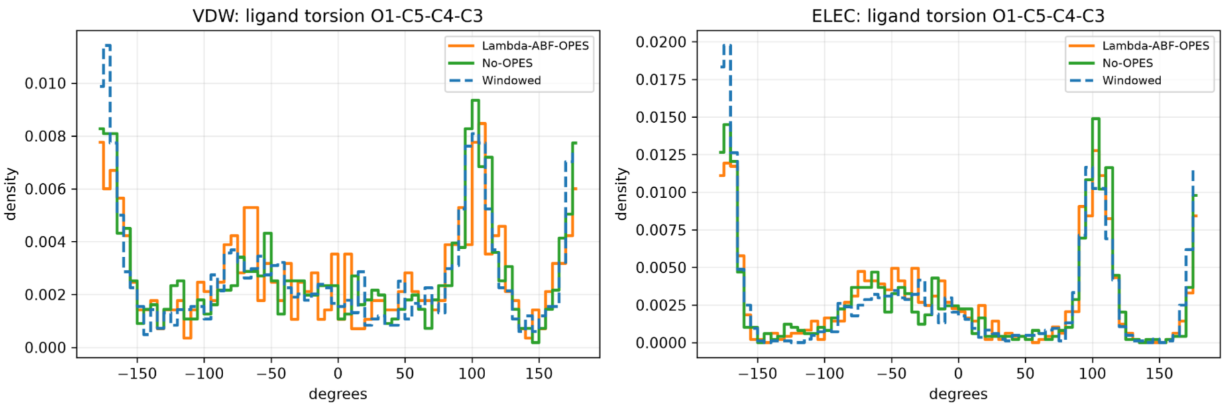


Figure S28. Distributions of the representative ligand torsion during the VDW and electrostatic legs for Lambda-ABF-OPES, No-OPES, and the narrow-window control. All protocols sample the principal torsional basins, indicating that the protocol-dependent VDW response is not caused by loss of a major ligand conformational state.

The PMF and structural controls therefore localize the principal protocol-dependent difference to the VDW transformation while excluding gross ligand displacement and loss of a major torsional basin as dominant causes. We next examined whether the remaining difference was associated with local solvent reorganization around the buried TYR138-centered hydration network.

**Continuous TYR138-OH hydration coordinate.** TYR138 was selected because its hydroxyl oxygen is a principal polar anchor of the buried water-mediated network in the 1I05 binding site. Water contacts were described using the smooth switching function

$$s(d) = \frac{1}{1 + \exp\left[(d - d_0)/\sigma\right]}, \tag{15}$$

where $d$ is a water–atom distance, $d_0 = 4.0$ Å is the midpoint of the switching function, and $\sigma = 0.15$ Å controls the sharpness of the transition. Distances well below $4.0$ Å therefore contribute approximately one, distances well above this value contribute approximately zero, and waters fluctuating near the cutoff are treated continuously.

For each saved configuration $t$, the continuous hydration of the TYR138 hydroxyl oxygen was calculated as

$$n_{\mathrm{TYR138OH}}(t) = \sum_{w \in W} s\left(r_{w,\mathrm{TYR138OH}}(t)\right), \tag{16}$$

where $W$ is the set of all water oxygen atoms and $r_{w,\mathrm{TYR138OH}}(t)$ is the distance between water oxygen $w$ and the TYR138 hydroxyl oxygen at frame $t$. Thus, $n_{\mathrm{TYR138OH}}$ reports the continuous number of waters hydrating TYR138 OH. Because it is evaluated directly from instantaneous atom–atom distances, this coordinate does not require assignment of a particular water molecule to a fixed crystallographic site.

**$\lambda$-conditioned hydration observables.** The three protocols accumulate different numbers of frames in different regions of the alchemical coordinate. Hydration was therefore analyzed conditionally on the instantaneous value of $\lambda$, using intervals of width

$$\Delta\lambda = 0.025.$$

For an interval centered at $\lambda_k$, the conditional mean hydration was calculated as

$$\langle n_{\mathrm{TYR138OH}} \rangle_{\lambda_k} = \frac{1}{N(\lambda_k)} \sum_{t \in \lambda_k} n_{\mathrm{TYR138OH}}(t), \tag{17}$$

where $N(\lambda_k)$ is the number of analyzed configurations in that interval. Each interval is therefore normalized by its own frame population, so the resulting observable reports hydration at a given alchemical coupling rather than the raw sampling population along $\lambda$.

As a complementary state-based measure, we calculated the conditional probability of a higher-hydration TYR138-OH state:

$$P^{(\geq 1.5)}_{\mathrm{TYR138OH}}(\lambda_k) = \frac{N\left(n_{\mathrm{TYR138OH}} \geq 1.5, \lambda_k\right)}{N(\lambda_k)}. \tag{18}$$

The threshold $n_{\mathrm{TYR138OH}} \geq 1.5$ separates the dominant approximately one-water hydration state from configurations containing additional water density around TYR138 OH. For visual representation, the resulting $\lambda$-resolved probability was smoothed along $\lambda$ using the same Gaussian bandwidth for all three protocols. This smoothing was applied only to reduce bin-to-bin noise and does not alter the underlying state definition.

To retain the complete hydration-state distribution, rather than reduce it to a mean value or a single threshold, we additionally constructed the conditional density

$$\widehat{p}\left(n_{\mathrm{TYR138OH}} \mid \lambda_k\right) = \frac{\widehat{p}\left(\lambda_k, n_{\mathrm{TYR138OH}}\right)}{\int \widehat{p}\left(\lambda_k, n'\right)\, dn'}. \tag{19}$$

The framewise pairs $\{\lambda(t), n_{\mathrm{TYR138OH}}(t)\}$ were accumulated into a two-dimensional histogram and smoothed using Gaussian bandwidths

$$h_\lambda = 0.030, \qquad h_n = 0.10.$$

Following smoothing, every $\lambda$ slice was renormalized such that

$$\int \widehat{p}\left(n_{\mathrm{TYR138OH}} \mid \lambda_k\right)\, dn_{\mathrm{TYR138OH}} = 1. \tag{20}$$

This procedure removes the direct influence of unequal sampling populations along $\lambda$. Because the adaptive biases depend on $\lambda$, rather than directly on $n_{\mathrm{TYR138OH}}$, no additional reweighting with respect to the hydration coordinate was applied.

For each protocol, the contours in Fig. S29 represent nested regions containing 50%, 75%, and 90% of the corresponding $\lambda$-balanced conditional distribution. More explicitly, the density threshold $c_\alpha$ for probability mass $\alpha$ was selected such that

$$\frac{1}{K}\sum_{k=1}^{K}\int_{\widehat{p}(n\mid\lambda_k)\geq c_\alpha} \widehat{p}(n \mid \lambda_k)\, dn = \alpha, \qquad \alpha \in \{0.50, 0.75, 0.90\}, \tag{21}$$

where $K$ is the number of populated $\lambda$ intervals. Thus, every $\lambda$ interval contributes equally to the displayed distribution. The innermost contour encloses the most highly populated hydration states, whereas the intermediate and outer contours progressively include less populated states and describe the broader hydration ensemble.

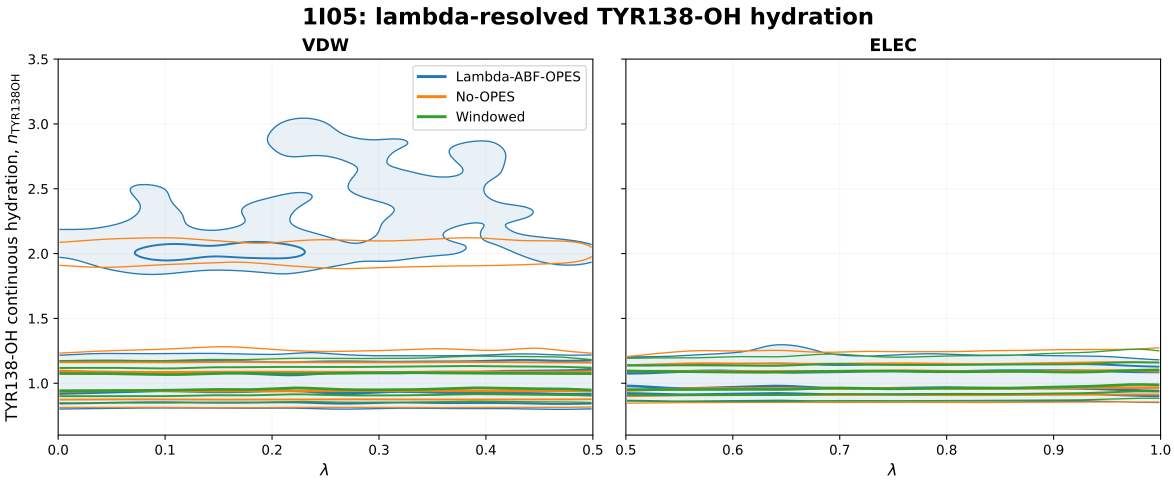


Figure S29. Conditional TYR138-OH hydration distributions for the 1I05 complex during the VDW and electrostatic alchemical legs. The plotted quantity, $\widehat{p}(n_{\mathrm{TYR138OH}} \mid \lambda)$, was normalized independently within each $\lambda$ slice. Blue, orange, and green contours enclose 50%, 75%, and 90% of the corresponding $\lambda$-balanced distributions for Lambda-ABF-OPES, No-OPES, and the narrow-window control, respectively. The outer Lambda-ABF-OPES region is lightly shaded for visual guidance. Closely spaced contours indicate a narrowly localized hydration state, whereas contours extending over a larger range of $n_{\mathrm{TYR138OH}}$ indicate access to a broader hydration ensemble.

All three protocols retain the dominant TYR138-OH hydration state near $n_{\mathrm{TYR138OH}} \approx 1$. The principal protocol-dependent difference appears during the VDW transformation. The Lambda-ABF-OPES distribution extends toward $n_{\mathrm{TYR138OH}} \approx 2$–3, indicating access to additional higher-hydration configurations. The No-OPES distribution shows a more limited extension toward $n_{\mathrm{TYR138OH}} \approx 2$, whereas the narrow-window distribution remains predominantly concentrated near the lower-hydration state. In the electrostatic leg, the three contour envelopes largely overlap and remain localized near the approximately one-water state.

The conditional mean and higher-hydration probability provide scalar quantification of the same distributions (Figs. S30 and S31).

For an integrated numerical comparison, we averaged the conditional observables over the populated $\lambda$ intervals, assigning equal weight to every interval. The resulting values are reported

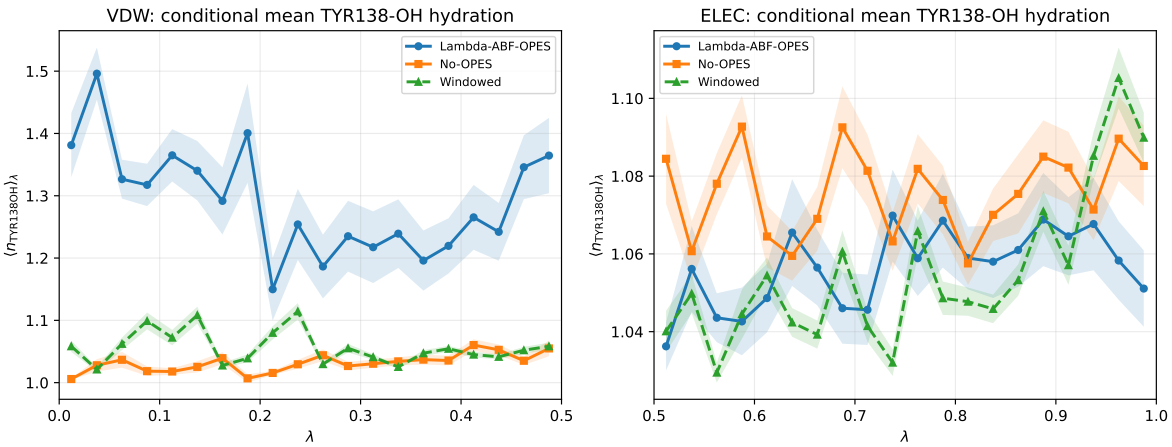


Figure S30. Conditional mean TYR138-OH hydration for the 1I05 complex. The quantity $\langle n_{\mathrm{TYR138OH}} \rangle_\lambda$ was calculated according to Eq. 17, with each $\lambda$ interval normalized by its own number of frames. During the VDW transformation, Lambda-ABF-OPES samples a higher mean hydration level than the No-OPES and narrow-window controls. The electrostatic-leg profiles remain substantially closer.

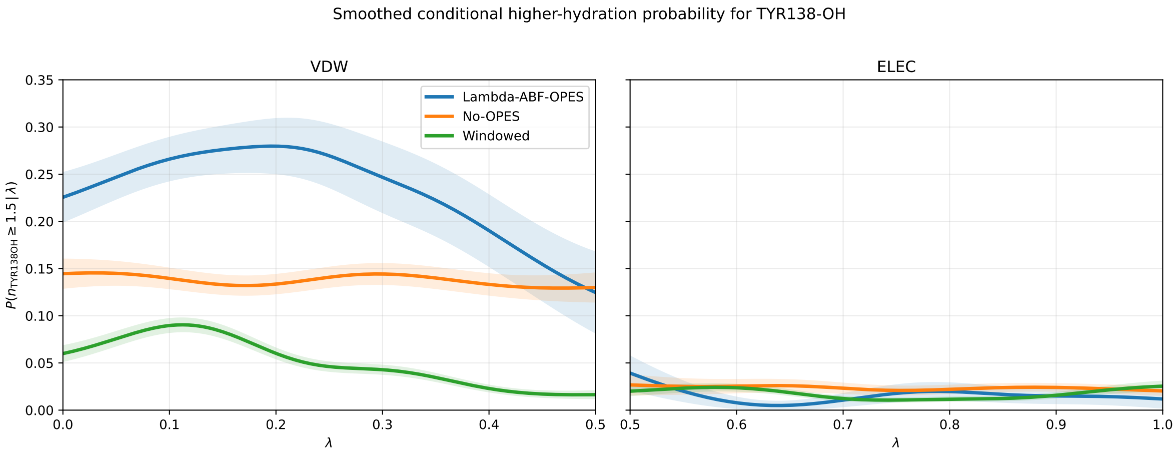


Figure S31. Conditional probability of the higher-hydration TYR138-OH state, defined by $n_{\mathrm{TYR138OH}} \geq 1.5$. The plotted quantity was calculated according to Eq. 18 and represented as a smoothed local average along $\lambda$. During the VDW transformation, Lambda-ABF-OPES samples the higher-hydration state more frequently than either the No-OPES or narrow-window control. In the electrostatic leg, this state remains weakly populated for all three protocols.

in Table S6.

**Table S6. $\lambda$-balanced TYR138-OH hydration statistics for the three sampling protocols. Every populated $\Delta\lambda = 0.025$ interval contributes equally to the reported averages.**

| | VDW leg | | Electrostatic leg | |
|---|---|---|---|---|
| Protocol | $\langle n_{\mathrm{TYR138OH}}\rangle_\lambda$ | $P(n_{\mathrm{TYR138OH}} \geq 1.5)$ | $\langle n_{\mathrm{TYR138OH}}\rangle_\lambda$ | $P(n_{\mathrm{TYR138OH}} \geq 1.5)$ |
| Lambda-ABF-OPES | 1.315 | 0.231 | 1.057 | 0.016 |
| No-OPES | 1.182 | 0.138 | 1.068 | 0.025 |
| Narrow-window | 1.053 | 0.027 | 1.053 | 0.010 |

The numerical results confirm that the principal difference is localized to the VDW leg. Lambda-ABF-OPES gives both the largest conditional mean hydration and the largest probability of the higher-hydration state. By contrast, the three electrostatic-leg values remain similar, and the higher-hydration state is only weakly populated in all three protocols. Because these observables are conditioned on $\lambda$, the differences cannot be attributed simply to protocol-dependent accumulation of frames along the alchemical coordinate.

**Crystallographic W1/W2 site occupancy.** The TYR138-centered analysis quantifies the extent and breadth of local hydration but does not, by itself, establish whether the sampled waters remain localized within the crystallographic W1 and W2 regions. We therefore performed a complementary site-resolved analysis for the production Lambda-ABF-OPES trajectories.

For each saved configuration, the binding-site region was locally aligned to the reference structure to remove global translation and rotation. The distance between every water oxygen and the reference oxygen position of each crystallographic hydration site was then evaluated. For $i \in \{\mathrm{W1}, \mathrm{W2}\}$, the instantaneous site occupancy was defined from the closest water oxygen:

$$o_i(t) = \frac{1}{1 + \exp\left[\left(d_i^{\min}(t) - d_{\mathrm{site}}\right)/\sigma_{\mathrm{site}}\right]}, \tag{22}$$

where

$$d_i^{\min}(t) = \min_{w \in W} \left\| \mathbf{r}_w(t) - \mathbf{R}_i^{\mathrm{ref}} \right\|. \tag{23}$$

Here, $\mathbf{R}_i^{\mathrm{ref}}$ is the reference oxygen position of site $i$, $d_{\mathrm{site}} = 2.0$ Å is the site-localization distance, and $\sigma_{\mathrm{site}} = 0.15$ Å provides a narrow continuous transition around this value. Thus, $o_i(t)$ approaches one when a water oxygen is localized within the corresponding crystallographic site and approaches zero when no water occupies that region.

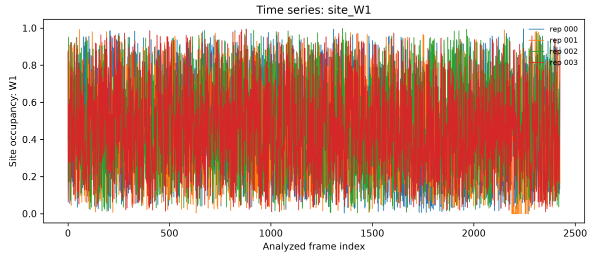


(a) Electrostatic leg: W1 occupancy.

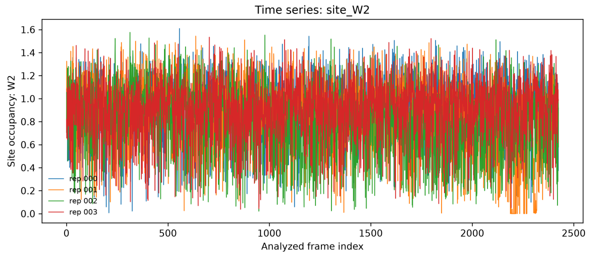


(b) Electrostatic leg: W2 occupancy.

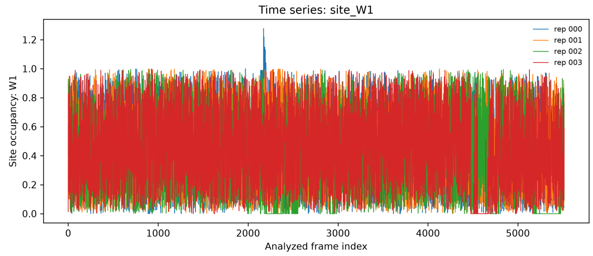


(c) VDW leg: W1 occupancy.

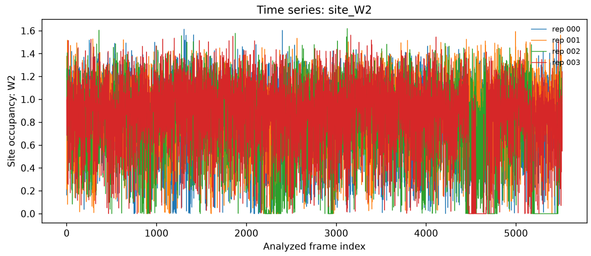


(d) VDW leg: W2 occupancy.

Figure S32. Time-resolved occupancies of the crystallographic W1 and W2 hydration sites during the electrostatic and VDW legs of the bound-state Lambda-ABF-OPES calculation for 1I05. Following local alignment of the binding-site region, each occupancy was calculated from the distance between the corresponding reference site and the closest water oxygen using the smooth switching function in Eq. 22. Values approaching one indicate localization of a water oxygen within the crystallographic site, whereas values approaching zero indicate temporary vacancy. The traces show repeated occupation of the W1 and W2 regions during both alchemical legs and provide spatial localization evidence complementary to the TYR138-centered coordination analysis.

The W1/W2 traces show that waters sampled during the production Lambda-ABF-OPES calculation repeatedly occupy the crystallographic buried-water regions, with temporary vacancy and subsequent replacement occurring during the trajectories. The TYR138-centered signal therefore does not arise solely from diffuse water accumulation near the ligand or protein surface; it is associated with water localization within the experimentally defined buried hydration environment.

**Integrated interpretation.** Together, the four analyses provide complementary information. The conditional density describes the range of TYR138-centered hydration states sampled at a given $\lambda$, the conditional mean and threshold probability quantify the protocol-dependent population of these states, and the W1/W2 occupancy traces establish their spatial relationship to the crystallographic buried-water region.

The dominant approximately one-water TYR138 hydration state is recovered by all three protocols. The clearest difference occurs during the VDW transformation, where Lambda-ABF-OPES samples a broader and more highly hydrated TYR138-centered ensemble than the No-OPES and narrow-window controls. The electrostatic-leg distributions remain substantially more similar, indicating that the persistent polar hydration motif is robust to the sampling protocol. The W1/W2 occupancy traces further show that the waters sampled in the production calculation remain localized within the known buried hydration network.

These results are interpreted as state-resolved structural sampling diagnostics. They demonstrate protocol-dependent differences in the hydration configurations sampled during steric decoupling and verify the spatial localization of the production hydration signal. They do not constitute an independent absolute hydration free-energy surface or, by themselves, establish thermodynamic exactness of hydration recovery.

# 5i40

The labels O/O, N/O, O/N, and N/N denote the heavy-atom element classes used for contact classification, with the first element corresponding to the protein atom and the second to the ligand atom. Residue labels in the figures are used as compact notation. The corresponding atom-level assignments, including protein atom name and backbone versus side-chain origin, are reported in Table S7

**Table S7. Atom-level direct and water-mediated polar contacts for 5i40. Motif labels are reported as protein-element/ligand-element. Direct contacts use a $4.0$ Å heavy-atom cutoff. Water-mediated contacts require both ligand–water and water–protein distances to be within $4.0$ Å, and ligand–protein proximity $\leq 6.0$ Å.**

| contact type | motif | contact label | atom origin | occ. fraction | mean events | direct dist Å | lig–wat dist Å | wat–prot dist Å | total path Å | unique water O |
|---|---|---|---|---|---|---|---|---|---|---|
| direct | O/N | 67N206:N1–ASN100:OD1 (side chain) | side chain | 0.929 | 1.00 | 3.64 | | | | |
| direct | O/N | 67N206:N2–VAL49:O (backbone) | backbone | 0.610 | 1.00 | 3.31 | | | | |
| direct | O/O | 67N206:O–VAL49:O (backbone) | backbone | 0.550 | 1.00 | 3.26 | | | | |
| direct | O/N | 67N206:N1–TYR106:OH (side chain) | side chain | 0.417 | 1.00 | 3.72 | | | | |
| direct | O/N | 67N206:N1–ALA54:O (backbone) | backbone | 0.407 | 1.00 | 3.38 | | | | |
| direct | O/N | 67N206:N1–TYR106:O (backbone) | backbone | 0.221 | 1.00 | 3.71 | | | | |
| direct | O/O | 67N206:O–TYR106:O (backbone) | backbone | 0.110 | 1.00 | 3.65 | | | | |
| direct | O/O | 67N206:O–ILE53:O (backbone) | backbone | 0.105 | 1.00 | 3.70 | | | | |
| direct | O/O | 67N206:O–ALA54:O (backbone) | backbone | 0.058 | 1.00 | 3.51 | | | | |
| direct | O/O | 67N206:O–TYR106:OH (side chain) | side chain | 0.048 | 1.00 | 3.75 | | | | |
| direct | O/N | 67N206:N2–ALA54:O (backbone) | backbone | 0.029 | 1.00 | 3.67 | | | | |
| direct | O/N | 67N206:N2–TYR106:O (backbone) | backbone | 0.012 | 1.00 | 3.82 | | | | |
| direct | O/N | 67N206:N2–ALA96:O (backbone) | backbone | 0.012 | 1.00 | 3.89 | | | | |

*Continued on next page*

#### Table S7. Atom-level direct and water-mediated polar contacts for 5i40 (continued)

| contact type | motif | contact label | atom origin | occ. fraction | mean events | direct dist Å | lig–wat dist Å | wat–prot dist Å | total path Å | unique water O |
|---|---|---|---|---|---|---|---|---|---|---|
| direct | O/N | 67N206:N1–ILE53:O (backbone) | backbone | 0.010 | 1.00 | 3.63 | | | | |
| water-mediated | O/N | 67N206:N1–W(O)–ASN100:OD1 (side chain) | side chain | 0.371 | 1.02 | | 3.68 | 2.87 | 6.55 | 333 |
| water-mediated | O/N | 67N206:N1–W(O)–TYR106:OH (side chain) | side chain | 0.303 | 1.03 | | 3.65 | 3.09 | 6.74 | 775 |
| water-mediated | O/N | 67N206:N1–W(O)–TYR106:O (backbone) | backbone | 0.277 | 1.04 | | 3.66 | 3.30 | 6.96 | 765 |
| water-mediated | O/O | 67N206:O–W(O)–ILE53:O (backbone) | backbone | 0.269 | 1.11 | | 3.58 | 3.41 | 6.99 | 372 |
| water-mediated | O/O | 67N206:O–W(O)–VAL49:O (backbone) | backbone | 0.262 | 1.05 | | 3.65 | 3.28 | 6.93 | 218 |
| water-mediated | O/N | 67N206:N1–W(O)–TYR99:OH (side chain) | side chain | 0.238 | 1.03 | | 3.63 | 3.25 | 6.88 | 683 |
| water-mediated | O/N | 67N206:N1–W(O)–PRO55:O (backbone) | backbone | 0.236 | 1.14 | | 3.61 | 3.08 | 6.69 | 719 |
| water-mediated | O/N | 67N206:N2–W(O)–TYR57:O (backbone) | backbone | 0.178 | 1.00 | | 3.87 | 1.81 | 5.67 | 3 |
| water-mediated | O/N | 67N206:N2–W(O)–TYR57:OH (side chain) | side chain | 0.172 | 1.00 | | 3.86 | 2.71 | 6.57 | 3 |
| water-mediated | O/O | 67N206:O–W(O)–TYR106:O (backbone) | backbone | 0.144 | 1.06 | | 3.63 | 2.45 | 6.07 | 232 |
| water-mediated | N/O | 67N206:O–W(O)–VAL49:N (backbone) | backbone | 0.124 | 1.06 | | 3.70 | 3.69 | 7.40 | 164 |
| water-mediated | O/N | 67N206:N1–W(O)–TYR99:O (backbone) | backbone | 0.113 | 1.04 | | 3.63 | 3.34 | 6.96 | 359 |
| water-mediated | O/N | 67N206:N2–W(O)–VAL49:O (backbone) | backbone | 0.113 | 1.02 | | 3.86 | 3.26 | 7.12 | 30 |
| water-mediated | O/O | 67N206:O–W(O)–TYR106:OH (side chain) | side chain | 0.110 | 1.01 | | 3.66 | 2.80 | 6.46 | 132 |
| water-mediated | O/N | 67N206:N1–W(O)–ALA54:O (backbone) | backbone | 0.084 | 1.02 | | 3.55 | 3.58 | 7.14 | 293 |
| water-mediated | O/N | 67N206:N2–W(O)–ALA96:O (backbone) | backbone | 0.072 | 1.00 | | 3.86 | 3.33 | 7.19 | 4 |
| water-mediated | O/N | 67N206:N1–W(O)–THR104:O (backbone) | backbone | 0.029 | 1.10 | | 3.67 | 2.97 | 6.64 | 90 |

*Continued on next page*

#### Table S7. Atom-level direct and water-mediated polar contacts for 5i40 (continued)

| contact type | motif | contact label | atom origin | occ. fraction | mean events | direct dist Å | lig–wat dist Å | wat–prot dist Å | total path Å | unique water O |
|---|---|---|---|---|---|---|---|---|---|---|
| water-mediated | N/O | 67N206:O–W(O)–PHE45:N (backbone) | backbone | 0.023 | 1.00 | | 3.76 | 3.91 | 7.67 | 11 |
| water-mediated | N/O | 67N206:O–W(O)–THR50:N (backbone) | backbone | 0.019 | 1.00 | | 3.65 | 3.31 | 6.96 | 43 |
| water-mediated | N/N | 67N206:N2–W(O)–VAL49:N (backbone) | backbone | 0.018 | 1.00 | | 3.83 | 3.66 | 7.49 | 8 |
| water-mediated | N/N | 67N206:N1–W(O)–ALA54:N (backbone) | backbone | 0.017 | 1.00 | | 3.58 | 3.86 | 7.45 | 91 |
| water-mediated | O/N | 67N206:N1–W(O)–ILE53:O (backbone) | backbone | 0.012 | 1.07 | | 3.67 | 3.36 | 7.04 | 57 |

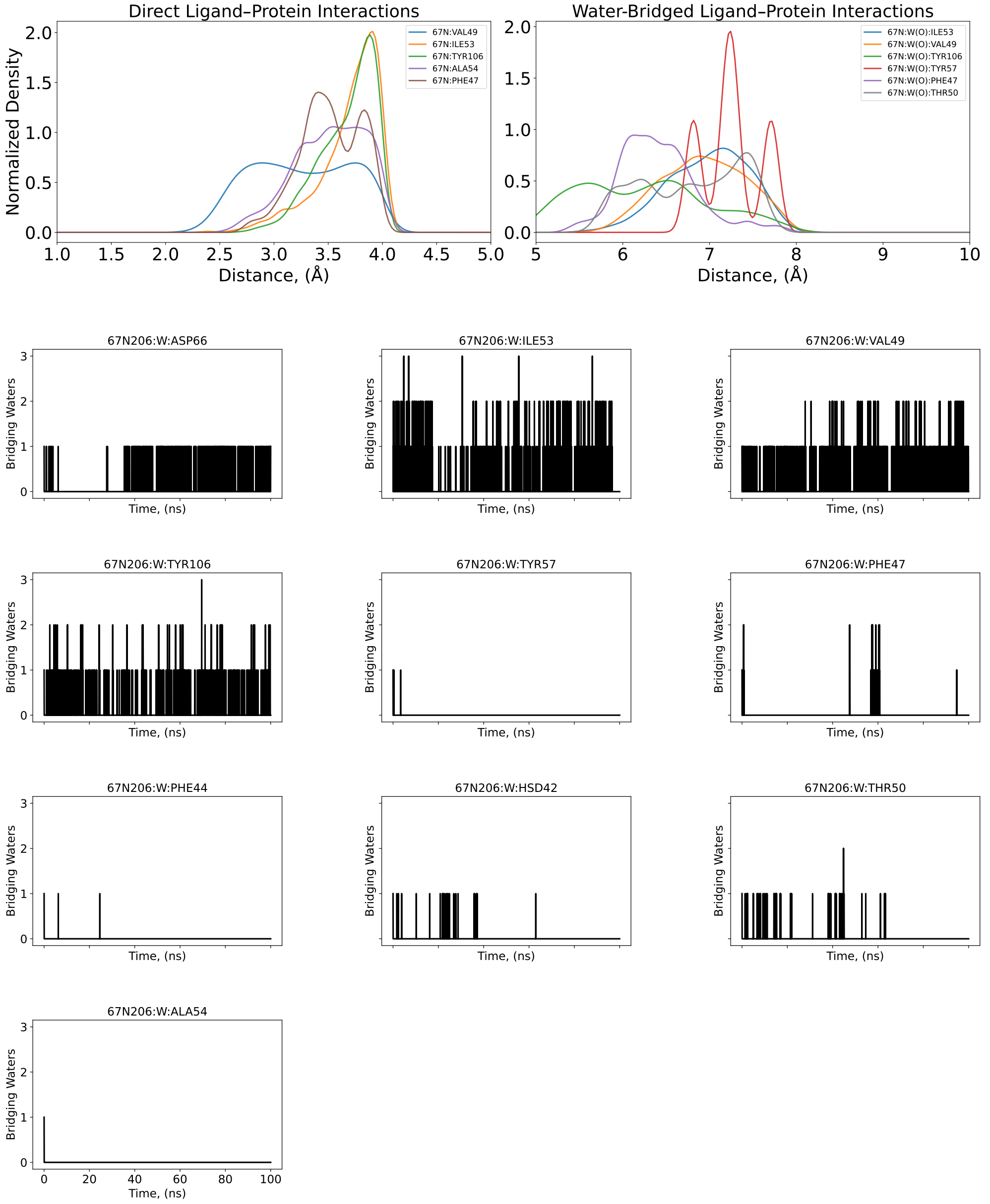


Figure S33. 4-Å–density distribution, averaged and individual coordination number of bridging oxygen water molecules (O-bridging) that mediate individual oxygen atoms of the protein and oxygen atoms of the ligand along a 100-ns conventional MD for BRD9 complex (PDB: 5I40).

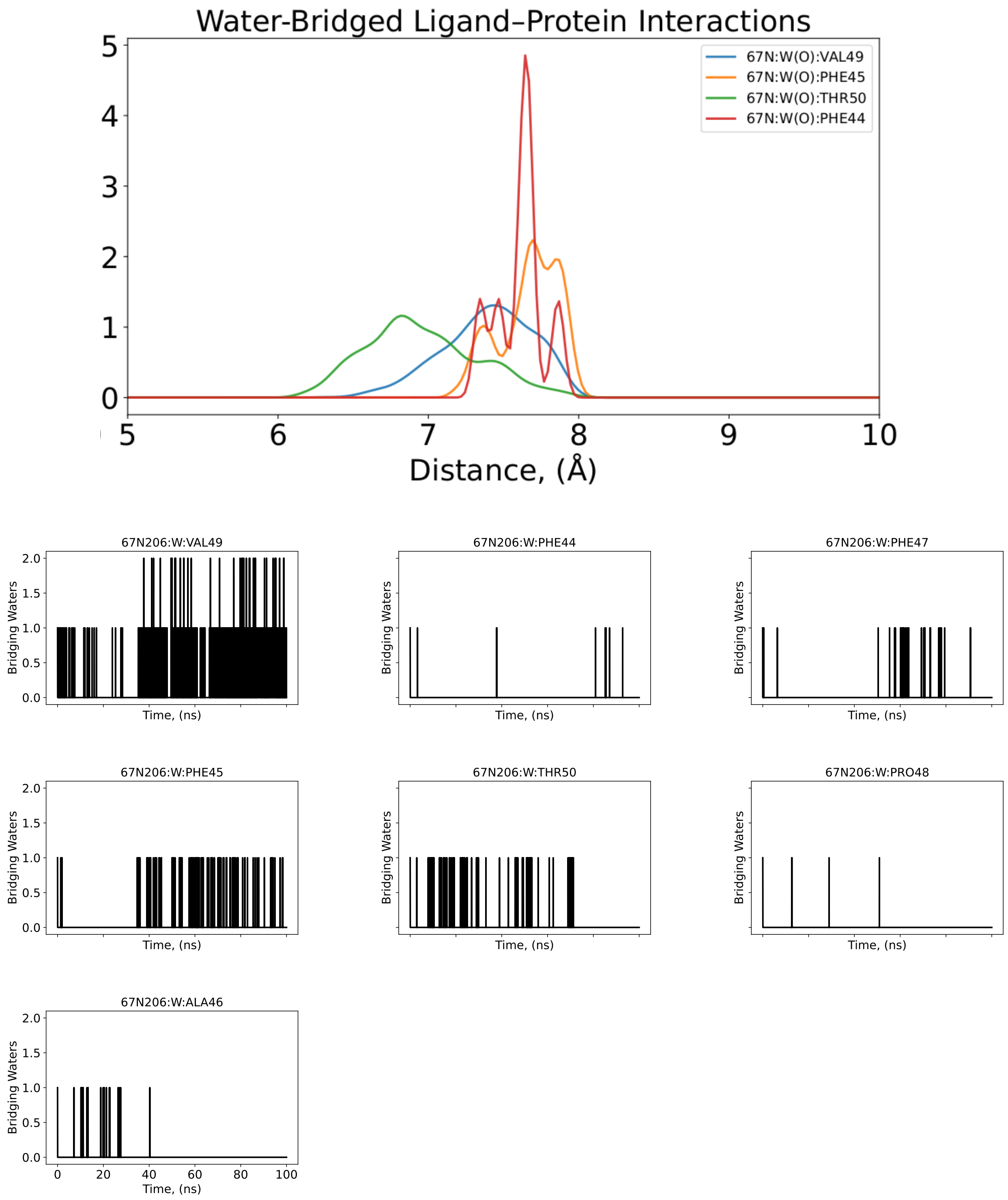


Figure S34. 4-Å–density distribution and individual coordination number of bridging oxygen water molecules (N/O-bridging) that mediate individual nitrogen atoms of the protein and oxygen atoms of the ligand along a 100-ns conventional MD for BRD9 complex (PDB: 5I40) .

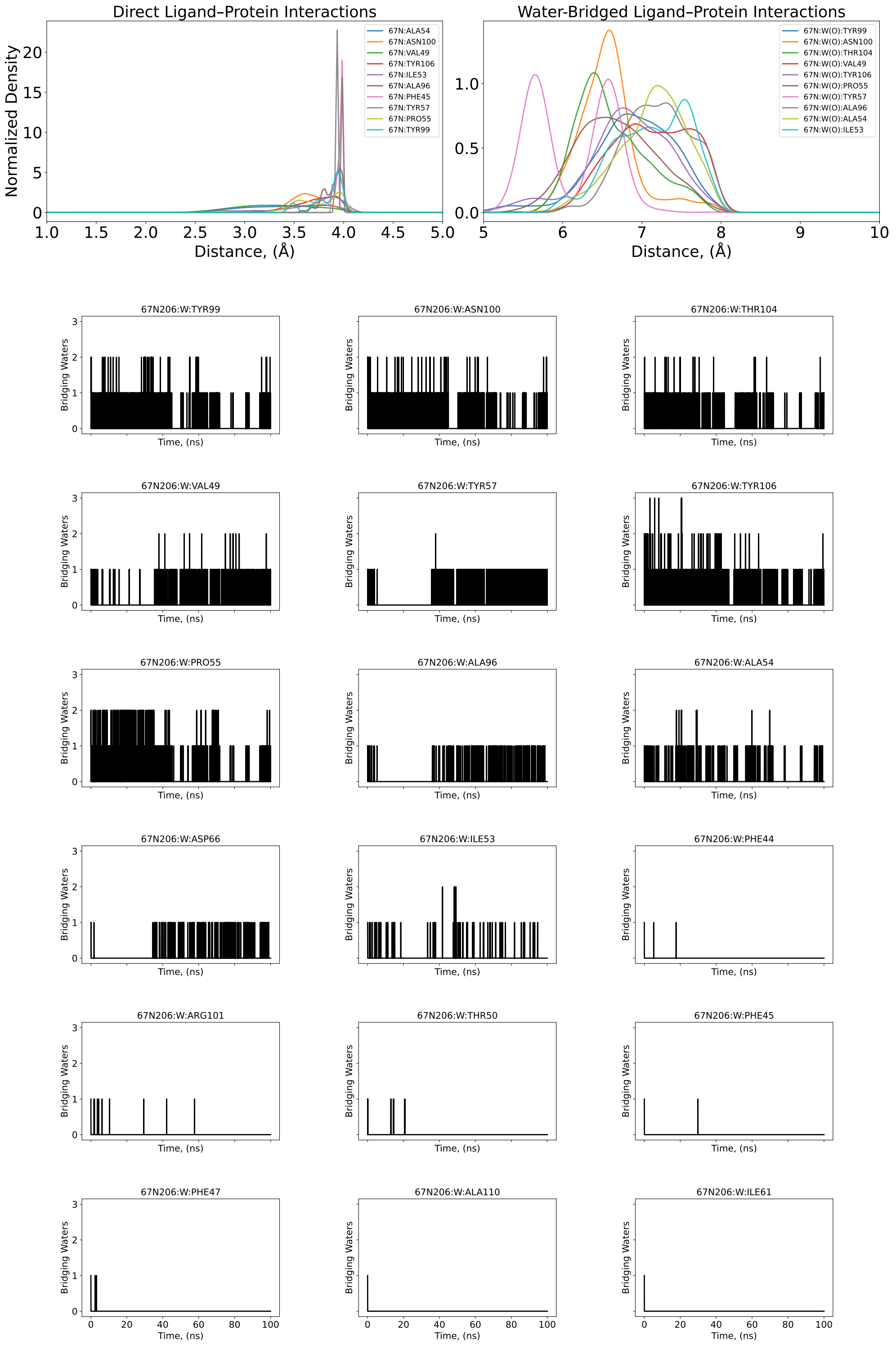


Figure S35. 4-Å–density distribution and individual coordination number of bridging oxygen water molecules (O/N-bridging) that mediate individual oxygen atoms of the protein and nitrogen atoms of the ligand along a 100-ns conventional MD for BRD9 complex (PDB: 5I40) .

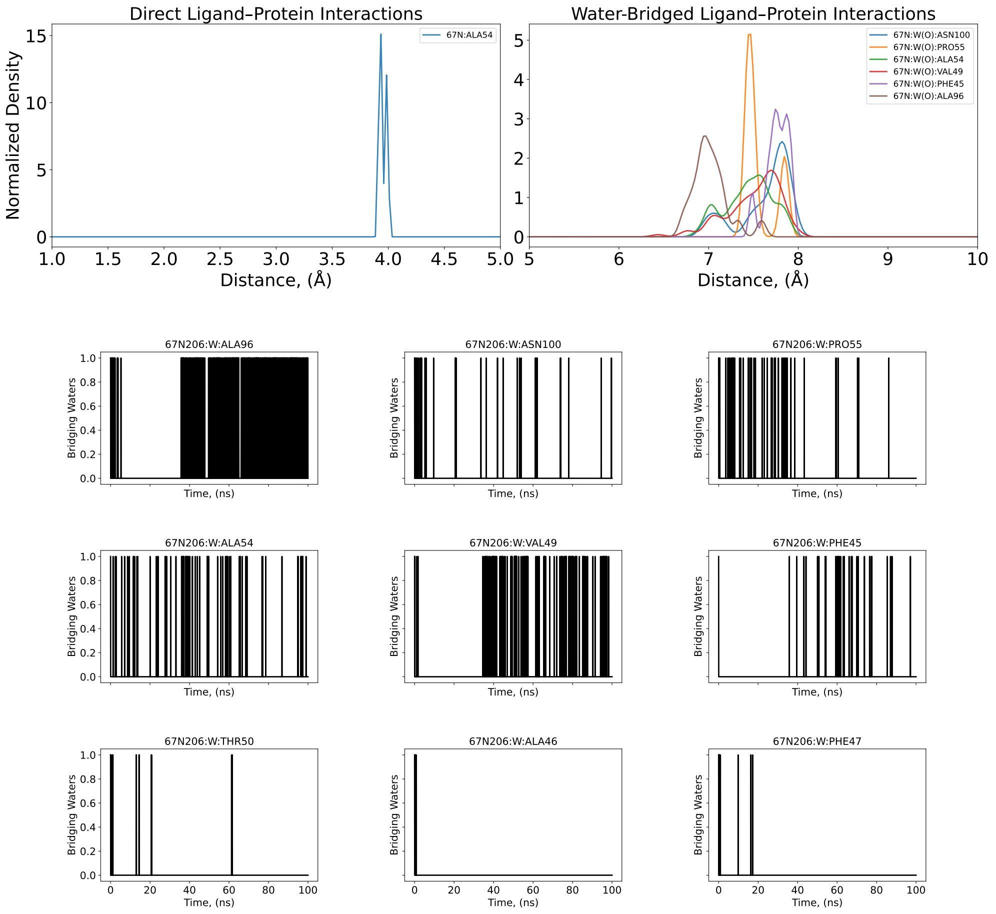


Figure S36. 4-Å–density distribution, averaged and individual coordination number of bridging oxygen water molecules (O/N-bridging) that mediate individual nitrogen atoms of the protein and nitrogen atoms of the ligand along a 100-ns conventional MD for BRD9 complex (PDB: 5I40).

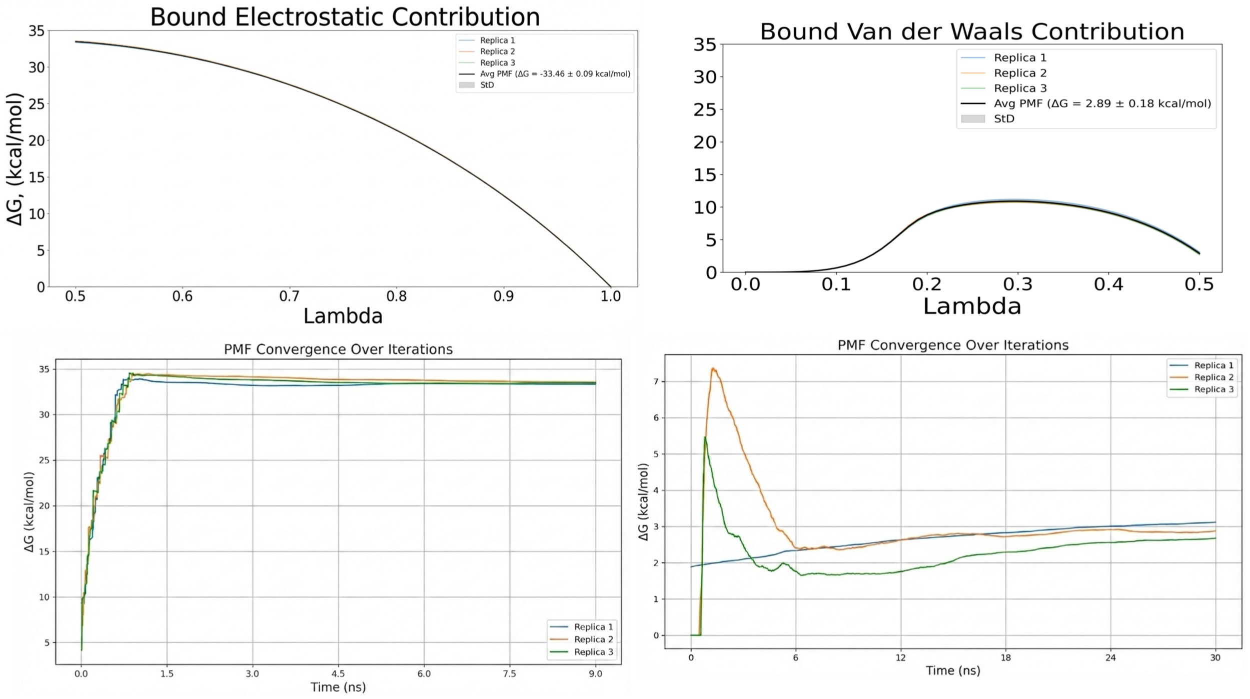


Figure S37. PMF and convergence plots of bound electrostatic and van der Waals energetic contribution for three replicas for BRD9 complex (PDB: 5I40).

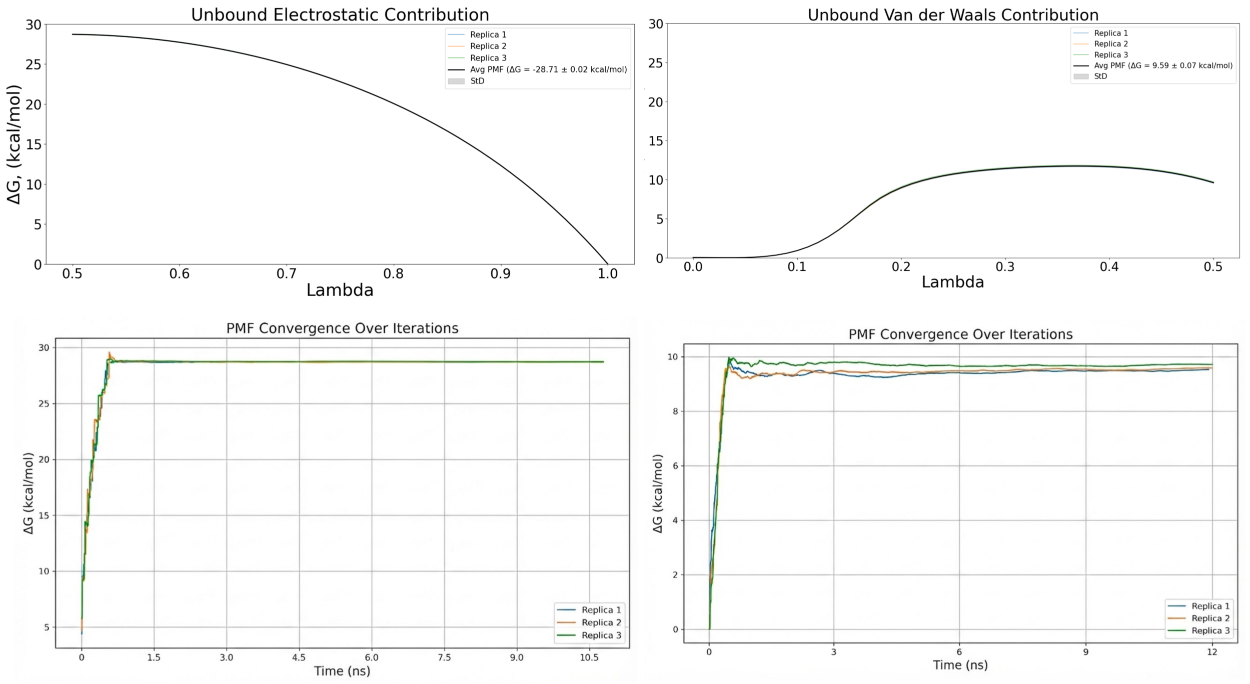


Figure S38. PMF and convergence plots of unbound electrostatic and van der Waals energetic contribution for three replicas for BRD9 complex (PDB: 5I40).

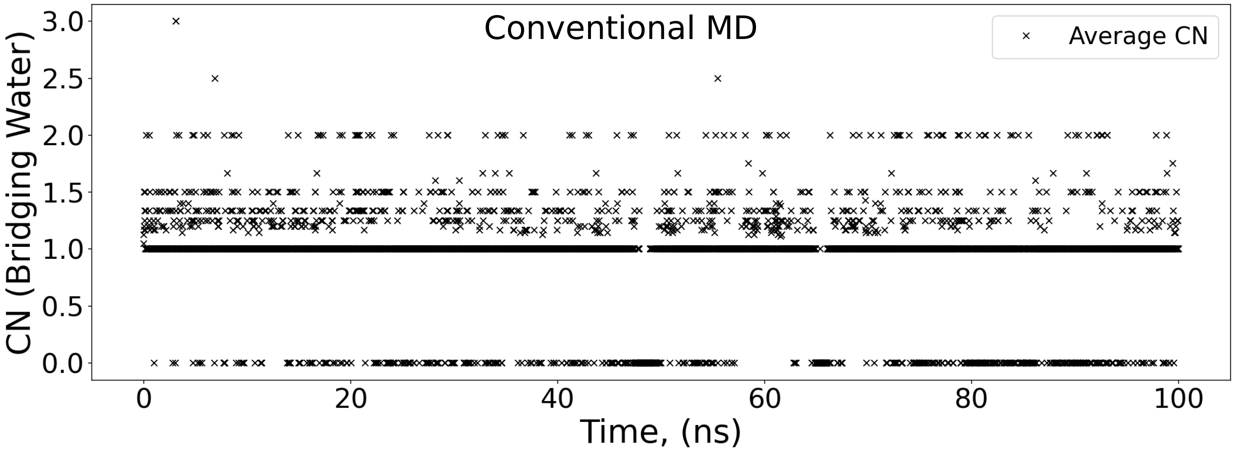


Figure S39. Time-series analysis of the cumulative bridging water coordination number (CN) over the conventional MD. Bridging events were defined by a geometric dual-occupancy criterion, requiring a water molecule to reside simultaneously within 4.0 Å of both a ligand and a protein heavy atom (oxygen or nitrogen) for BRD9 complex (PDB: 5I40).

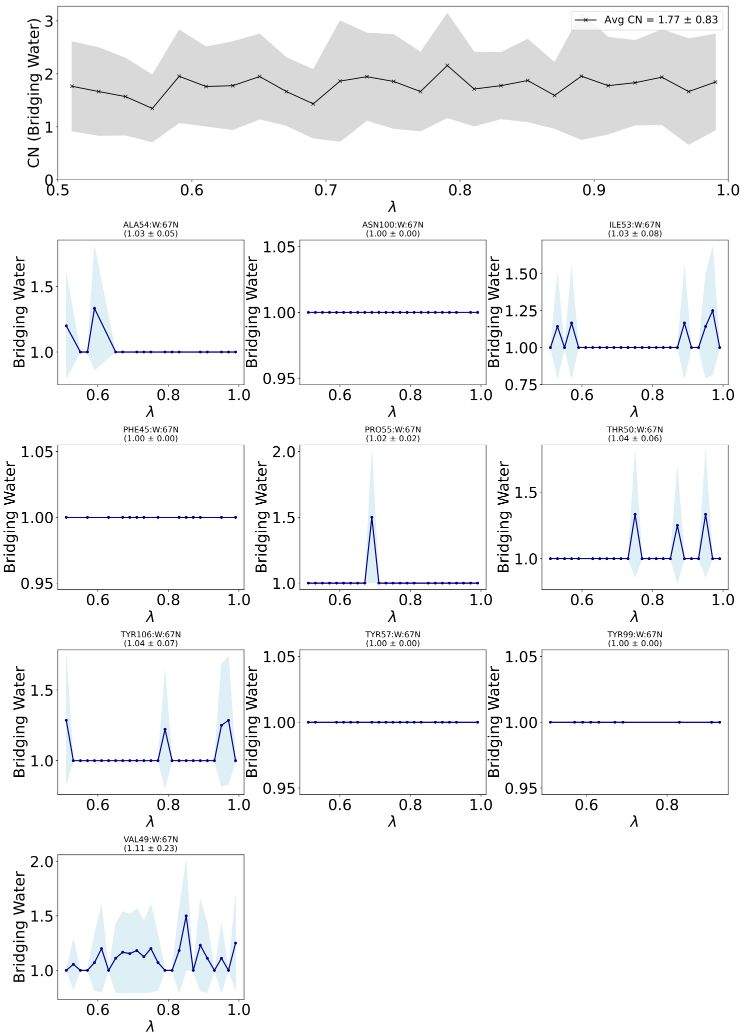


Figure S40. Water-mediated interaction profiles for the bridging motif for electrostatic bound contribution in Lambda-ABF-OPES binding affinity calculations for for BRD9 complex (PDB: 5I40). (Top Panel) Global analysis of the total hydration stoichiometry within the binding pocket. The black trace represents the conditional average count of water molecules participating in bridging interactions across all detected residue pairs. The gray shaded region indicates the standard deviation ($\pm 1\sigma$) across the ensemble, reflecting fluctuations in solvent multiplicity. (Bottom Panels) Site-specific hydration profiles for key anchoring residues. Each panel displays the average number of water molecules bridging the specific residue–ligand pair as a function of the alchemical coordinate $\lambda$. Error bands represent the fluctuation in water count during bridging events.

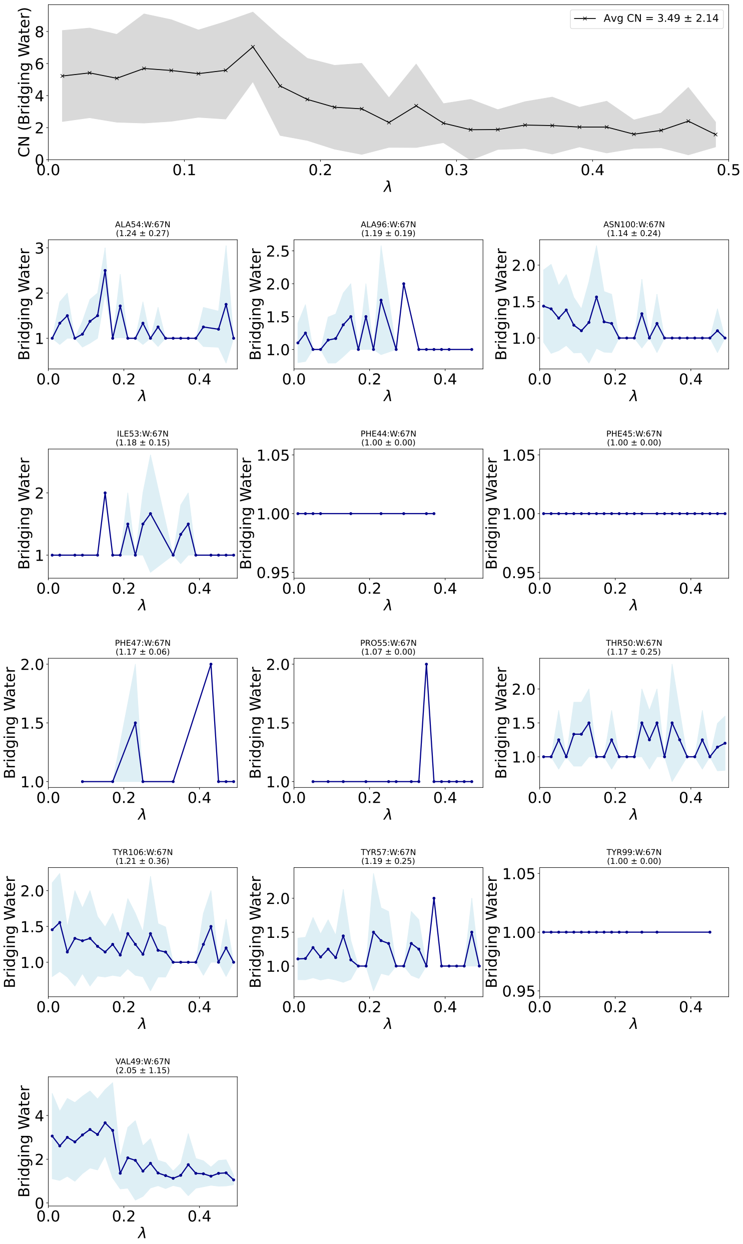


Figure S41. Water-mediated interaction profiles for the bridging motif for van der Waals bound contribution in Lambda-ABF-OPES binding affinity calculations for for BRD9 complex (PDB: 5I40). (Top Panel) Global analysis of the total hydration stoichiometry within the binding pocket. The black trace represents the conditional average count of water molecules participating in bridging interactions across all detected residue pairs. The gray shaded region indicates the standard deviation ($\pm 1\sigma$) across the ensemble, reflecting fluctuations in solvent multiplicity. (Bottom Panels) Site-specific hydration profiles for key anchoring residues. Each panel displays the average number of water molecules bridging the specific residue–ligand pair as a function of the alchemical coordinate $\lambda$. Error bands represent the fluctuation in water count during bridging events.

### HSP-90 Related Complexes

#### 2XAB

The labels O/O, N/O, O/N, and N/N denote the heavy-atom element classes used for contact classification, with the first element corresponding to the protein atom and the second to the ligand atom. Residue labels in the figures are used as compact notation. The corresponding atom-level assignments, including protein atom name and backbone versus side-chain origin, are reported in Table S8

**Table S8. Atom-level direct and water-mediated polar contacts for 2XAB. Motif labels are reported as protein-element/ligand-element. Direct contacts use a $4.0$ Å heavy-atom cutoff. Water-mediated contacts require both ligand–water and water–protein distances to be within $4.0$ Å, and ligand–protein proximity $\leq 6.0$ Å.**

| contact type | motif | contact label | atom origin | occ. fraction | mean events | direct dist Å | lig–wat dist Å | wat–prot dist Å | total path Å | unique water O |
|---|---|---|---|---|---|---|---|---|---|---|
| direct | O/O | XJG1224:O2–ALA55:O (backbone) | backbone | 0.592 | 1.00 | 3.27 | | | | |
| direct | N/O | XJG1224:O3–LYS58:NZ (side chain) | side chain | 0.421 | 1.00 | 3.50 | | | | |
| direct | O/N | XJG1224:N–GLY108:O (backbone) | backbone | 0.339 | 1.00 | 3.01 | | | | |
| direct | O/O | XJG1224:O1–GLY108:O (backbone) | backbone | 0.334 | 1.00 | 3.04 | | | | |
| direct | O/O | XJG1224:O1–VAL186:O (backbone) | backbone | 0.280 | 1.00 | 3.26 | | | | |
| direct | O/N | XJG1224:N–VAL186:O (backbone) | backbone | 0.264 | 1.00 | 3.24 | | | | |
| direct | O/O | XJG1224:O1–VAL150:O (backbone) | backbone | 0.252 | 1.00 | 3.34 | | | | |
| direct | O/N | XJG1224:N–VAL150:O (backbone) | backbone | 0.249 | 1.00 | 3.34 | | | | |
| direct | O/O | XJG1224:O1–ASN51:O (backbone) | backbone | 0.248 | 1.00 | 3.34 | | | | |
| direct | O/N | XJG1224:N–ASN51:O (backbone) | backbone | 0.241 | 1.00 | 3.29 | | | | |

*Continued on next page*

#### Table S8. Atom-level direct and water-mediated polar contacts for 2XAB (continued)

| contact type | motif | contact label | atom origin | occ. fraction | mean events | direct dist Å | lig–wat dist Å | wat–prot dist Å | total path Å | unique water O |
|---|---|---|---|---|---|---|---|---|---|---|
| direct | N/O | XJG1224:O2–GLY97:N (backbone) | backbone | 0.228 | 1.00 | 3.80 | | | | |
| direct | N/O | XJG1224:O2–LYS58:NZ (side chain) | side chain | 0.220 | 1.00 | 3.21 | | | | |
| direct | N/N | XJG1224:N–ASN51:ND2 (side chain) | side chain | 0.204 | 1.00 | 3.30 | | | | |
| direct | N/O | XJG1224:O1–ASN51:ND2 (side chain) | side chain | 0.202 | 1.00 | 3.32 | | | | |
| direct | O/O | XJG1224:O1–MET98:O (backbone) | backbone | 0.187 | 1.00 | 3.35 | | | | |
| direct | O/N | XJG1224:N–LEU107:O (backbone) | backbone | 0.185 | 1.00 | 3.36 | | | | |
| direct | O/N | XJG1224:N–MET98:O (backbone) | backbone | 0.181 | 1.00 | 3.35 | | | | |
| direct | O/O | XJG1224:O1–LEU107:O (backbone) | backbone | 0.180 | 1.00 | 3.36 | | | | |
| direct | N/N | XJG1224:N–GLY108:N (backbone) | backbone | 0.148 | 1.00 | 3.47 | | | | |
| direct | N/O | XJG1224:O1–GLY108:N (backbone) | backbone | 0.140 | 1.00 | 3.49 | | | | |
| direct | O/O | XJG1224:O1–THR184:O (backbone) | backbone | 0.125 | 1.00 | 3.53 | | | | |
| direct | O/N | XJG1224:N–THR184:O (backbone) | backbone | 0.106 | 1.00 | 3.52 | | | | |
| direct | O/O | XJG1224:O1–ASN51:OD1 (side chain) | side chain | 0.089 | 1.00 | 3.43 | | | | |
| direct | O/O | XJG1224:O3–ALA55:O (backbone) | backbone | 0.086 | 1.00 | 3.74 | | | | |
| direct | O/N | XJG1224:N–ASN51:OD1 (side chain) | side chain | 0.085 | 1.00 | 3.40 | | | | |
| direct | O/O | XJG1224:O3–GLY108:O (backbone) | backbone | 0.074 | 1.00 | 3.59 | | | | |
| direct | O/N | XJG1224:N–LEU48:O (backbone) | backbone | 0.067 | 1.00 | 3.53 | | | | |
| direct | O/O | XJG1224:O1–LEU48:O (backbone) | backbone | 0.066 | 1.00 | 3.58 | | | | |

*Continued on next page*

### Table S8. Atom-level direct and water-mediated polar contacts for 2XAB (continued)

| contact type | motif | contact label | atom origin | occ. fraction | mean events | direct dist Å | lig–wat dist Å | wat–prot dist Å | total path Å | unique water O |
|---|---|---|---|---|---|---|---|---|---|---|
| direct | O/N | XJG1224:N–THR109:O (backbone) | backbone | 0.061 | 1.00 | 3.30 | | | | |
| direct | O/O | XJG1224:O1–THR109:O (backbone) | backbone | 0.060 | 1.00 | 3.30 | | | | |
| direct | O/N | XJG1224:N–PHE138:O (backbone) | backbone | 0.056 | 1.00 | 3.76 | | | | |
| direct | O/O | XJG1224:O1–PHE138:O (backbone) | backbone | 0.051 | 1.00 | 3.75 | | | | |
| direct | N/O | XJG1224:O2–MET98:N (backbone) | backbone | 0.051 | 1.00 | 3.84 | | | | |
| direct | O/O | XJG1224:O3–LYS58:O (backbone) | backbone | 0.041 | 1.00 | 3.71 | | | | |
| direct | O/O | XJG1224:O2–GLY108:O (backbone) | backbone | 0.037 | 1.00 | 3.81 | | | | |
| direct | N/O | XJG1224:O2–ALA55:N (backbone) | backbone | 0.024 | 1.00 | 3.82 | | | | |
| direct | N/N | XJG1224:N–THR109:N (backbone) | backbone | 0.021 | 1.00 | 3.69 | | | | |
| direct | O/O | XJG1224:O3–ASP102:OD2 (side chain) | side chain | 0.020 | 1.00 | 3.77 | | | | |
| direct | N/O | XJG1224:O1–THR109:N (backbone) | backbone | 0.019 | 1.00 | 3.65 | | | | |
| water-mediated | N/O | XJG1224:O3–W(O)–LYS58:NZ (side chain) | side chain | 0.338 | 1.13 | | 3.60 | 3.24 | 6.84 | 1000 |
| water-mediated | O/N | XJG1224:N–W(O)–GLY108:O (backbone) | backbone | 0.253 | 1.28 | | 3.32 | 3.13 | 6.45 | 195 |
| water-mediated | O/O | XJG1224:O1–W(O)–GLY108:O (backbone) | backbone | 0.249 | 1.26 | | 3.35 | 3.14 | 6.48 | 194 |
| water-mediated | N/N | XJG1224:N–W(O)–THR109:N (backbone) | backbone | 0.233 | 1.28 | | 3.33 | 3.33 | 6.66 | 181 |
| water-mediated | N/O | XJG1224:O1–W(O)–THR109:N (backbone) | backbone | 0.223 | 1.26 | | 3.35 | 3.33 | 6.68 | 175 |
| water-mediated | O/O | XJG1224:O3–W(O)–LYS58:O (backbone) | backbone | 0.209 | 1.21 | | 3.60 | 2.95 | 6.56 | 597 |
| water-mediated | O/N | XJG1224:N–W(O)–ASN51:OD1 (side chain) | side chain | 0.179 | 1.23 | | 3.36 | 3.37 | 6.73 | 222 |

*Continued on next page*

#### Table S8. Atom-level direct and water-mediated polar contacts for 2XAB (continued)

| contact type | motif | contact label | atom origin | occ. fraction | mean events | direct dist Å | lig–wat dist Å | wat–prot dist Å | total path Å | unique water O |
|---|---|---|---|---|---|---|---|---|---|---|
| water-mediated | O/O | XJG1224:O1–W(O)–ASN51:OD1 (side chain) | side chain | 0.179 | 1.23 | | 3.37 | 3.36 | 6.74 | 223 |
| water-mediated | O/O | XJG1224:O2–W(O)–HSD154:O (backbone) | backbone | 0.173 | 1.01 | | 3.44 | 3.17 | 6.61 | 110 |
| water-mediated | N/O | XJG1224:O1–W(O)–ASN51:ND2 (side chain) | side chain | 0.170 | 1.17 | | 3.40 | 3.48 | 6.88 | 210 |
| water-mediated | N/N | XJG1224:N–W(O)–ASN51:ND2 (side chain) | side chain | 0.169 | 1.16 | | 3.38 | 3.49 | 6.87 | 224 |
| water-mediated | N/O | XJG1224:O2–W(O)–GLY97:N (backbone) | backbone | 0.168 | 1.00 | | 3.80 | 3.00 | 6.80 | 1 |
| water-mediated | N/O | XJG1224:O2–W(O)–ILE96:N (backbone) | backbone | 0.167 | 1.00 | | 3.80 | 3.44 | 7.24 | 1 |
| water-mediated | O/O | XJG1224:O2–W(O)–GLY95:O (backbone) | backbone | 0.166 | 1.00 | | 3.80 | 2.61 | 6.40 | 1 |
| water-mediated | O/O | XJG1224:O2–W(O)–ALA55:O (backbone) | backbone | 0.164 | 1.00 | | 3.79 | 3.18 | 6.98 | 7 |
| water-mediated | O/O | XJG1224:O3–W(O)–ASP102:OD2 (side chain) | side chain | 0.143 | 1.20 | | 3.60 | 3.15 | 6.75 | 325 |
| water-mediated | O/O | XJG1224:O3–W(O)–GLY108:O (backbone) | backbone | 0.118 | 1.15 | | 3.52 | 3.50 | 7.03 | 223 |
| water-mediated | O/O | XJG1224:O3–W(O)–ASP102:OD1 (side chain) | side chain | 0.109 | 1.23 | | 3.59 | 3.24 | 6.83 | 327 |
| water-mediated | N/O | XJG1224:O3–W(O)–GLY108:N (backbone) | backbone | 0.102 | 1.01 | | 3.60 | 3.30 | 6.90 | 124 |
| water-mediated | O/O | XJG1224:O1–W(O)–THR109:OG1 (side chain) | side chain | 0.090 | 1.35 | | 3.25 | 3.60 | 6.84 | 149 |
| water-mediated | O/O | XJG1224:O3–W(O)–ILE96:O (backbone) | backbone | 0.087 | 1.13 | | 3.62 | 3.19 | 6.82 | 340 |
| water-mediated | O/N | XJG1224:N–W(O)–THR109:OG1 (side chain) | side chain | 0.087 | 1.38 | | 3.24 | 3.60 | 6.83 | 155 |
| water-mediated | O/O | XJG1224:O1–W(O)–ASN51:O (backbone) | backbone | 0.080 | 1.10 | | 3.38 | 3.60 | 6.98 | 160 |
| water-mediated | O/O | XJG1224:O2–W(O)–ASP102:OD1 (side chain) | side chain | 0.080 | 1.11 | | 3.46 | 3.09 | 6.55 | 196 |
| water-mediated | O/N | XJG1224:N–W(O)–ASN51:O (backbone) | backbone | 0.080 | 1.08 | | 3.39 | 3.60 | 7.00 | 172 |

*Continued on next page*

#### Table S8. Atom-level direct and water-mediated polar contacts for 2XAB (continued)

| contact type | motif | contact label | atom origin | occ. fraction | mean events | direct dist Å | lig–wat dist Å | wat–prot dist Å | total path Å | unique water O |
|---|---|---|---|---|---|---|---|---|---|---|
| water-mediated | N/N | XJG1224:N–W(O)–PHE138:N (backbone) | backbone | 0.076 | 1.02 | | 3.61 | 3.18 | 6.79 | 107 |
| water-mediated | N/O | XJG1224:O1–W(O)–PHE138:N (backbone) | backbone | 0.075 | 1.01 | | 3.61 | 3.19 | 6.79 | 109 |
| water-mediated | O/O | XJG1224:O2–W(O)–ASP102:OD2 (side chain) | side chain | 0.057 | 1.12 | | 3.47 | 3.10 | 6.57 | 169 |
| water-mediated | O/O | XJG1224:O2–W(O)–GLY108:O (backbone) | backbone | 0.052 | 1.02 | | 3.60 | 3.39 | 6.99 | 118 |
| water-mediated | O/N | XJG1224:N–W(O)–THR109:O (backbone) | backbone | 0.047 | 1.19 | | 3.49 | 3.23 | 6.72 | 85 |
| water-mediated | N/O | XJG1224:O2–W(O)–LYS58:NZ (side chain) | side chain | 0.046 | 1.02 | | 3.64 | 3.09 | 6.73 | 160 |
| water-mediated | O/O | XJG1224:O3–W(O)–ASP102:O (backbone) | backbone | 0.044 | 1.09 | | 3.55 | 2.85 | 6.40 | 103 |
| water-mediated | O/O | XJG1224:O1–W(O)–THR109:O (backbone) | backbone | 0.042 | 1.25 | | 3.47 | 3.21 | 6.68 | 72 |
| water-mediated | O/O | XJG1224:O3–W(O)–HSD154:O (backbone) | backbone | 0.031 | 1.32 | | 3.61 | 3.11 | 6.72 | 131 |
| water-mediated | O/O | XJG1224:O2–W(O)–ILE96:O (backbone) | backbone | 0.030 | 1.07 | | 3.59 | 3.40 | 6.99 | 132 |
| water-mediated | O/N | XJG1224:N–W(O)–LEU107:O (backbone) | backbone | 0.029 | 1.01 | | 3.37 | 3.26 | 6.63 | 43 |
| water-mediated | N/O | XJG1224:O2–W(O)–MET98:N (backbone) | backbone | 0.029 | 1.00 | | 3.42 | 3.91 | 7.34 | 94 |
| water-mediated | O/O | XJG1224:O1–W(O)–LEU107:O (backbone) | backbone | 0.027 | 1.01 | | 3.41 | 3.31 | 6.73 | 51 |
| water-mediated | N/O | XJG1224:O3–W(O)–THR109:N (backbone) | backbone | 0.024 | 1.06 | | 3.61 | 3.54 | 7.15 | 77 |
| water-mediated | O/O | XJG1224:O2–W(O)–THR184:OG1 (side chain) | side chain | 0.023 | 1.00 | | 3.75 | 3.70 | 7.46 | 1 |
| water-mediated | N/N | XJG1224:N–W(O)–GLY108:N (backbone) | backbone | 0.021 | 1.02 | | 3.42 | 3.75 | 7.17 | 56 |
| water-mediated | O/O | XJG1224:O2–W(O)–LYS58:O (backbone) | backbone | 0.019 | 1.07 | | 3.62 | 2.83 | 6.45 | 66 |
| water-mediated | N/O | XJG1224:O1–W(O)–GLY108:N (backbone) | backbone | 0.018 | 1.00 | | 3.40 | 3.72 | 7.12 | 48 |

*Continued on next page*

#### Table S8. Atom-level direct and water-mediated polar contacts for 2XAB (continued)

| contact type | motif | contact label | atom origin | occ. fraction | mean events | direct dist Å | lig–wat dist Å | wat–prot dist Å | total path Å | unique water O |
|---|---|---|---|---|---|---|---|---|---|---|
| water-mediated | O/O | XJG1224:O1–W(O)–TYR139:O (backbone) | backbone | 0.014 | 1.05 | | 3.36 | 3.22 | 6.58 | 43 |
| water-mediated | N/O | XJG1224:O3–W(O)–THR99:N (backbone) | backbone | 0.014 | 1.00 | | 3.60 | 3.68 | 7.28 | 29 |
| water-mediated | O/N | XJG1224:N–W(O)–TYR139:O (backbone) | backbone | 0.012 | 1.03 | | 3.27 | 3.30 | 6.57 | 34 |

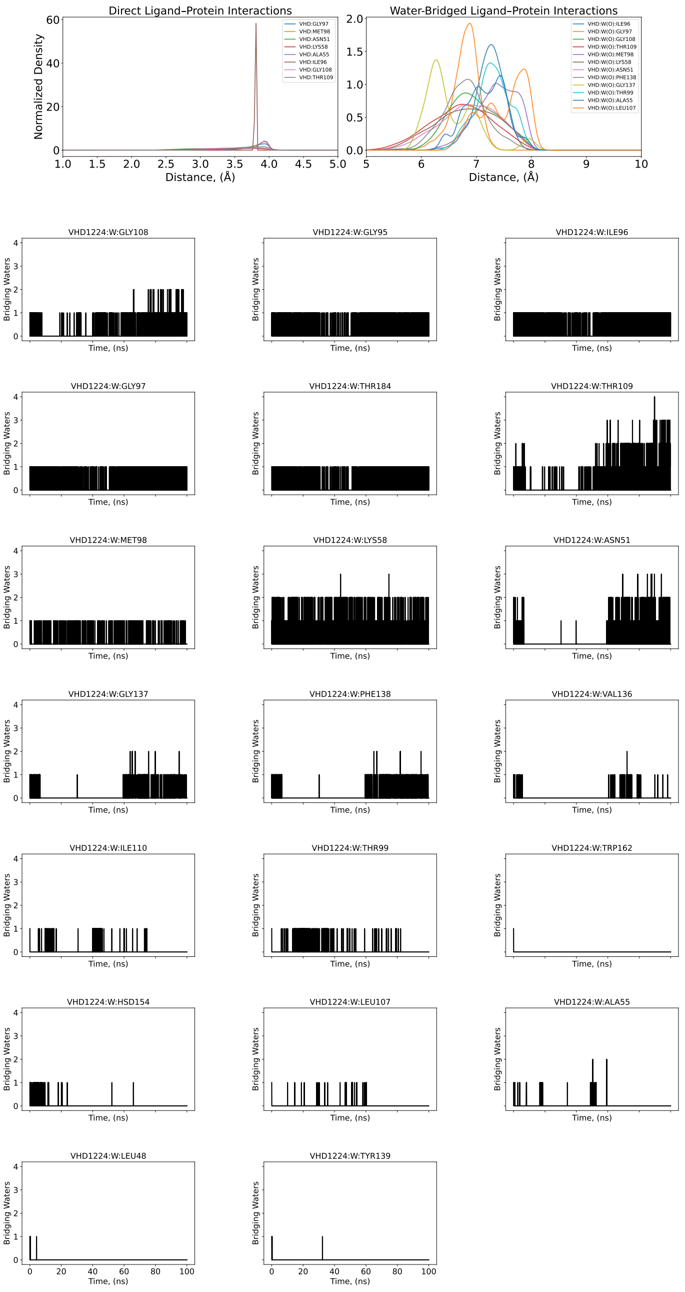


Figure S42. 4-Å–density distribution and individual coordination number of N/O-bridging water molecules that mediate individual nitrogen atoms of protein and oxygen atoms of ligand pairs along 100-ns conventional MD for the heat shock protein HSP90-$\alpha$ complex (PDB: 2XAB).

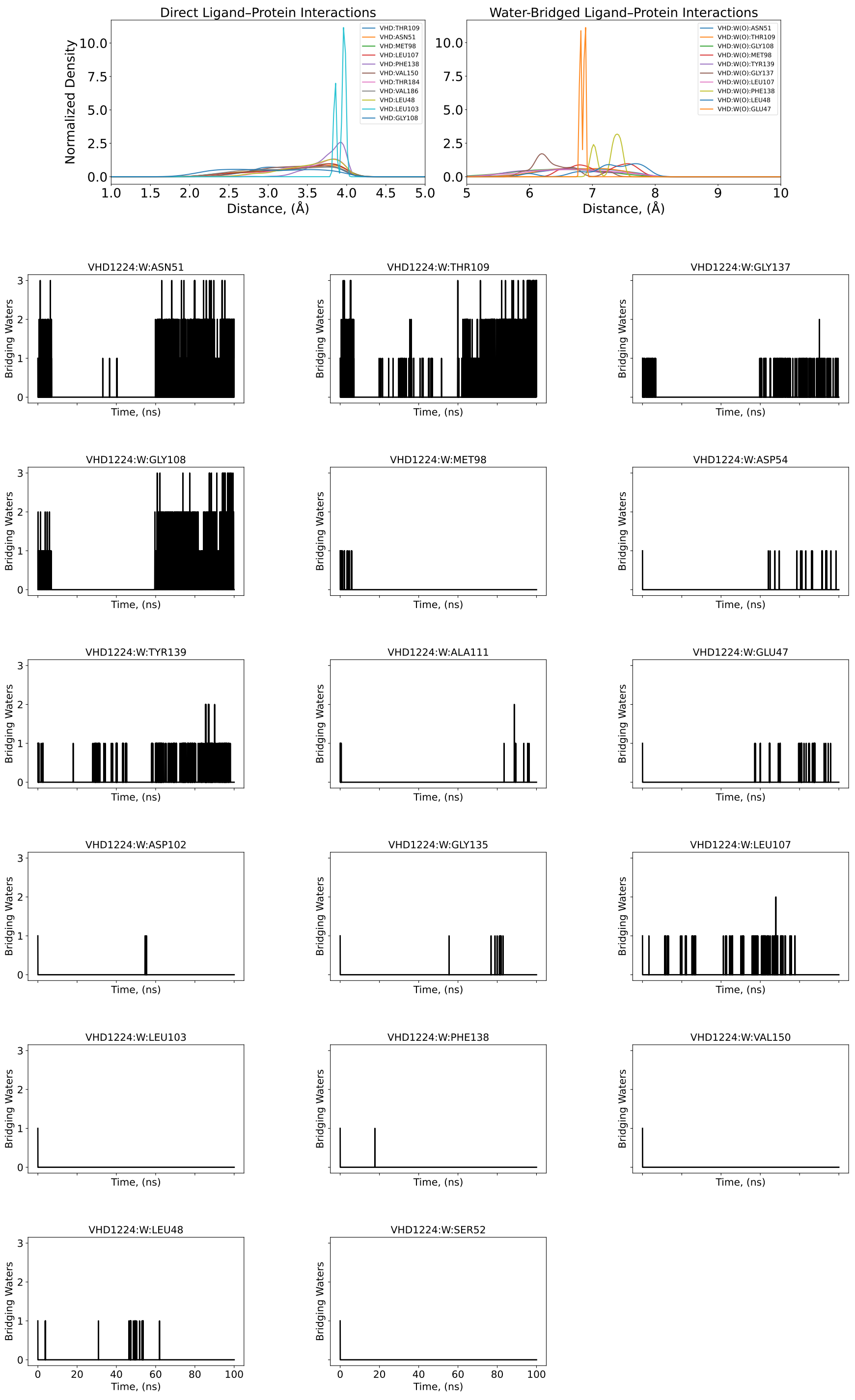

Figure S43. 4-Å–density distribution and individual coordination number of O/N-bridging water molecules that mediate individual oxygen atoms of protein and nitrogen atoms of ligand pairs along 100-ns conventional MD for the heat shock protein HSP90-$\alpha$ complex (PDB: 2XAB).

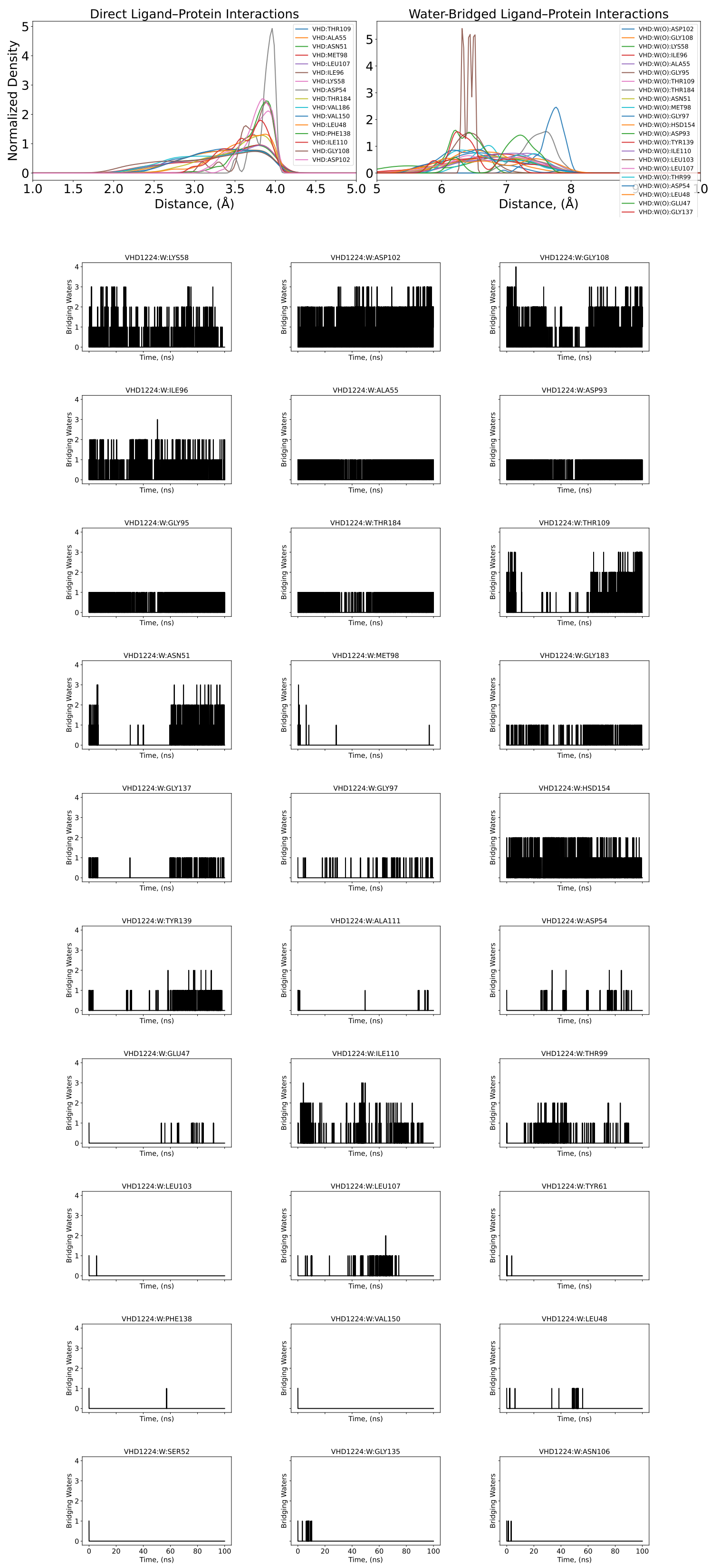


Figure S44. 4-Å–density distribution and individual coordination number of O-bridging water molecules that mediate individual oxygen protein-ligand atom pairs along 100-ns conventional MD for the heat shock protein HSP90-$\alpha$ complex (PDB: 2XAB).

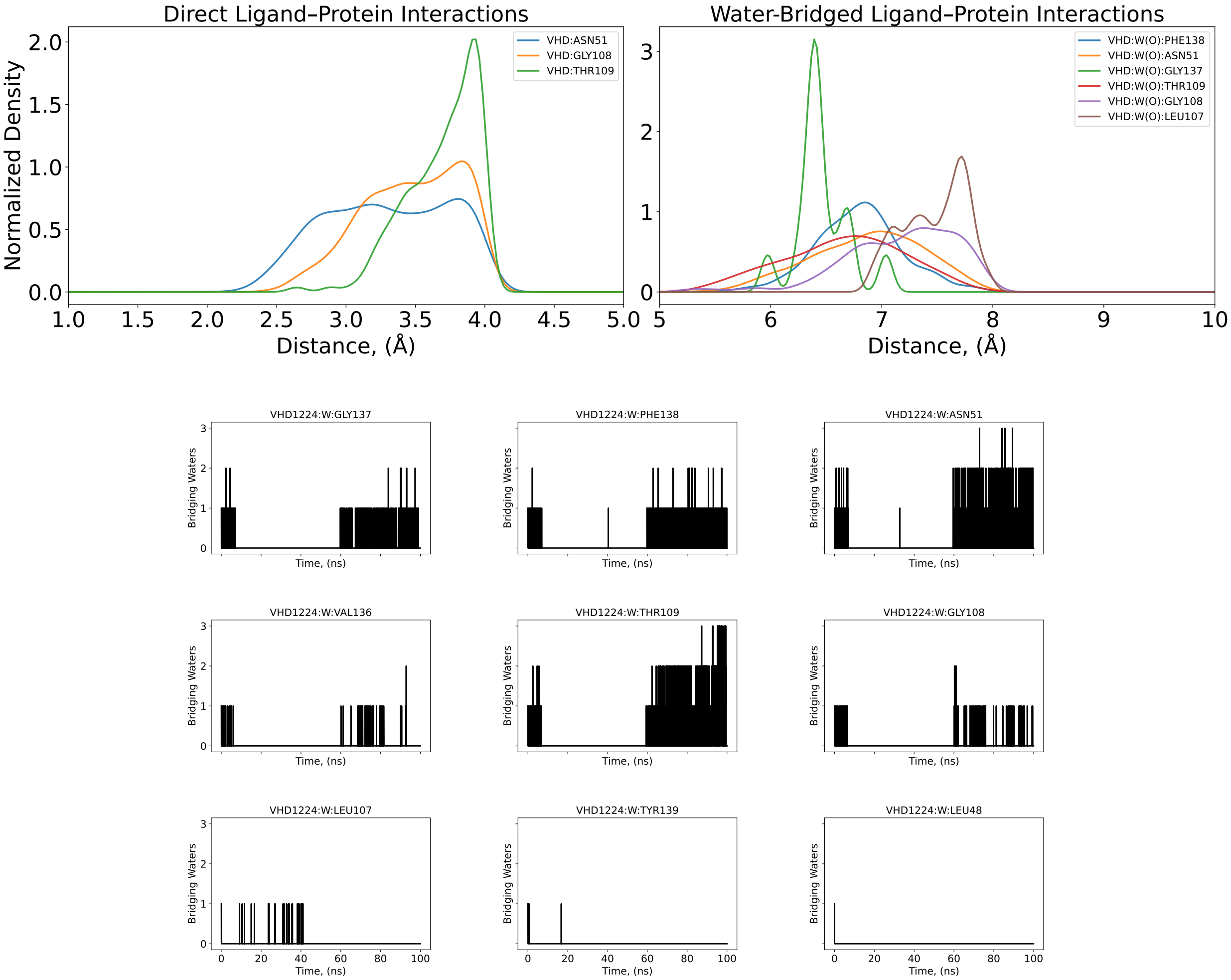


Figure S45. 4-Å–density distribution and individual coordination number of N-bridging water molecules that mediate individual nitrogen protein-ligand atom pairs along 100-ns conventional MD for the heat shock protein HSP90-$\alpha$ complex (PDB: 2XAB).

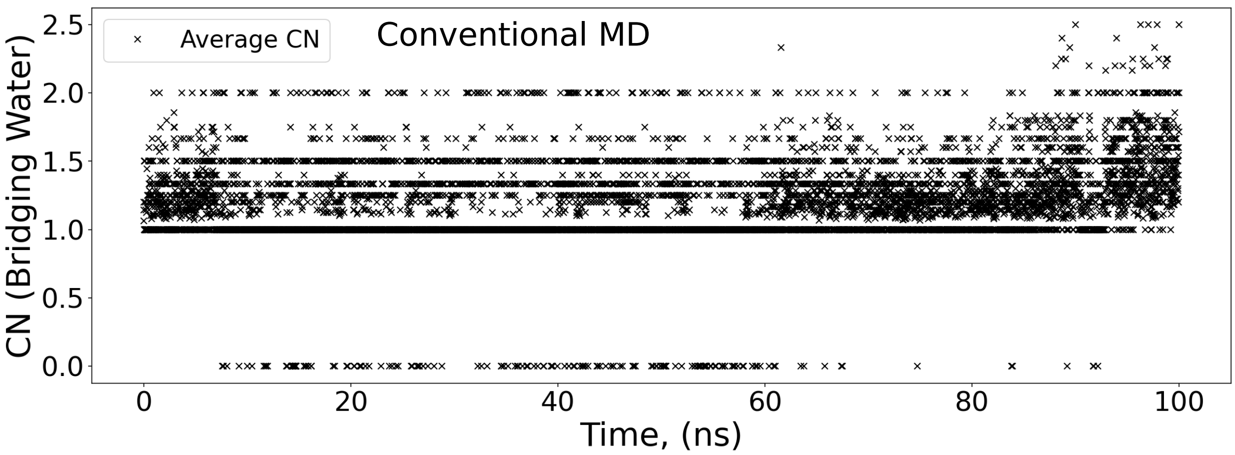


Figure S46. Time-series analysis of the cumulative bridging water coordination number (CN) over the conventional MD. Bridging events were defined by a geometric dual-occupancy criterion, requiring a water molecule to reside simultaneously within 4.0 Å of both a ligand and a protein heavy atom (oxygen or nitrogen) for the heat shock protein HSP90-$\alpha$ complex (PDB: 2XAB).

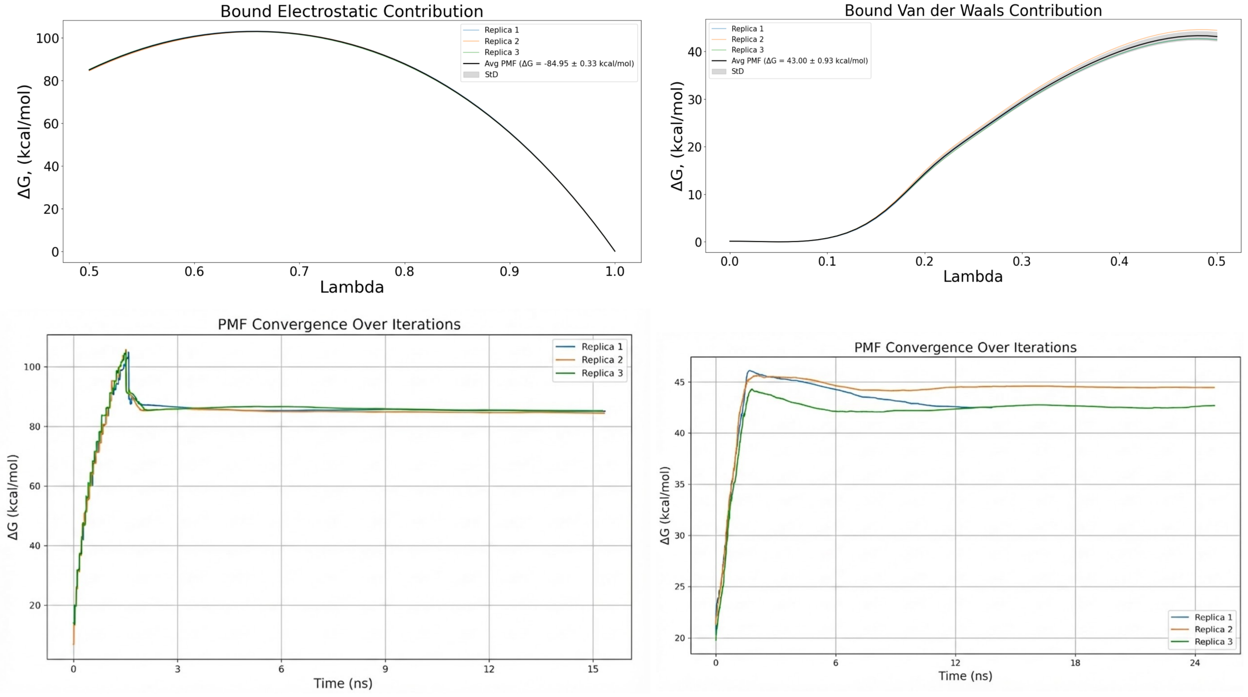


Figure S47. PMF and convergence plots of bound electrostatic and van der Waals energetic contribution for three replicas for the heat shock protein HSP90-$\alpha$ complex (PDB: 2XAB).

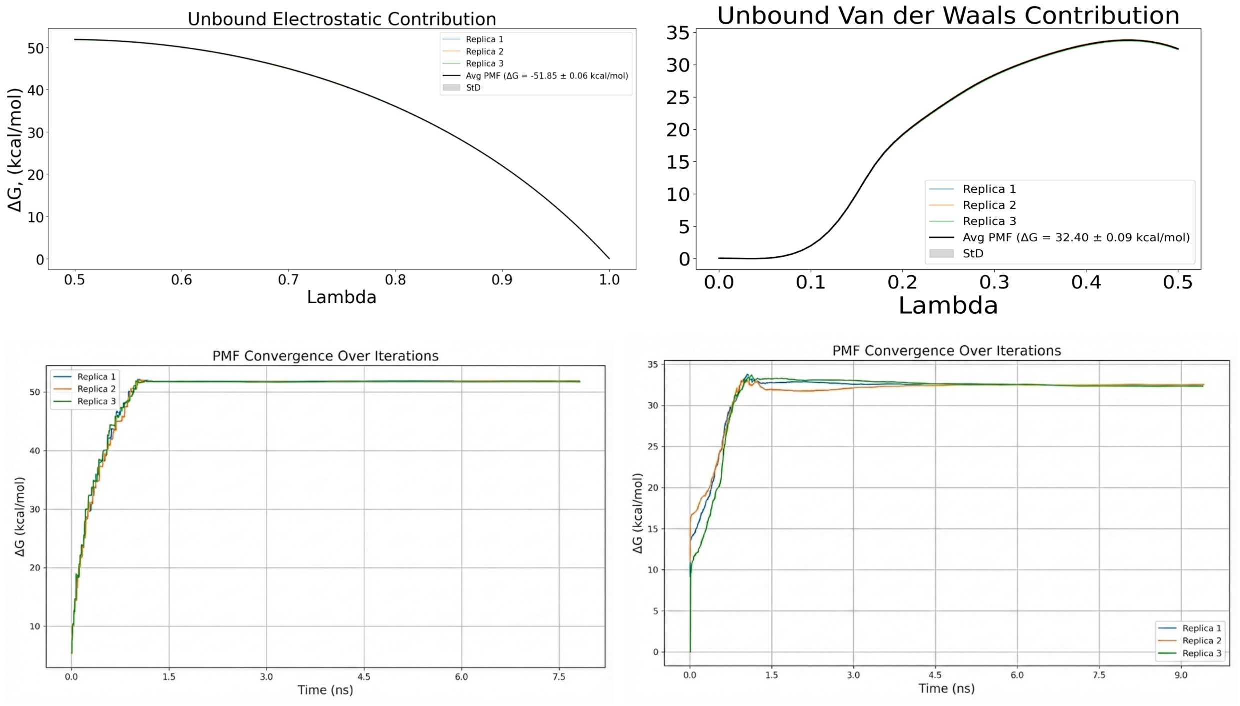


Figure S48. PMF and convergence plots of bound electrostatic and van der Waals energetic contribution for three replicas for the heat shock protein HSP90-$\alpha$ complex (PDB: 2XAB).

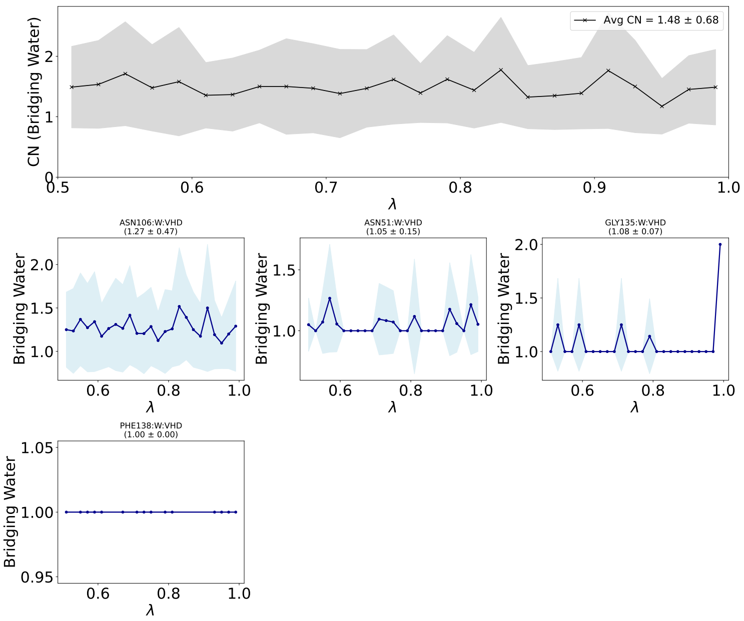


Figure S49. Water-mediated interaction profiles for the bridging motif for electrostatic bound contribution in Lambda-ABF-OPES binding affinity calculations for for the heat shock protein HSP90-$\alpha$ complex (PDB: 2XAB). (Top Panel) Global analysis of the total hydration stoichiometry within the binding pocket. The black trace represents the conditional average count of water molecules participating in bridging interactions across all detected residue pairs. The gray shaded region indicates the standard deviation ($\pm 1\sigma$) across the ensemble, reflecting fluctuations in solvent multiplicity. (Bottom Panels) Site-specific hydration profiles for key anchoring residues. Each panel displays the average number of water molecules bridging the specific residue–ligand pair as a function of the alchemical coordinate $\lambda$. Error bands represent the fluctuation in water count during bridging events.

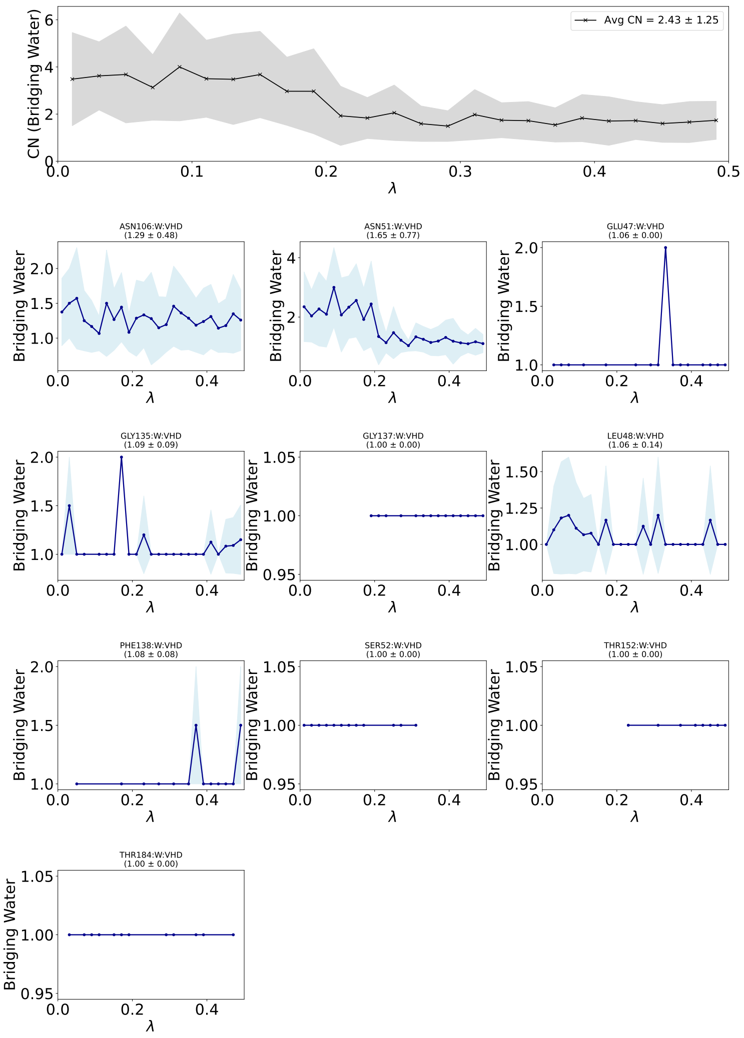


Figure S50. Water-mediated interaction profiles for the bridging motif for van der Waals bound contribution in Lambda-ABF-OPES binding affinity calculations for for the heat shock protein HSP90-$\alpha$ complex (PDB: 2XAB). (Top Panel) Global analysis of the total hydration stoichiometry within the binding pocket. The black trace represents the conditional average count of water molecules participating in bridging interactions across all detected residue pairs. The gray shaded region indicates the standard deviation ($\pm 1\sigma$) across the ensemble, reflecting fluctuations in solvent multiplicity. (Bottom Panels) Site-specific hydration profiles for key anchoring residues. Each panel displays the average number of water molecules bridging the specific residue–ligand pair as a function of the alchemical coordinate $\lambda$. Error bands represent the fluctuation in water count during bridging events.

#### 2XJG

The labels O/O, N/O, O/N, and N/N denote the heavy-atom element classes used for contact classification, with the first element corresponding to the protein atom and the second to the ligand atom. Residue labels in the figures are used as compact notation. The corresponding atom-level assignments, including protein atom name and backbone versus side-chain origin, are reported in Table S9

**Table S9. Atom-level direct and water-mediated polar contacts for 2XJG. Motif labels are reported as protein-element/ligand-element. Direct contacts use a 4.0 Å heavy-atom cutoff. Water-mediated contacts require both ligand–water and water–protein distances to be within 4.0 Å, total path length ≤ 10.0 Å, and ligand–protein proximity ≤ 6.0 Å.**

| contact type | motif | contact label | atom origin | occ. fraction | mean events | direct dist Å | lig–wat dist Å | wat–prot dist Å | total path Å | unique water O |
|---|---|---|---|---|---|---|---|---|---|---|
| direct | O/O | XJG1224:O2–ALA55:O (backbone) | backbone | 0.592 | 1.00 | 3.27 | | | | |
| direct | N/O | XJG1224:O3–LYS58:NZ (side chain) | side chain | 0.421 | 1.00 | 3.50 | | | | |
| direct | O/N | XJG1224:N–GLY108:O (backbone) | backbone | 0.339 | 1.00 | 3.01 | | | | |
| direct | O/O | XJG1224:O1–GLY108:O (backbone) | backbone | 0.334 | 1.00 | 3.04 | | | | |
| direct | O/O | XJG1224:O1–VAL186:O (backbone) | backbone | 0.280 | 1.00 | 3.26 | | | | |
| direct | O/N | XJG1224:N–VAL186:O (backbone) | backbone | 0.264 | 1.00 | 3.24 | | | | |
| direct | O/O | XJG1224:O1–VAL150:O (backbone) | backbone | 0.252 | 1.00 | 3.34 | | | | |
| direct | O/N | XJG1224:N–VAL150:O (backbone) | backbone | 0.249 | 1.00 | 3.34 | | | | |
| direct | O/O | XJG1224:O1–ASN51:O (backbone) | backbone | 0.248 | 1.00 | 3.34 | | | | |
| direct | O/N | XJG1224:N–ASN51:O (backbone) | backbone | 0.241 | 1.00 | 3.29 | | | | |
| direct | N/O | XJG1224:O2–GLY97:N (backbone) | backbone | 0.228 | 1.00 | 3.80 | | | | |

*Continued on next page*

### Table S9. Atom-level direct and water-mediated polar contacts for 2XJG (continued)

| contact type | motif | contact label | atom origin | occ. fraction | mean events | direct dist Å | lig–wat dist Å | wat–prot dist Å | total path Å | unique water O |
|---|---|---|---|---|---|---|---|---|---|---|
| direct | N/O | XJG1224:O2–LYS58:NZ (side chain) | side chain | 0.220 | 1.00 | 3.21 | | | | |
| direct | N/N | XJG1224:N–ASN51:ND2 (side chain) | side chain | 0.204 | 1.00 | 3.30 | | | | |
| direct | N/O | XJG1224:O1–ASN51:ND2 (side chain) | side chain | 0.202 | 1.00 | 3.32 | | | | |
| direct | O/O | XJG1224:O1–MET98:O (backbone) | backbone | 0.187 | 1.00 | 3.35 | | | | |
| direct | O/N | XJG1224:N–LEU107:O (backbone) | backbone | 0.185 | 1.00 | 3.36 | | | | |
| direct | O/N | XJG1224:N–MET98:O (backbone) | backbone | 0.181 | 1.00 | 3.35 | | | | |
| direct | O/O | XJG1224:O1–LEU107:O (backbone) | backbone | 0.180 | 1.00 | 3.36 | | | | |
| direct | N/N | XJG1224:N–GLY108:N (backbone) | backbone | 0.148 | 1.00 | 3.47 | | | | |
| direct | N/O | XJG1224:O1–GLY108:N (backbone) | backbone | 0.140 | 1.00 | 3.49 | | | | |
| direct | O/O | XJG1224:O1–THR184:O (backbone) | backbone | 0.125 | 1.00 | 3.53 | | | | |
| direct | O/N | XJG1224:N–THR184:O (backbone) | backbone | 0.106 | 1.00 | 3.52 | | | | |
| direct | O/O | XJG1224:O1–ASN51:OD1 (side chain) | side chain | 0.089 | 1.00 | 3.43 | | | | |
| direct | O/O | XJG1224:O3–ALA55:O (backbone) | backbone | 0.086 | 1.00 | 3.74 | | | | |
| direct | O/N | XJG1224:N–ASN51:OD1 (side chain) | side chain | 0.085 | 1.00 | 3.40 | | | | |
| direct | O/O | XJG1224:O3–GLY108:O (backbone) | backbone | 0.074 | 1.00 | 3.59 | | | | |
| direct | O/N | XJG1224:N–LEU48:O (backbone) | backbone | 0.067 | 1.00 | 3.53 | | | | |
| direct | O/O | XJG1224:O1–LEU48:O (backbone) | backbone | 0.066 | 1.00 | 3.58 | | | | |
| direct | O/N | XJG1224:N–THR109:O (backbone) | backbone | 0.061 | 1.00 | 3.30 | | | | |

*Continued on next page*

### Table S9. Atom-level direct and water-mediated polar contacts for 2XJG (continued)

| contact type | motif | contact label | atom origin | occ. fraction | mean events | direct dist Å | lig–wat dist Å | wat–prot dist Å | total path Å | unique water O |
|---|---|---|---|---|---|---|---|---|---|---|
| direct | O/O | XJG1224:O1–THR109:O (backbone) | backbone | 0.060 | 1.00 | 3.30 | | | | |
| direct | O/N | XJG1224:N–PHE138:O (backbone) | backbone | 0.056 | 1.00 | 3.76 | | | | |
| direct | O/O | XJG1224:O1–PHE138:O (backbone) | backbone | 0.051 | 1.00 | 3.75 | | | | |
| direct | N/O | XJG1224:O2–MET98:N (backbone) | backbone | 0.051 | 1.00 | 3.84 | | | | |
| direct | O/O | XJG1224:O3–LYS58:O (backbone) | backbone | 0.041 | 1.00 | 3.71 | | | | |
| direct | O/O | XJG1224:O2–GLY108:O (backbone) | backbone | 0.037 | 1.00 | 3.81 | | | | |
| direct | N/O | XJG1224:O2–ALA55:N (backbone) | backbone | 0.024 | 1.00 | 3.82 | | | | |
| direct | N/N | XJG1224:N–THR109:N (backbone) | backbone | 0.021 | 1.00 | 3.69 | | | | |
| direct | O/O | XJG1224:O3–ASP102:OD2 (side chain) | side chain | 0.020 | 1.00 | 3.77 | | | | |
| direct | N/O | XJG1224:O1–THR109:N (backbone) | backbone | 0.019 | 1.00 | 3.65 | | | | |
| water-mediated | N/O | XJG1224:O3–W(O)–LYS58:NZ (side chain) | side chain | 0.338 | 1.13 | | 3.60 | 3.24 | 6.84 | 1000 |
| water-mediated | O/N | XJG1224:N–W(O)–GLY108:O (backbone) | backbone | 0.253 | 1.28 | | 3.32 | 3.13 | 6.45 | 195 |
| water-mediated | O/O | XJG1224:O1–W(O)–GLY108:O (backbone) | backbone | 0.249 | 1.26 | | 3.35 | 3.14 | 6.48 | 194 |
| water-mediated | N/N | XJG1224:N–W(O)–THR109:N (backbone) | backbone | 0.233 | 1.28 | | 3.33 | 3.33 | 6.66 | 181 |
| water-mediated | N/O | XJG1224:O1–W(O)–THR109:N (backbone) | backbone | 0.223 | 1.26 | | 3.35 | 3.33 | 6.68 | 175 |
| water-mediated | O/O | XJG1224:O3–W(O)–LYS58:O (backbone) | backbone | 0.209 | 1.21 | | 3.60 | 2.95 | 6.56 | 597 |
| water-mediated | O/N | XJG1224:N–W(O)–ASN51:OD1 (side chain) | side chain | 0.179 | 1.23 | | 3.36 | 3.37 | 6.73 | 222 |
| water-mediated | O/O | XJG1224:O1–W(O)–ASN51:OD1 (side chain) | side chain | 0.179 | 1.23 | | 3.37 | 3.36 | 6.74 | 223 |

*Continued on next page*

### Table S9. Atom-level direct and water-mediated polar contacts for 2XJG (continued)

| contact type | motif | contact label | atom origin | occ. fraction | mean events | direct dist Å | lig–wat dist Å | wat–prot dist Å | total path Å | unique water O |
|---|---|---|---|---|---|---|---|---|---|---|
| water-mediated | O/O | XJG1224:O2–W(O)–HSD154:O (backbone) | backbone | 0.173 | 1.01 | | 3.44 | 3.17 | 6.61 | 110 |
| water-mediated | N/O | XJG1224:O1–W(O)–ASN51:ND2 (side chain) | side chain | 0.170 | 1.17 | | 3.40 | 3.48 | 6.88 | 210 |
| water-mediated | N/N | XJG1224:N–W(O)–ASN51:ND2 (side chain) | side chain | 0.169 | 1.16 | | 3.38 | 3.49 | 6.87 | 224 |
| water-mediated | N/O | XJG1224:O2–W(O)–GLY97:N (backbone) | backbone | 0.168 | 1.00 | | 3.80 | 3.00 | 6.80 | 1 |
| water-mediated | N/O | XJG1224:O2–W(O)–ILE96:N (backbone) | backbone | 0.167 | 1.00 | | 3.80 | 3.44 | 7.24 | 1 |
| water-mediated | O/O | XJG1224:O2–W(O)–GLY95:O (backbone) | backbone | 0.166 | 1.00 | | 3.80 | 2.61 | 6.40 | 1 |
| water-mediated | O/O | XJG1224:O2–W(O)–ALA55:O (backbone) | backbone | 0.164 | 1.00 | | 3.79 | 3.18 | 6.98 | 7 |
| water-mediated | O/O | XJG1224:O3–W(O)–ASP102:OD2 (side chain) | side chain | 0.143 | 1.20 | | 3.60 | 3.15 | 6.75 | 325 |
| water-mediated | O/O | XJG1224:O3–W(O)–GLY108:O (backbone) | backbone | 0.118 | 1.15 | | 3.52 | 3.50 | 7.03 | 223 |
| water-mediated | O/O | XJG1224:O3–W(O)–ASP102:OD1 (side chain) | side chain | 0.109 | 1.23 | | 3.59 | 3.24 | 6.83 | 327 |
| water-mediated | N/O | XJG1224:O3–W(O)–GLY108:N (backbone) | backbone | 0.102 | 1.01 | | 3.60 | 3.30 | 6.90 | 124 |
| water-mediated | O/O | XJG1224:O1–W(O)–THR109:OG1 (side chain) | side chain | 0.090 | 1.35 | | 3.25 | 3.60 | 6.84 | 149 |
| water-mediated | O/O | XJG1224:O3–W(O)–ILE96:O (backbone) | backbone | 0.087 | 1.13 | | 3.62 | 3.19 | 6.82 | 340 |
| water-mediated | O/N | XJG1224:N–W(O)–THR109:OG1 (side chain) | side chain | 0.087 | 1.38 | | 3.24 | 3.60 | 6.83 | 155 |
| water-mediated | O/O | XJG1224:O1–W(O)–ASN51:O (backbone) | backbone | 0.080 | 1.10 | | 3.38 | 3.60 | 6.98 | 160 |
| water-mediated | O/O | XJG1224:O2–W(O)–ASP102:OD1 (side chain) | side chain | 0.080 | 1.11 | | 3.46 | 3.09 | 6.55 | 196 |
| water-mediated | O/N | XJG1224:N–W(O)–ASN51:O (backbone) | backbone | 0.080 | 1.08 | | 3.39 | 3.60 | 7.00 | 172 |
| water-mediated | N/N | XJG1224:N–W(O)–PHE138:N (backbone) | backbone | 0.076 | 1.02 | | 3.61 | 3.18 | 6.79 | 107 |

*Continued on next page*

### Table S9. Atom-level direct and water-mediated polar contacts for 2XJG (continued)

| contact type | motif | contact label | atom origin | occ. fraction | mean events | direct dist Å | lig–wat dist Å | wat–prot dist Å | total path Å | unique water O |
|---|---|---|---|---|---|---|---|---|---|---|
| water-mediated | N/O | XJG1224:O1–W(O)–PHE138:N (backbone) | backbone | 0.075 | 1.01 | | 3.61 | 3.19 | 6.79 | 109 |
| water-mediated | O/O | XJG1224:O2–W(O)–ASP102:OD2 (side chain) | side chain | 0.057 | 1.12 | | 3.47 | 3.10 | 6.57 | 169 |
| water-mediated | O/O | XJG1224:O2–W(O)–GLY108:O (backbone) | backbone | 0.052 | 1.02 | | 3.60 | 3.39 | 6.99 | 118 |
| water-mediated | O/N | XJG1224:N–W(O)–THR109:O (backbone) | backbone | 0.047 | 1.19 | | 3.49 | 3.23 | 6.72 | 85 |
| water-mediated | N/O | XJG1224:O2–W(O)–LYS58:NZ (side chain) | side chain | 0.046 | 1.02 | | 3.64 | 3.09 | 6.73 | 160 |
| water-mediated | O/O | XJG1224:O3–W(O)–ASP102:O (backbone) | backbone | 0.044 | 1.09 | | 3.55 | 2.85 | 6.40 | 103 |
| water-mediated | O/O | XJG1224:O1–W(O)–THR109:O (backbone) | backbone | 0.042 | 1.25 | | 3.47 | 3.21 | 6.68 | 72 |
| water-mediated | O/O | XJG1224:O3–W(O)–HSD154:O (backbone) | backbone | 0.031 | 1.32 | | 3.61 | 3.11 | 6.72 | 131 |
| water-mediated | O/O | XJG1224:O2–W(O)–ILE96:O (backbone) | backbone | 0.030 | 1.07 | | 3.59 | 3.40 | 6.99 | 132 |
| water-mediated | O/N | XJG1224:N–W(O)–LEU107:O (backbone) | backbone | 0.029 | 1.01 | | 3.37 | 3.26 | 6.63 | 43 |
| water-mediated | N/O | XJG1224:O2–W(O)–MET98:N (backbone) | backbone | 0.029 | 1.00 | | 3.42 | 3.91 | 7.34 | 94 |
| water-mediated | O/O | XJG1224:O1–W(O)–LEU107:O (backbone) | backbone | 0.027 | 1.01 | | 3.41 | 3.31 | 6.73 | 51 |
| water-mediated | N/O | XJG1224:O3–W(O)–THR109:N (backbone) | backbone | 0.024 | 1.06 | | 3.61 | 3.54 | 7.15 | 77 |
| water-mediated | O/O | XJG1224:O2–W(O)–THR184:OG1 (side chain) | side chain | 0.023 | 1.00 | | 3.75 | 3.70 | 7.46 | 1 |
| water-mediated | N/N | XJG1224:N–W(O)–GLY108:N (backbone) | backbone | 0.021 | 1.02 | | 3.42 | 3.75 | 7.17 | 56 |
| water-mediated | O/O | XJG1224:O2–W(O)–LYS58:O (backbone) | backbone | 0.019 | 1.07 | | 3.62 | 2.83 | 6.45 | 66 |
| water-mediated | N/O | XJG1224:O1–W(O)–GLY108:N (backbone) | backbone | 0.018 | 1.00 | | 3.40 | 3.72 | 7.12 | 48 |
| water-mediated | O/O | XJG1224:O1–W(O)–TYR139:O (backbone) | backbone | 0.014 | 1.05 | | 3.36 | 3.22 | 6.58 | 43 |

*Continued on next page*

#### Table S9. Atom-level direct and water-mediated polar contacts for 2XJG (continued)

| contact type | motif | contact label | atom origin | occ. fraction | mean events | direct dist Å | lig–wat dist Å | wat–prot dist Å | total path Å | unique water O |
|---|---|---|---|---|---|---|---|---|---|---|
| water-mediated | N/O | XJG1224:O3–W(O)–THR99:N (backbone) | backbone | 0.014 | 1.00 | | 3.60 | 3.68 | 7.28 | 29 |
| water-mediated | O/N | XJG1224:N–W(O)–TYR139:O (backbone) | backbone | 0.012 | 1.03 | | 3.27 | 3.30 | 6.57 | 34 |

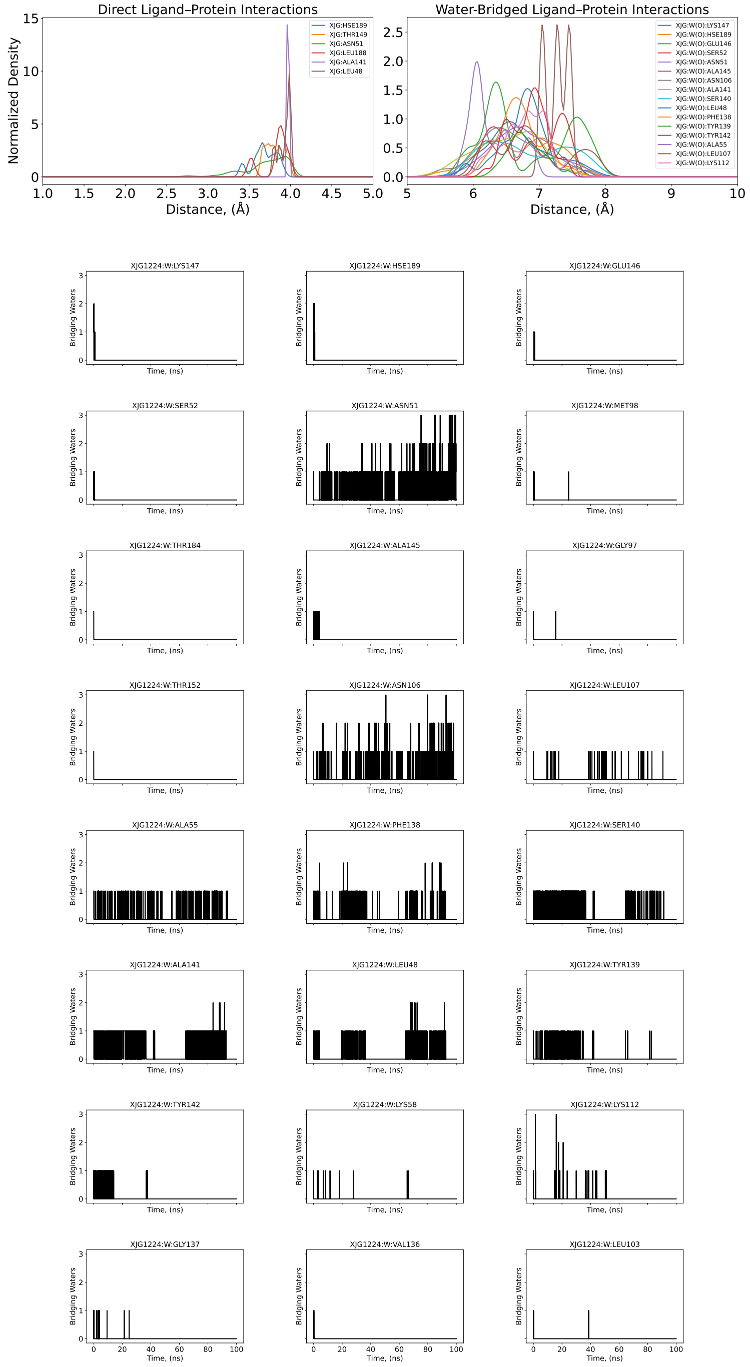


Figure S51. 4-Å–density distribution and individual coordination number of N/O-bridging water molecules that mediate individual nitrogen atoms of protein and oxygen atoms of ligand pairs along 100-ns conventional MD for the heat shock protein HSP90-$\alpha$ complex (PDB: 2XJG).

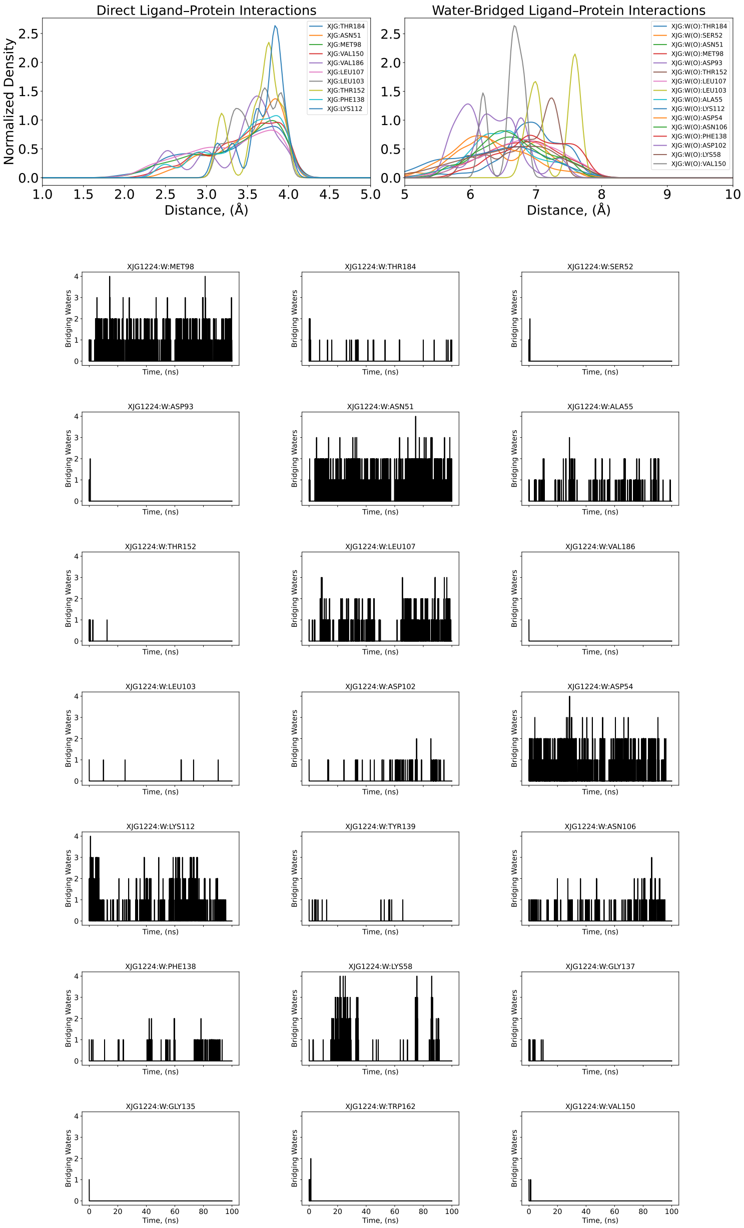

Figure S52. 4-Å–density distribution and individual coordination number of O/N-bridging water molecules that mediate individual oxygen atoms of protein and nitrogen atoms of ligand pairs along 100-ns conventional MD for the heat shock protein HSP90-$\alpha$ complex (PDB: 2XJG).

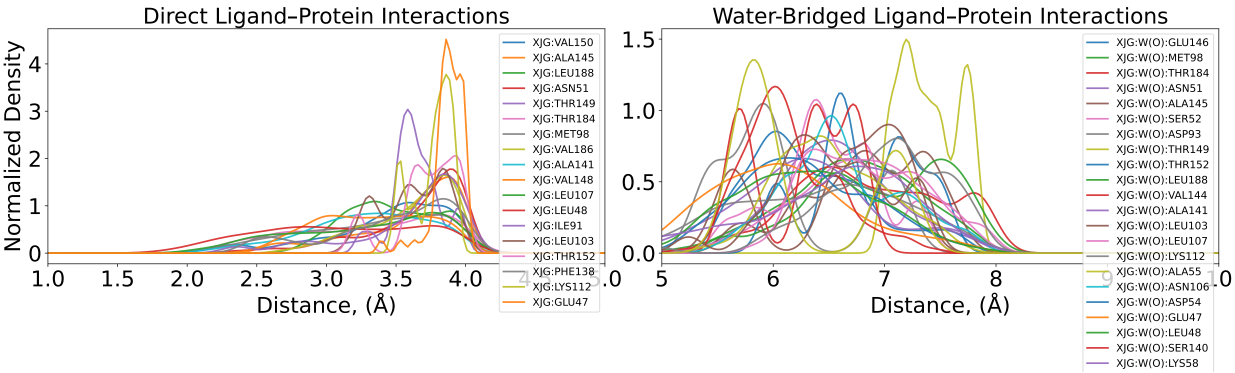
Direct Ligand–Protein Interactions
Normalized Density
Distance, (Å)
XJG:VAL150
XJG:ALA145
XJG:LEU188
XJG:ASN51
XJG:THR149
XJG:THR184
XJG:MET98
XJG:VAL186
XJG:ALA141
XJG:VAL148
XJG:LEU107
XJG:LEU48
XJG:ILE91
XJG:LEU103
XJG:THR152
XJG:PHE138
XJG:LYS112
XJG:GLU47
Water-Bridged Ligand–Protein Interactions
Distance, (Å)
XJG:W(O):GLU146
XJG:W(O):MET98
XJG:W(O):THR184
XJG:W(O):ASN51
XJG:W(O):ALA145
XJG:W(O):SER52
XJG:W(O):ASP93
XJG:W(O):THR149
XJG:W(O):THR152
XJG:W(O):LEU188
XJG:W(O):VAL144
XJG:W(O):ALA141
XJG:W(O):LEU103
XJG:W(O):LEU107
XJG:W(O):LYS112
XJG:W(O):ALA55
XJG:W(O):ASN106
XJG:W(O):ASP54
XJG:W(O):GLU47
XJG:W(O):LEU48
XJG:W(O):SER140
XJG:W(O):LYS58

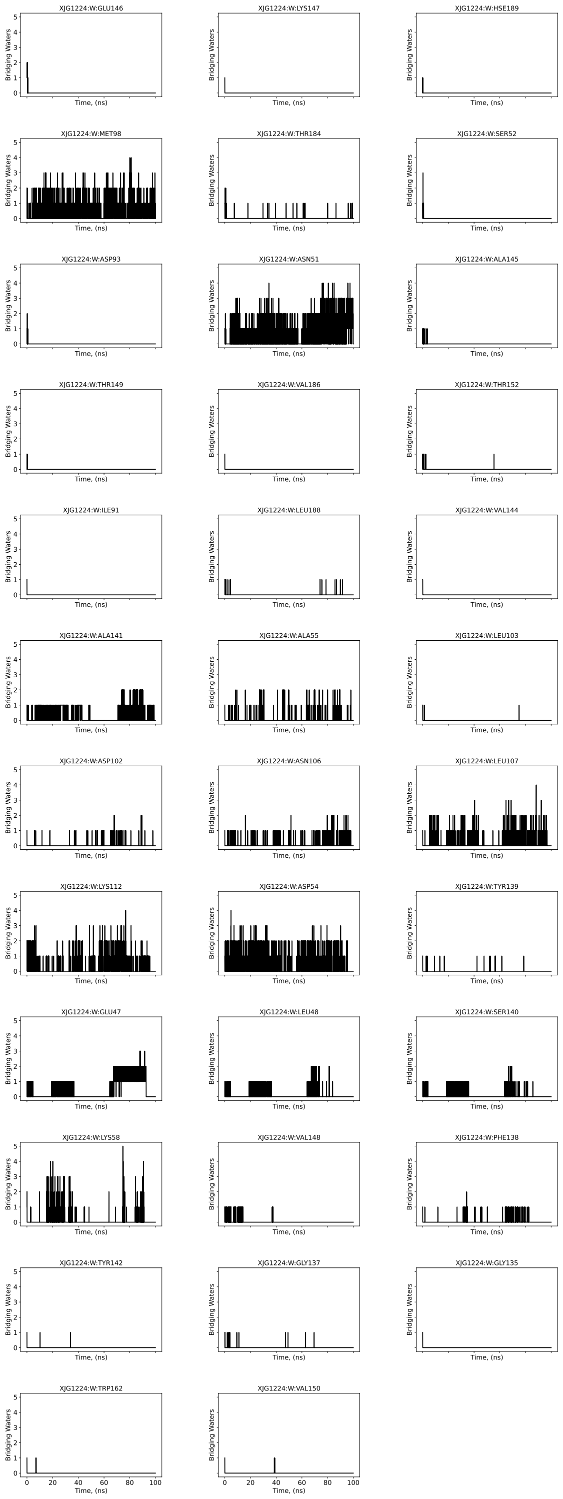


Figure S53. 4-Å–density distribution and individual coordination number of O-bridging water molecules that mediate individual oxygen protein-ligand atom pairs along 100-ns conventional MD for the heat shock protein HSP90-$\alpha$ complex (PDB: 2XJG).

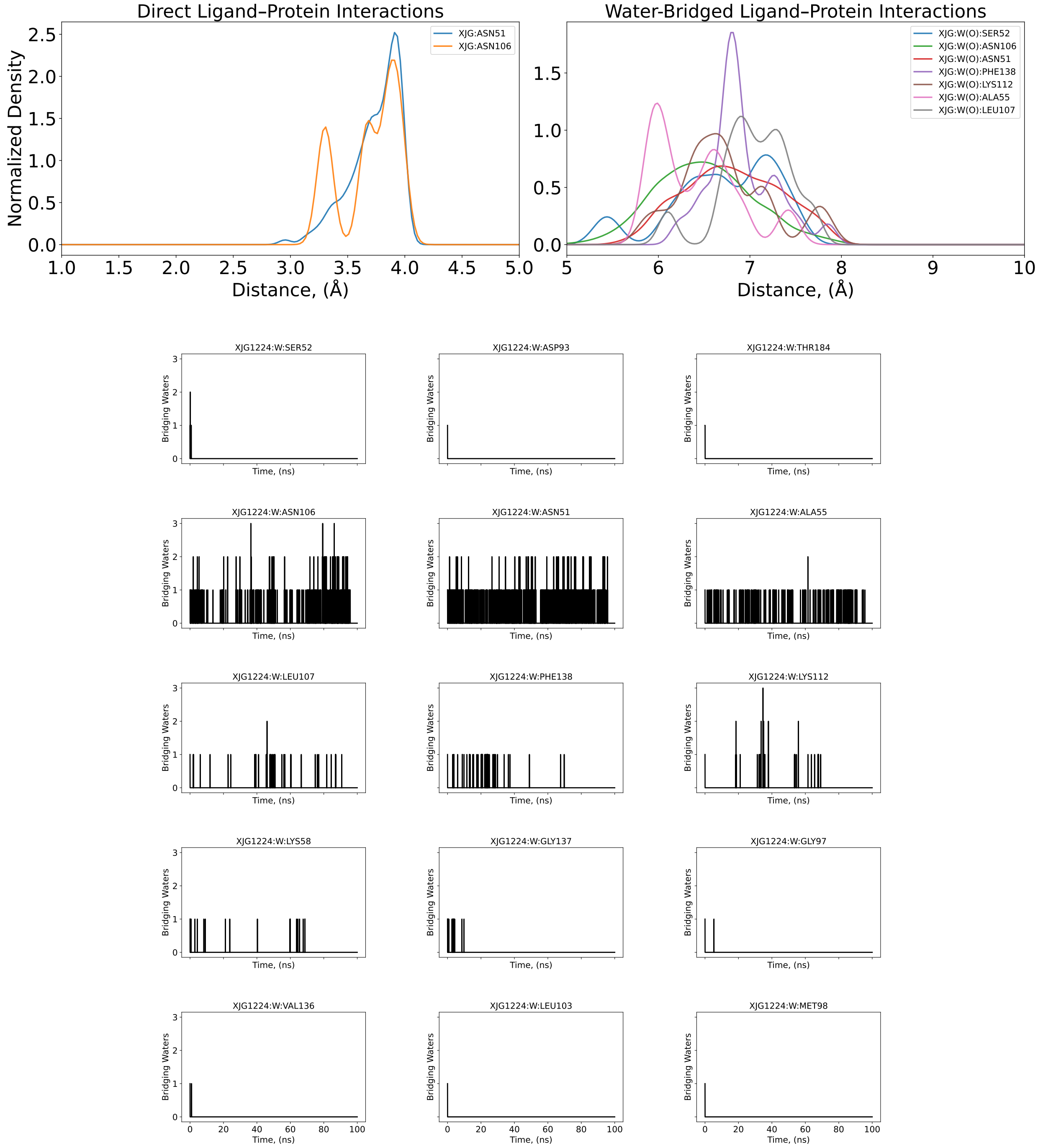


Figure S54. 4-Å–density distribution and individual coordination number of N-bridging water molecules that mediate individual nitrogen protein-ligand atom pairs along 100-ns conventional MD for the heat shock protein HSP90-$\alpha$ complex (PDB: 2XJG).

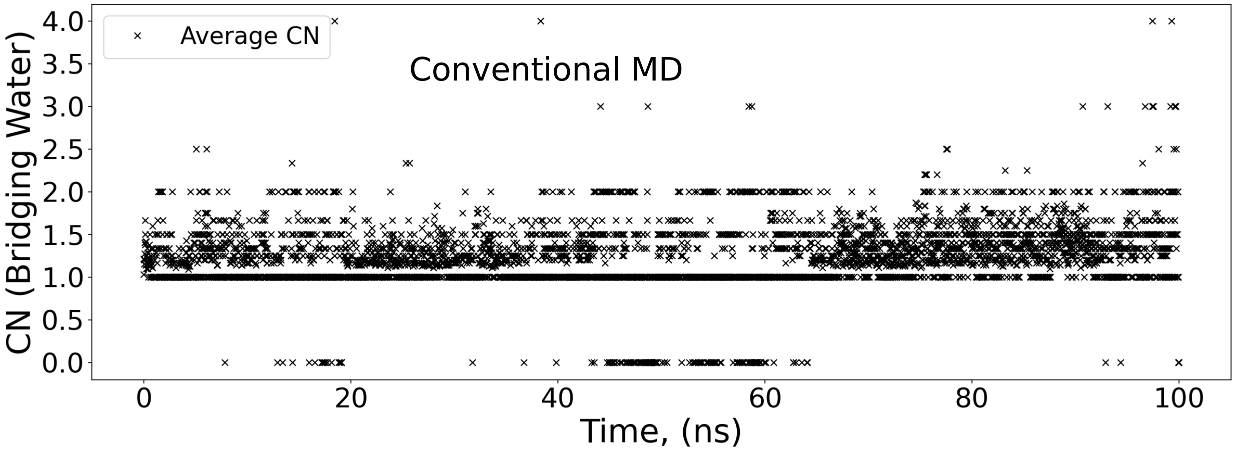


Figure S55. Time-series analysis of the cumulative bridging water coordination number (CN) over the conventional MD. Bridging events were defined by a geometric dual-occupancy criterion, requiring a water molecule to reside simultaneously within 4.0 Å of both a ligand and a protein heavy atom (oxygen or nitrogen) for the heat shock protein HSP90-$\alpha$ complex (PDB: 2XJG).

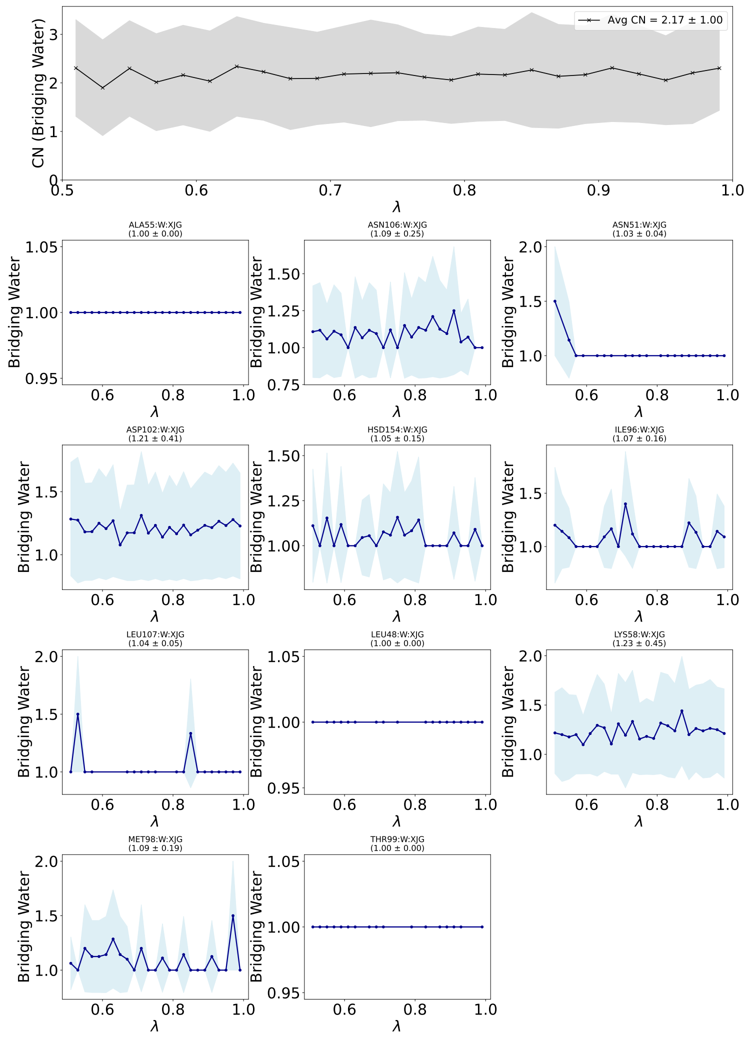


Figure S56. Water-mediated interaction profiles for the bridging motif for electrostatic bound contribution in Lambda-ABF-OPES binding affinity calculations for for the heat shock protein HSP90-$\alpha$ complex (PDB: 2XJG). (Top Panel) Global analysis of the total hydration stoichiometry within the binding pocket. The black trace represents the conditional average count of water molecules participating in bridging interactions across all detected residue pairs. The gray shaded region indicates the standard deviation ($\pm 1\sigma$) across the ensemble, reflecting fluctuations in solvent multiplicity. (Bottom Panels) Site-specific hydration profiles for key anchoring residues. Each panel displays the average number of water molecules bridging the specific residue–ligand pair as a function of the alchemical coordinate $\lambda$. Error bands represent the fluctuation in water count during bridging events.

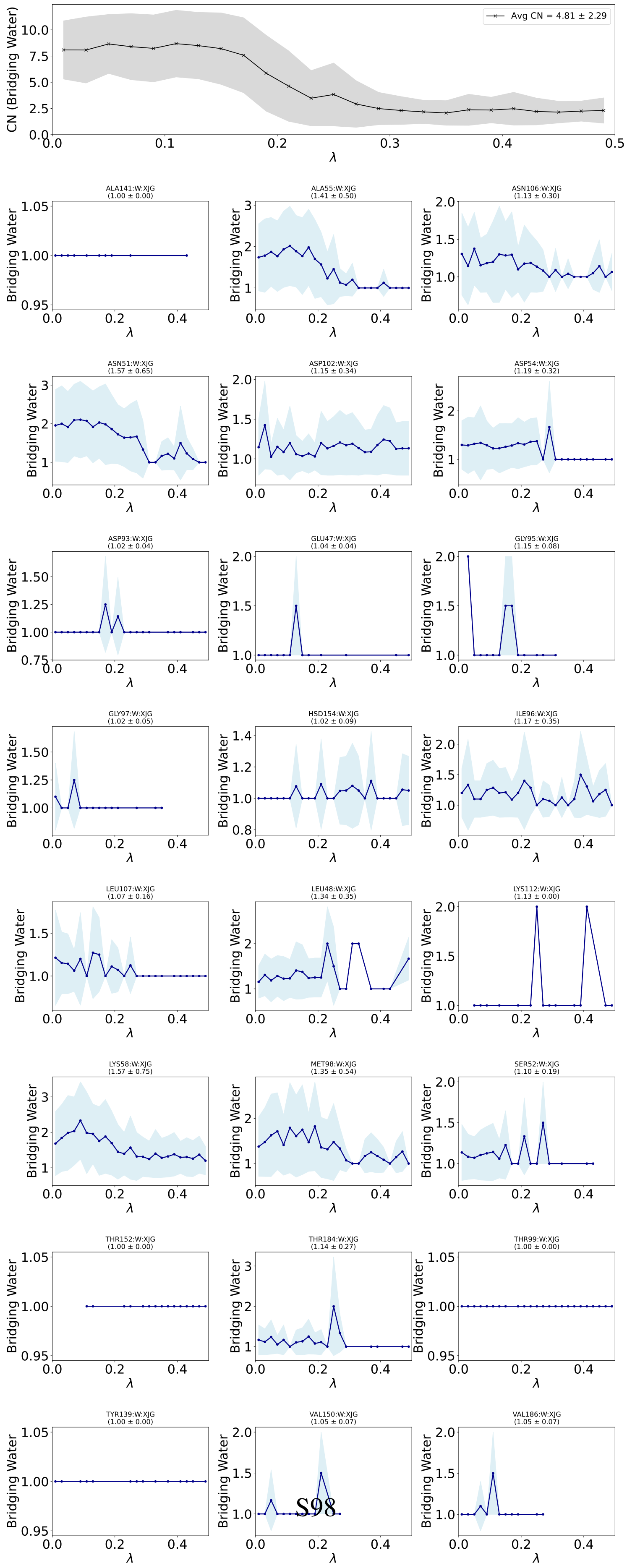
Avg CN = 4.81 ± 2.29
CN (Bridging Water)
λ
Bridging Water
ALA141:W:XJG (1.00 ± 0.00)
ALA55:W:XJG (1.41 ± 0.50)
ASN106:W:XJG (1.13 ± 0.30)
ASN51:W:XJG (1.57 ± 0.65)
ASP102:W:XJG (1.15 ± 0.34)
ASP54:W:XJG (1.19 ± 0.32)
ASP93:W:XJG (1.02 ± 0.04)
GLU47:W:XJG (1.04 ± 0.04)
GLY95:W:XJG (1.15 ± 0.08)
GLY97:W:XJG (1.02 ± 0.05)
HSD154:W:XJG (1.02 ± 0.09)
ILE96:W:XJG (1.17 ± 0.35)
LEU107:W:XJG (1.07 ± 0.16)
LEU48:W:XJG (1.34 ± 0.35)
LYS112:W:XJG (1.13 ± 0.00)
LYS58:W:XJG (1.57 ± 0.75)
MET98:W:XJG (1.35 ± 0.54)
SER52:W:XJG (1.10 ± 0.19)
THR152:W:XJG (1.00 ± 0.00)
THR184:W:XJG (1.14 ± 0.27)
THR99:W:XJG (1.00 ± 0.00)
TYR139:W:XJG (1.00 ± 0.00)
VAL150:W:XJG (1.05 ± 0.07)
VAL186:W:XJG (1.05 ± 0.07)

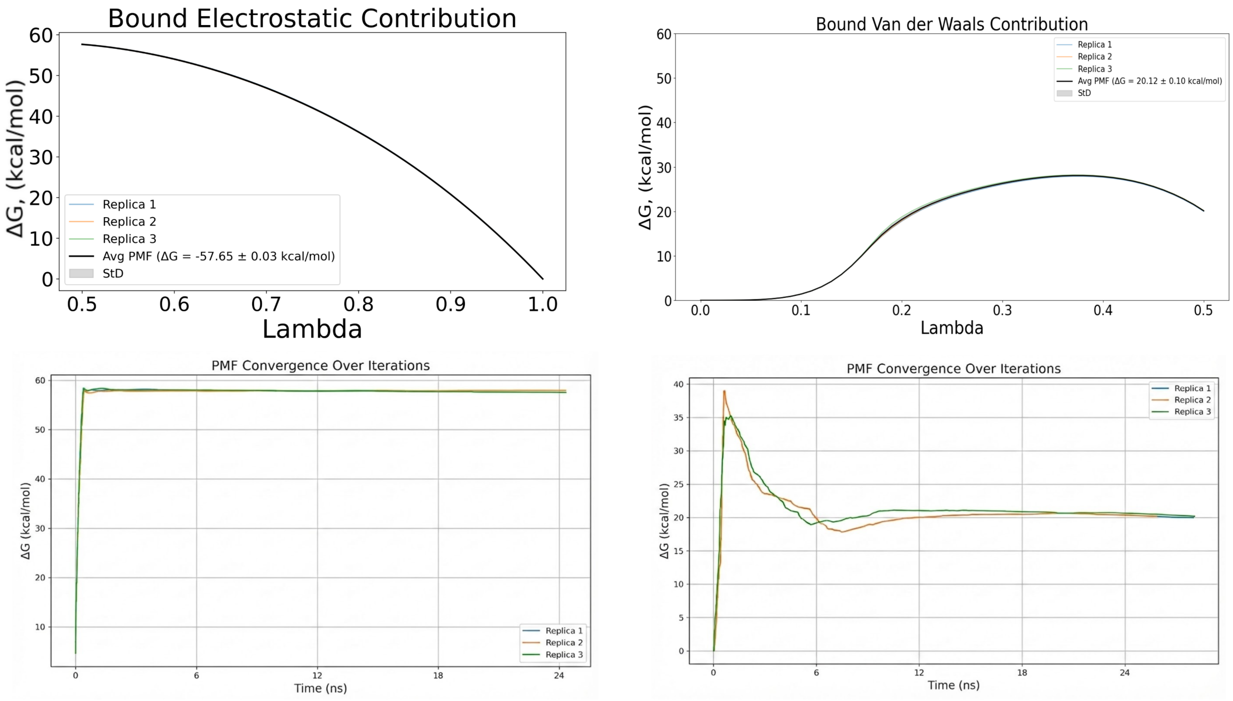


Figure S58. PMF and convergence plots of bound electrostatic and van der Waals energetic contribution for three replicas for the heat shock protein HSP90-$\alpha$ complex (PDB: 2XJG).

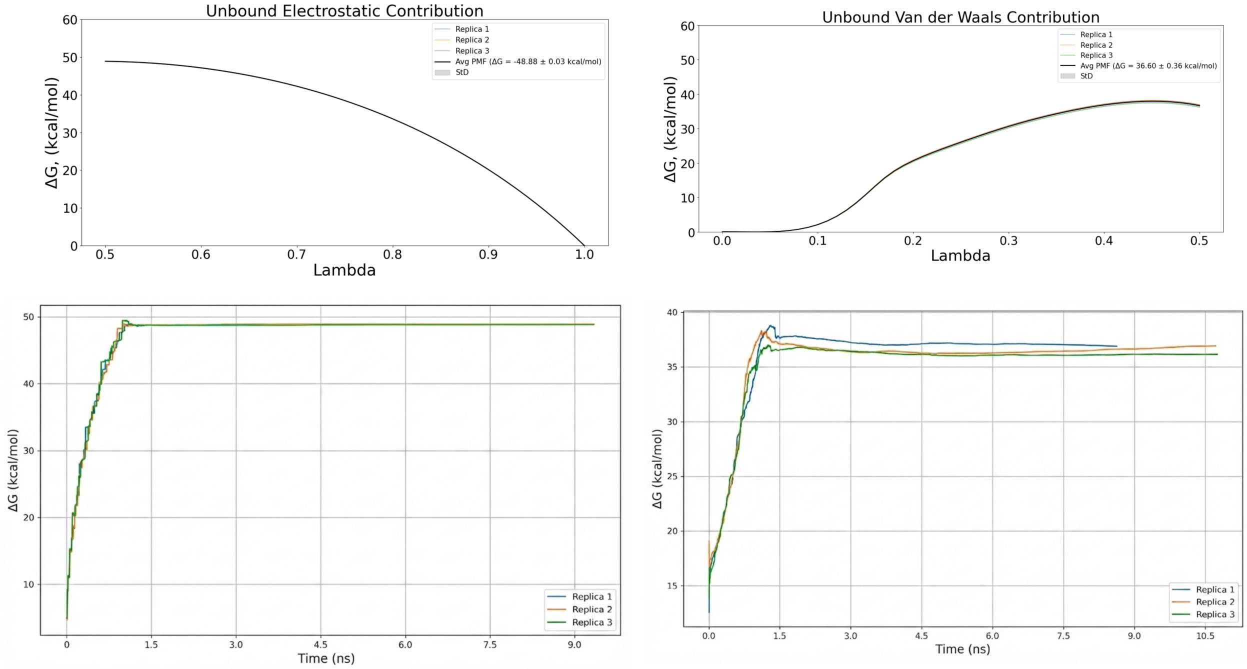


Figure S59. PMF and convergence plots of bound electrostatic and van der Waals energetic contribution for three replicas for the heat shock protein HSP90-$\alpha$ complex (PDB: 2XJG).

#### 2XJG & 2XAB: Binding Affinity Dependence of Protein Modeled Conditions

**Table S10. Comparison of 2XJG and 2XAB under Different Protein Modeled Conditions**

| 2XJG | | | |
|---|---|---|---|
| Protein With Modeled Disordered Region | | Protein Without Modeled Disordered Region | |
| $\Delta G_{VDW}^{site}$ | $\Delta G_{ELEC}^{site}$ | $\Delta G_{VDW}^{site}$ | $\Delta G_{ELEC}^{site}$ |
| 24.5 ± 1.2 | -45.9 ± 0.2 | 20.1 ± 0.1 | - 57.7 ± 0.0 |
| 2XAB | | | |
| 43.0 ± 0.9 | -85.0 ± 0.3 | 19.7 ± 0.9 | -56.9 ± 0.3 |

All the $\Delta G$ contributions are represented in kcal·mol$^{-1}$ units.

### Continuous coordination number analysis of binding-site hydration over Lambda-ABF-OPES Bound Calculations

To quantify ligand-induced rehydration of the binding pocket along the alchemical transformations, we analyzed the coordination of water molecules mediating protein–ligand interactions using a continuous coordination number (CN) formalism. The analysis focuses on electronegative atoms exposed to the binding cavity and involved in water-mediated contacts, identified from conventional equilibrium simulations. All analyses were performed using MDAnalysis, with trajectories aligned to a reference structure based on protein $C_\alpha$ atoms to remove global rotational and translational motion. Water molecules were defined by their oxygen atoms, and distances between water oxygen and corresponding atom of the residue of interest were considered in the hydration analysis.

#### Continuous Coordination Number Definition

The coordination number at a given frame was computed as a distance-weighted sum over all water oxygen atoms surrounding a selected protein atom, using a rational switching function of the form introduced by Ansari et al.[5]:

$$\mathrm{CN} = \sum_{i}^{N_{\text{water}}} \frac{1 - (r_i/r_0)^n}{1 - (r_i/r_0)^m} \tag{24}$$

where $r_i$ is the minimum distance between the protein atom and water oxygen $i$, $r_0 = 4$ Å is the characteristic hydration cutoff, and the exponents $n = 6, m = 12$ control the smooth decay of the switching function. This continuous formulation avoids the discretization artifacts inherent to hard distance cutoffs and ensures that partial hydration contributions are preserved.

#### Replica-Resolved, $\lambda$-Aligned Evaluation

Unlike cutoff-based approaches applied to concatenated trajectories, the continuous CN was evaluated independently at each simulation frame and explicitly aligned with the corresponding alchemical coupling parameter $\lambda$. For each alchemical leg (van der Waals and electrostatic decoupling), CN values were computed across all walkers and replicas and subsequently pooled according to their instantaneous $\lambda$ values.

#### Hydration Probability Landscapes

To visualize the $\lambda$-dependent populations and exchange behavior of site-specific hydration motifs, two-dimensional probability-density maps were constructed from the joint distribution of CN and $\lambda$ using kernel density estimation (KDE). Separate landscapes were generated for the van der Waals and electrostatic decoupling legs, allowing direct comparison of steric- versus electrostatically driven hydration changes. This representation captures both the mean hydration level and the breadth of fluctuations at each alchemical state, providing insight into the sampled persistence, exchange behavior, and structural organization of the hydration motifs of bridging water molecules.

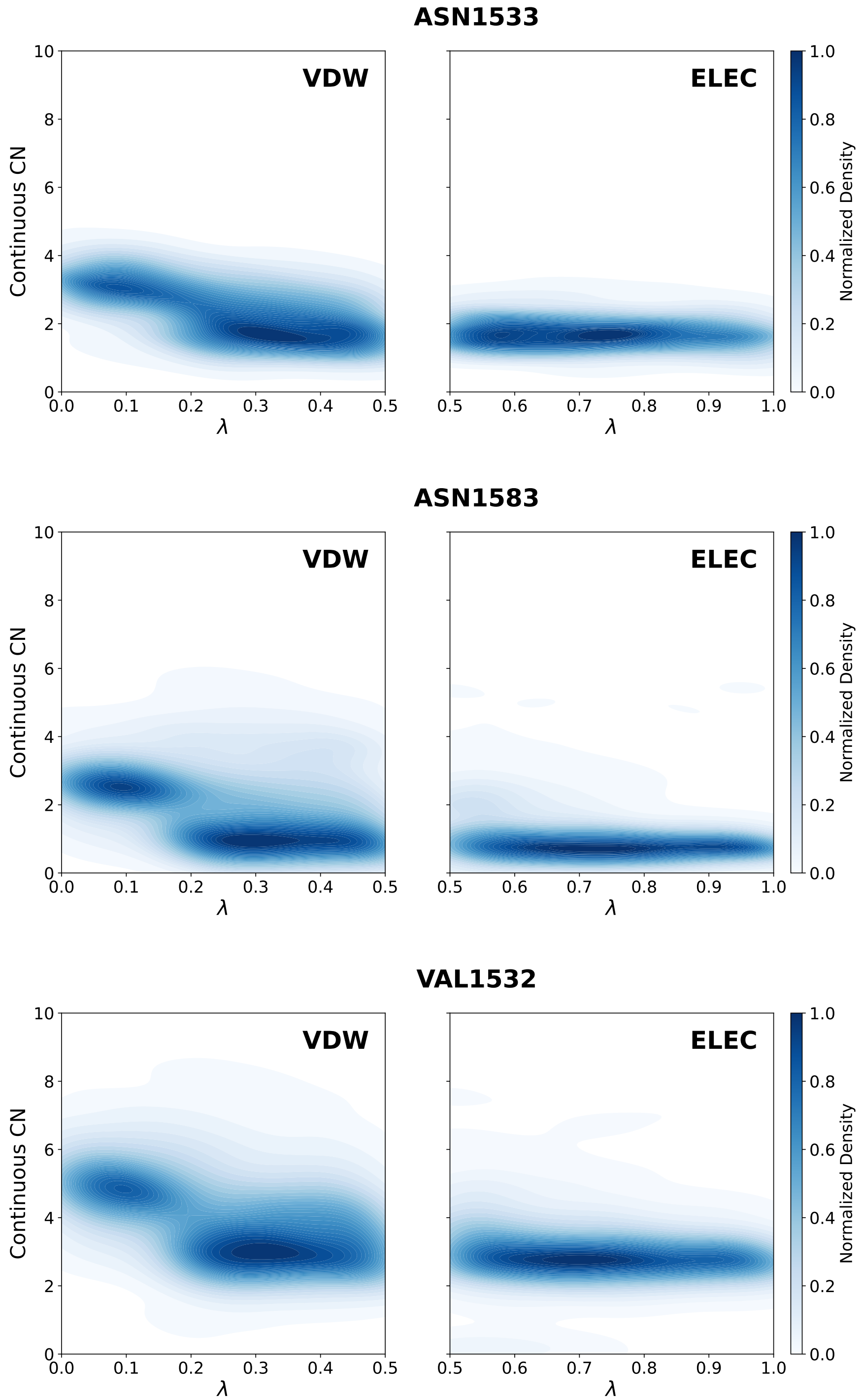


Figure S60. Lambda-dependent hydration profiles of the TAF1(2) binding pocket (PDB id 5I1Q). 2D probability density maps of the continuous CN for nitrogen atom of peptide bond of ASN1533 (top), nitrogen atom of carboxamide of ASN1583 (middle) and oxygen atom of VAL1532 (bottom) as a function of the alchemical coupling parameter ($\lambda$). (Left) The VDW decoupling leg ($\lambda : 0.5 \rightarrow 0.0$). (Right) The electrostatic decoupling leg ($\lambda : 1.0 \rightarrow 0.5$).

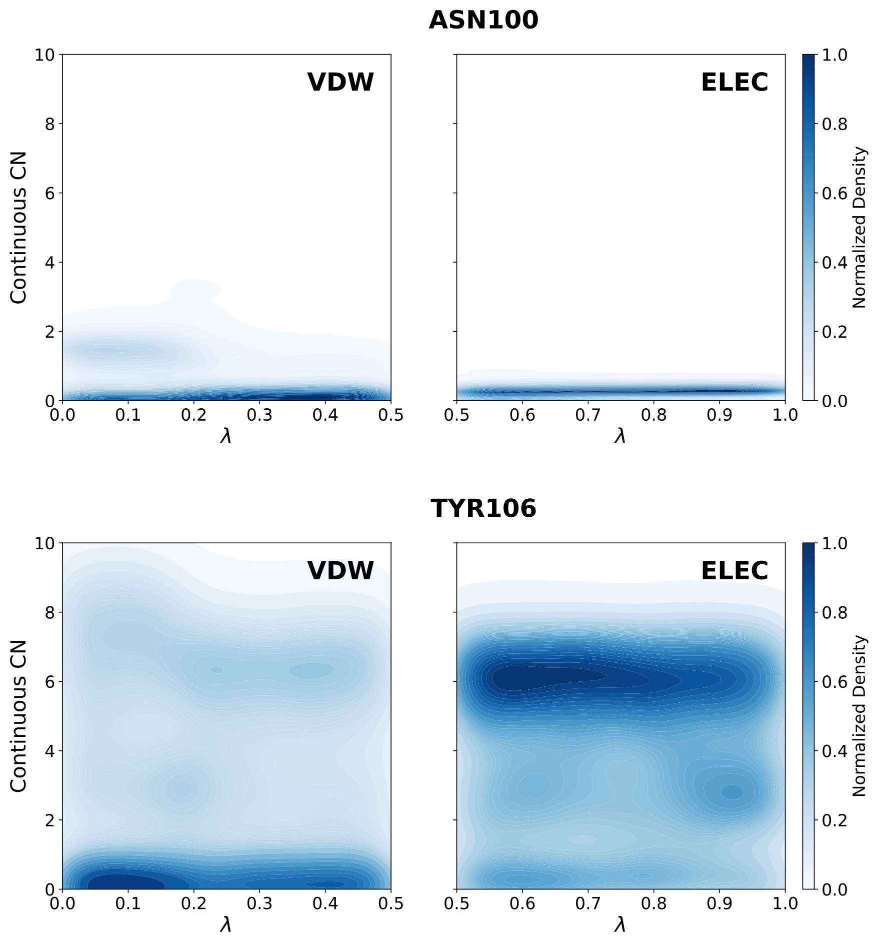


Figure S61. Lambda-dependent hydration profiles of the BRD9 binding pocket (PDB id 5I40). 2D probability density maps of the continuous CN for oxygen atom of ASN100 (top), oxygen atom of hydroxyl group of TYR106 (bottom) as a function of the alchemical coupling parameter ($\lambda$). (Left) The VDW decoupling leg ($\lambda : 0.5 \rightarrow 0.0$). (Right) The electrostatic decoupling leg ($\lambda : 1.0 \rightarrow 0.5$).

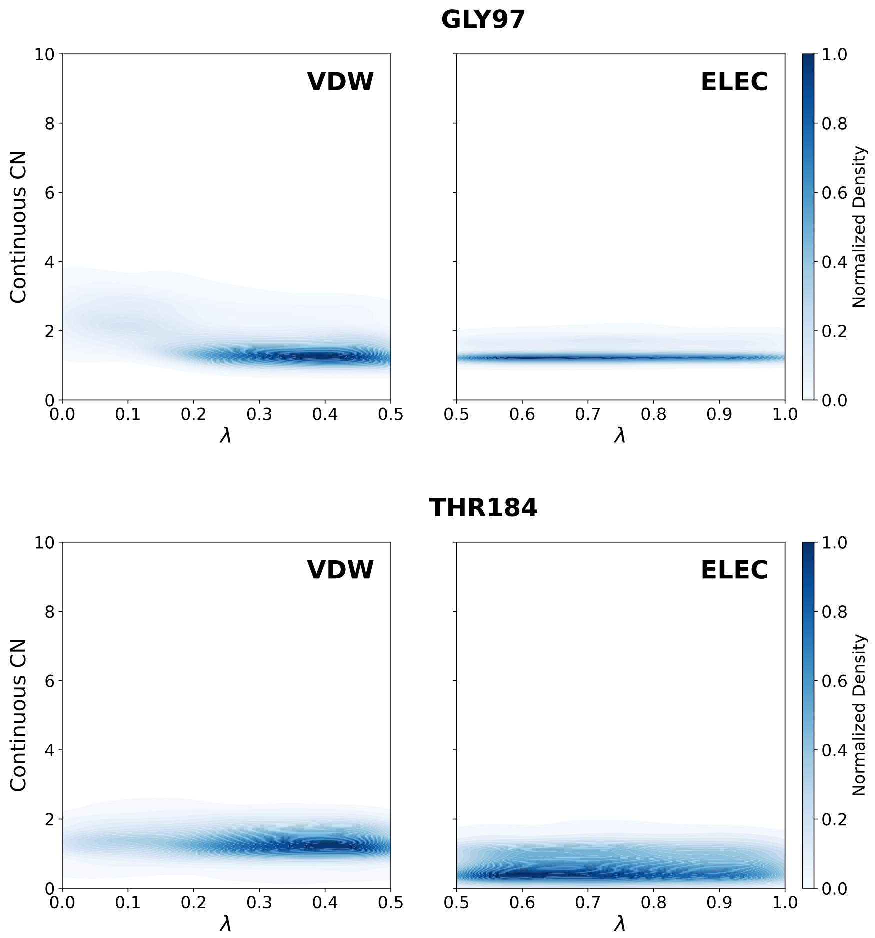


Figure S62. Lambda-dependent hydration profiles of the Hsp90 binding pocket (PDB id 2XAB). 2D probability density maps of the continuous CN for nitrogen atom of GLY97 (top), oxygen atom of THR184 (bottom) as a function of the alchemical coupling parameter ($\lambda$). (Left) The VDW decoupling leg ($\lambda : 0.5 \to 0.0$). (Right) The electrostatic decoupling leg ($\lambda : 1.0 \to 0.5$).

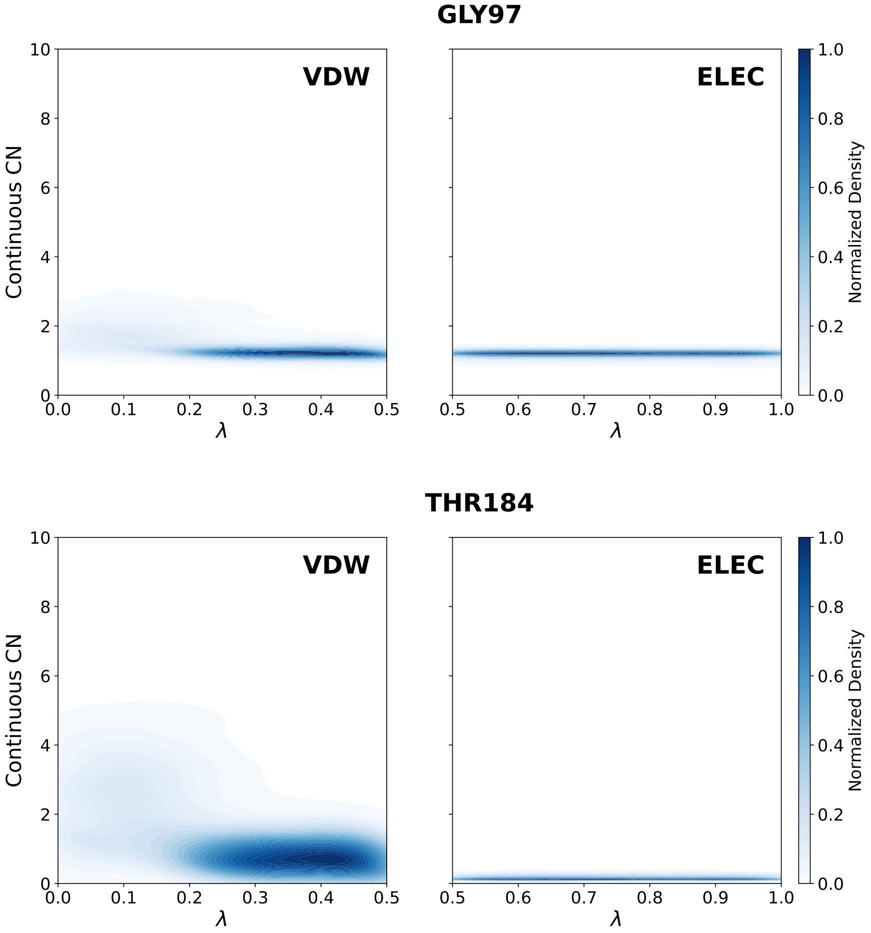


Figure S63. Lambda-dependent hydration profiles of the Hsp90 binding pocket (PDB id 2XJG). 2D probability density maps of the continuous CN for nitrogen atom of GLY97 (top), oxygen atom of THR184 (bottom) as a function of the alchemical coupling parameter ($\lambda$). (Left) The VDW decoupling leg ($\lambda : 0.5 \rightarrow 0.0$). (Right) The electrostatic decoupling leg ($\lambda : 1.0 \rightarrow 0.5$).
